# MondoA mediates transcriptional coordination between the MYC network and the integrated stress response in pancreatic ductal adenocarcinoma

**DOI:** 10.1101/2025.09.03.674106

**Authors:** Erin L. Ramsey, Stephanie Dobersch, Brian Freie, Nan Hyung Hong, Xiaoying Wu, Sita Kugel, Robert N. Eisenman, Patrick A. Carroll

## Abstract

MYC amplification contributes to poor survival and outcome in pancreatic ductal adenocarcinoma (PDAC). Here we show that in PDAC cell lines with amplified MYC, MondoA is required for viability, facilitating proliferation while suppressing apoptosis *in vitro* and *in vivo*. Transcriptional and genomic profiling demonstrates that loss of MondoA leads to altered expression of direct MondoA targets as well as MYC target genes and is accompanied by shifts in genomic occupancy of MYC, MNT, and the MondoA paralog ChREBP. This altered genomic binding by MYC network members is associated with transcriptional perturbation of multiple metabolic and stress pathways, as well as global changes in N6-methyladenosine modification (m^6^A) of mRNA. MondoA inhibition disrupts coordination between MYC network members and the Integrated Stress Response (ISR), resulting in decreased translation of ATF4 mRNA, discordant gene regulation of shared targets of MYC and ATF4 and, ultimately, apoptosis. Re-establishing ATF4 protein expression rescues the diminished viability due to loss of MondoA expression or activity, providing direct evidence of a link between deregulated MYC and the transcriptional machinery of the ISR. Lastly, we find that small-molecule inhibition of MondoA is lethal in a subset of PDAC cell lines, including patient-derived organoids, suggesting that the ability to target MYC via chemical inhibition of MondoA transcriptional activity may have broad efficacy.

**Significance Statement:** This report investigates mechanisms underlying the dependence of MYC-amplified pancreatic cancer cells on the MYC network member MondoA which, as a heterodimer with MLX, is a nutrient-sensing transcription factor. We show this dependency is linked to genomic crosstalk between MYC, components of the proximal MYC network, and the master regulator of the integrated stress response, ATF4. Moreover, we find that small molecule inhibitors of MondoA-MLX transcriptional activity abrogate survival of MYC-amplified PDAC lines and patient derived organoids. The significance of this work relates to its focus on a unique vulnerability intrinsic to MYC, an oncogenic driver associated with a wide range of cancers, which is considered to be “undruggable”.

## Introduction

Metabolic sufficiency during normal development is required for biomass accumulation, genome replication, and subsequent cell division induced by growth factors. These linked mitogenic and biosynthetic pathways are targets of mutation (e.g. PI3K-AKT, RAS, MTOR) and can be activated or inhibited in malignancies. Components of the MYC network (reviewed in [1]) consist of interconnected mitogenic (MYC-MAX) and nutrient-responsive (MondoA/ChREBP-MLX) nodes of heterodimeric transcriptional activators and repressors. These transcriptional regulators, while acting to coordinate normal growth and metabolism during development, are also implicated in the pathological trajectories of the same pathways in diabetes, obesity, and cancer. Mitogenic stimulation of MYC leads to cell cycle progression and concurrent activation of biosynthetic pathways required for cell division. Deregulation of MYC in cancer drives a “selfish” biomass accumulation program coupled with sensitization to metabolic apoptotic stimuli [2]. The MYC network exhibits complex interactions and cell-type specific functions, including both context-dependent oncogenic or tumor suppressive roles for individual members. Alterations to the network have been found to elicit new dependencies and synthetic lethal interactions between the MAX and MLX arms of the network. Examples include MYC/MYCN-amplified tumors requiring MondoA-MLX [3], and MAX-null tumors being sensitive to chemical inhibition of MondoA [4].

Thus, the nutrient-sensing MLX arm of the network is responsive to alterations in the mitogen-responsive MAX arm.

Cellular metabolic state is monitored by a variety of nutrient-sensors that provide feedback to activate or inhibit both mitogenic and metabolic pathways. This includes Mondo proteins (MondoA and ChREBP, reviewed in[5]), which respond to flux through the major glucose and lipid metabolic pathways (e.g. glucose induces Mondo-dependent TXNIP, which negatively regulates glucose uptake). Prolonged metabolic stress (either deficiency or overabundance) can lead to activation of the Integrated Stress Response (ISR) and its transcriptional effector, ATF4, to either re-balance cellular metabolism or, alternatively, undergo apoptosis.(reviewed in [6]).

The ISR integrates disparate signals (e.g. glucose, lipid, amino acid, ROS levels) with transcriptional programs that can be remedial (pro-survival) or terminal (pro-apoptotic), dependent upon duration and intensity of the signal. A fundamental output of the ISR is inhibition of global protein synthesis coupled with a switch to selective translation of uORF-containing transcripts, such as ATF4 (reviewed in[7]). Deregulated oncogenes, such as MYC or KRAS, are known to induce the ISR through multiple pathways, such as metabolic stress leading to ROS production, increased translation leading to ER stress (ERS) via protein misfolding, and the Unfolded Protein Response (UPR). MYC requires components of the UPR/ISR (reviewed in [8]) and cooperatively shares targets and promoter co-occupancy with ATF4 for critical transcriptional crosstalk between MYC network and the ISR [9]. The MondoA/ChREBP-MLX target TXNIP is a component of this apoptotic cascade, as deletion of either ChREBP or TXNIP prevents glucotoxicity-induced beta-islet cell death in diabetes [10, 11]. Taken together, the literature suggests crosstalk between multiple components of the MYC network and the ISR is critical for cell function.

Pancreatic cancer progression, in transitioning from pancreatitis to pancreatic intraepithelial neoplasia (PanIN) and ultimately to pancreatic ductal adenocarcinoma (PDAC), also involves MYC-driven metabolism, RAS-induced senescence, and ISR components (see review of UPR/ISR in PDAC progression [12]). MYC copy number gain has been shown to correlate with decreased PDAC patient survival[13]) but can also activate or intensify putative targetable vulnerabilities (e.g. hyper-transcription, metabolic and other downstream effectors such as hyper-translation, proteostasis[14] and ISR pathways). PDAC has an abysmal outcome and survival rate, and actionable targets are required to develop effective therapeutics. We hypothesized that MYC network interactions could be critically important in sustaining oncogenesis in MYC amplified PDAC. We therefore sought to determine the functional interactions between MYC and MondoA in this malignancy, as well as explore potential for targeting such interactions.

## Results

### MondoA regulates viability in MYC-amplified PDAC lines

To determine whether pancreatic cancer cells require MondoA for growth and survival, we utilized siRNA against MondoA and assessed relative growth rates in 4 PDAC lines compared to non-targeting siControl. As MYC genomic amplification correlates with poor patient outcome and survival [13], this cell line panel included two lines with MYC amplification (PSN-1 and DAN-G) and two without (PANC 08.13 and SU.86.86). Following MondoA knockdown, we observed a significant inhibition of growth only in MYC-amplified cells compared to non-MYC-amplified cells (Fig. 1A-B). As monitored by western blot, the levels of MondoA protein knockdown were similar in the four lines while MYC levels were unaffected (Fig. 1C). To directly test whether higher expression of MYC potentiates growth inhibition upon MondoA knockdown, we engineered a dox-inducible MYC system in the non-amplified SU.86.86 cell line and monitored growth in the presence and absence of MondoA. As shown in Figure 1D-E, ectopic MYC increases cell number in the SU.86.86 cells, and this increase is abrogated upon MondoA knockdown. This suggests that deregulated MYC sensitizes these cells to loss of MondoA. If this is the case, then decreasing MYC levels should attenuate cell death upon siMondoA treatment. As shown in Figure 1F, baseline caspase 3/7 activity is diminished significantly by siMYC treatment in all 4 lines, while it is increased significantly by siMondoA treatment in only the PSN-1 and DAN-G cells. This increase is mitigated by the combination of siMYC with siMondoA co-treatment. Therefore, MondoA loss promotes MYC-dependent apoptosis in MYC-amplified PDAC lines.

**Figure 1.**
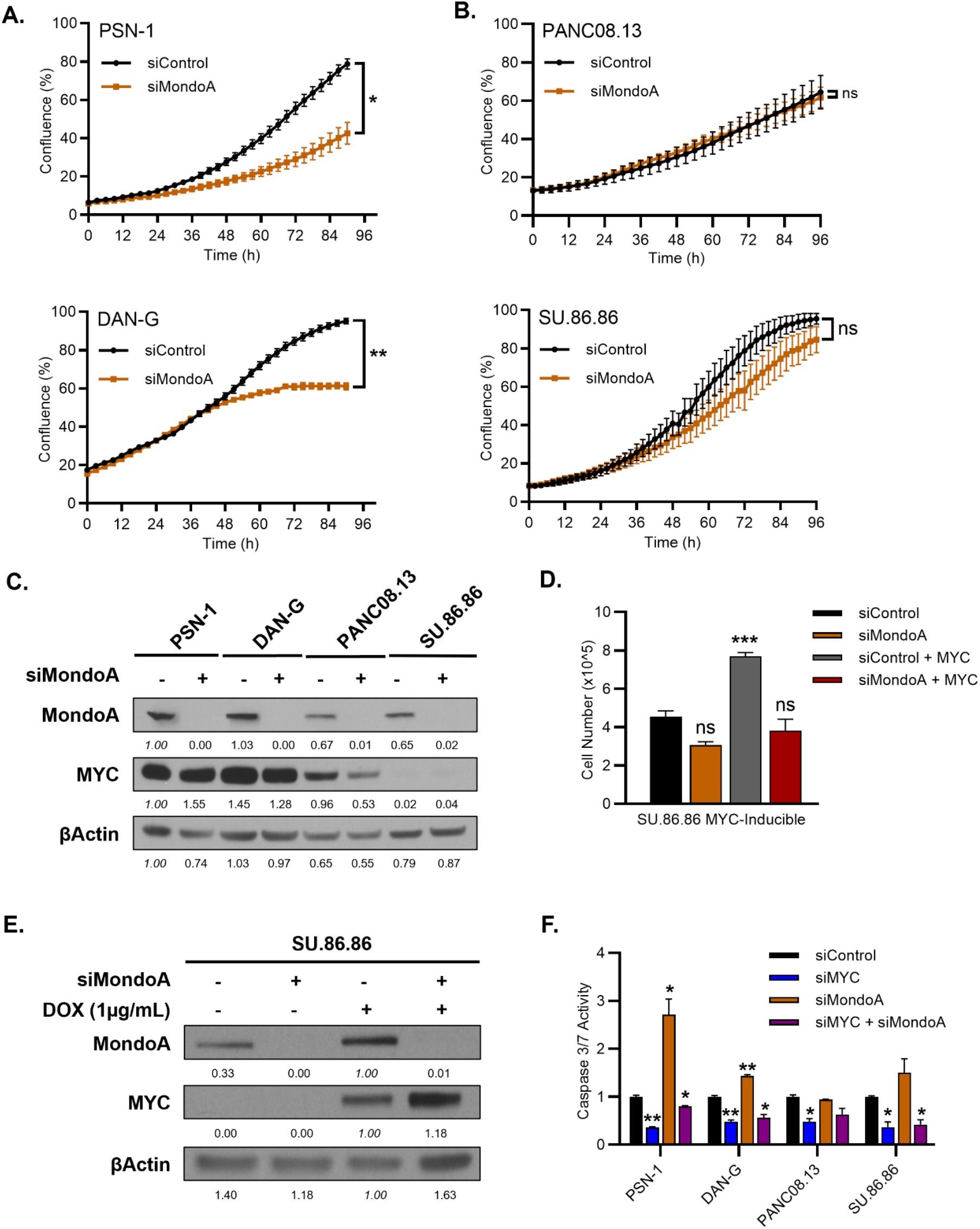
Functional Interactions between MYC and MondoA in PDAC cell lines. (A) Incucyte growth curves of MYC High PSN-1 and DAN-G (n=4) or (B) MYC Low PANC08.13 and SU.86.86 (n=3) PDAC cell lines treated with siControl (Black) or siMondoA (Orange) over 96 hours. Data expressed as plate confluency and significant differences (two-tailed t-test) as compared to final confluence of corresponding siControl. (C) Western blot for MondoA, MYC, and βActin in a panel of PDAC lines (PSN-1, DAN-G, PANC08.13, SU.86.86) treated with either siControl or siMondoA for 72 hours. Relative abundance of protein expressed in relation to siControl treated PSN-1 cells and normalized to relative abundance of βActin. (D) SU.86.86 cells with a DOX-Inducible MYC vector were treated with either siControl or siMondoA and presence or absence of DOX (1µg/ml) for 72 hours. Data expressed as mean cell count (x10^5) and significant differences (n=3, one-way ANOVA) as compared to siControl without DOX. (E) Western blot for MondoA, MYC, and βActin corresponding to panel 1D. Relative abundance of protein expressed in relation to siControl cells with DOX treatment and normalized to βActin. (F) Caspase-Glo 3/7 system was used to assess apoptosis in the indicated panel of PDAC lines treated with either siControl or siMYC and siMondoA alone or in combination for 72 hours. Data expressed as mean luminescence and normalized within each line to siControl cells. Significant differences (n=4, one-way ANOVA) as compared to siControl within each cell line. All data expressed as mean±SEM and p-values expressed as ns= not significant; * p≤0.05; ** p≤0.005, *** p≤0.0005, **** p≤0.0001.

### MondoA promotes clonogenic and xenograft outgrowth of PSN-1 cells

To extend our findings on the relationship between MYC levels in PDAC cells and MondoA inhibition, we assessed the growth of cells as foci in a clonogenic assay. As shown in Figure 2A, siRNA against MondoA impairs clonogenic survival of the MYC-amplified PSN-1 cells, demonstrating a significant decrease in colony formation frequency upon plating at a range of low densities compared to siControl cells. To test the effect of stable knockdown of MondoA, we utilized shRNA expression via a dox-inducible viral vector and compared the effect of shControl versus shMondoA in the presence and absence of doxycycline. (Fig. S1A). This inducible system recapitulates the phenotype of transient siRNA knockdown, as PSN-1 cells harboring the shMondoA construct were impaired for clonogenic outgrowth only in the presence of doxycycline when seeded at an identical cell number (Fig. 2B).

**Figure 2.**
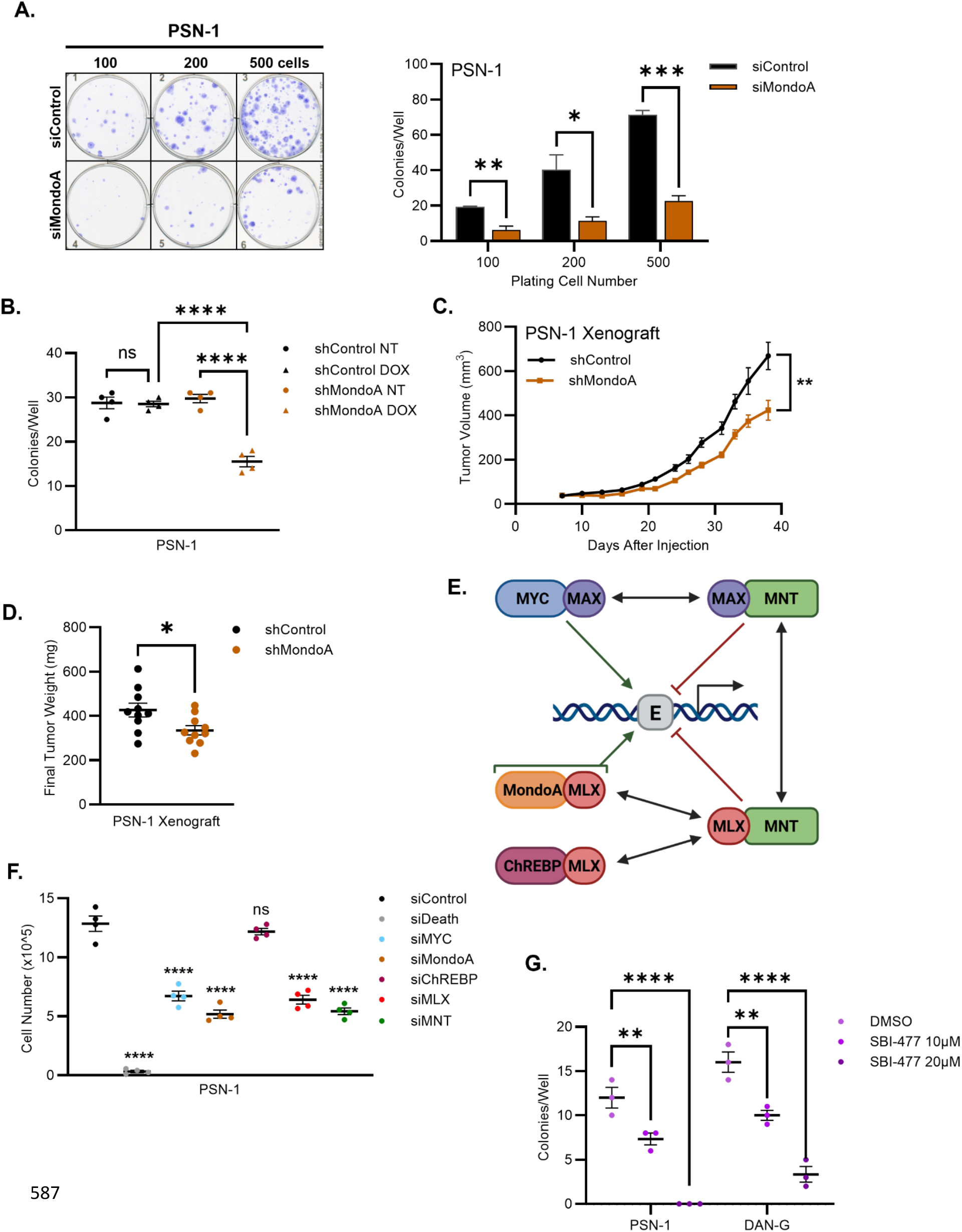
Effects of MondoA-MLX inhibition on MYC-driven PDAC *in vitro* and *in vivo*. (A) Clonogenicity assay of PSN-1 cells seeded at the indicated cells per well and treated with either siControl or siMondoA. Representative plate stained with crystal violet after 12 days, colonies per well plotted. Significant differences (n=3, two-tailed t-test) as compared to siControl within each plating number. (B) Clonogenicity assay of PSN-1 cells with DOX-inducible shMondoA or a control shRNA treated with or without DOX and seeded at 100 cells per well. Colonies per well after 12 days plotted, with significant differences (n=4, one-way ANOVA) within each treatment. (C) Mean tumor volume over time for PSN-1 subcutaneous xenograft with either shControl (left flank) or shMondoA (right flank). Significant differences (n=10, two-tailed t-test) expressed between vectors. (D) Final tumor weight for the 10 subcutaneous xenografts of PSN-1 cells at 38 days post injection. Significant differences (two-tailed t-test) expressed between shRNAs. (E) Simplified schematic of the MYC-network of related bHLHZ transcription factors. (F) Viability expressed as mean cell number for PSN-1 cells following an siRNA screen of MYC-network members for 72 hours. Significant differences (n=4, one-way ANOVA) as compared to siControl cells. (G) Clonogenicity assay of PSN-1 or DAN-G PDAC cells seeded at 50 cells and treated with 0 (DMSO), 10, or 20µM of SBI-477 for 12 days. Mean colonies per well denoted, significant differences (n=3, one-way ANOVA) as compared to DMSO treatment within each cell line. All data expressed as mean±SEM and p-values expressed as ns= not significant; * p≤0.05; ** p≤0.005, *** p≤0.0005, **** p≤0.0001.

We then used this system to probe the role of MondoA for *in vivo* xenograft growth. PSN-1 cells expressing either shControl (left flank) or shMondoA (right flank) were subcutaneously injected into the same mouse for pair-wise comparison. These mice were then provided with 200g/L doxycycline-containing drinking water, and tumor growth and volume were monitored over time. As shown in Figure 2C-D, *in vivo* targeting of MondoA via shRNA significantly decreased both tumor volume and final tumor weight compared to shRNA control (representative mouse and tumors images in Figure S1B-C). Therefore, MondoA is a viable target in MYC-amplified PDAC for blocking growth both *in vitro* and *in vivo*.

### Multiple MYC network members regulate PDAC survival

In order to define the role of MondoA interactors in the PSN-1 cells and directly test any requirement for distinct MLX-containing heterodimers (Network composition in Fig. 2E), we employed siRNA knockdown to determine their potential role in PSN-1 cell survival. Knockdown of MYC, MondoA, MLX, or MNT, but not ChREBP, attenuates viability in PSN-1 cells (Fig. 2F). This suggests that multiple components of the MYC network can support viability. The dependency upon MondoA, MLX, and MNT is also observed in other tumor types, such as B cell lymphoma cell lines[3], murine lymphoma models [15], male germ cell tumor lines [16], and a subset of neuroendocrine tumors [4]. Consistent with these results, we also find that SBI-477, a small-molecule inhibitor of MondoA shown to block MondoA-MLX nuclear translocation [17], significantly attenuates clonogenic growth of both MYC-amplified PSN-1 and DAN-G cells (Fig. 2G), suggesting that chemical targeting of MondoA-MLX transcriptional activity in PDAC could provide a viable therapeutic option. We also observe similar results with SBI-993, a more bioactive derivative of SBI-477 that has been used to target liver cancer models[18](Fig. S1D), further supporting the therapeutic potential of MondoA.

### MondoA loss leads to altered transcription and genomic occupancy of MYC network members

To identify the transcriptional program(s) maintained by MondoA in MYC-amplified PDAC, we performed RNA-seq on PSN-1 cells in which MondoA transcriptional activity was impaired by either siRNA knockdown of MondoA (siMondoA), siRNA knockdown of the MondoA heterodimerization partner MLX (siMLX), or chemical inhibition of MondoA-MLX via SBI-477 treatment. Each of these treatments were compared to either non-targeting siRNA (siControl) or the vehicle control for SBI-477 (DMSO). We identified significantly up and down-regulated differentially expressed genes (denoted as DEGs, adjusted p-value < 0.05) in cells treated with either siMondoA (1640 up, 1489 down), siMLX (655 up, 806 down) or SBI-477 (1547 up, 1394 down) (Fig. 3A-B, Fig. S2A-C, Supplemental Tables 1-2).

**Figure 3.**
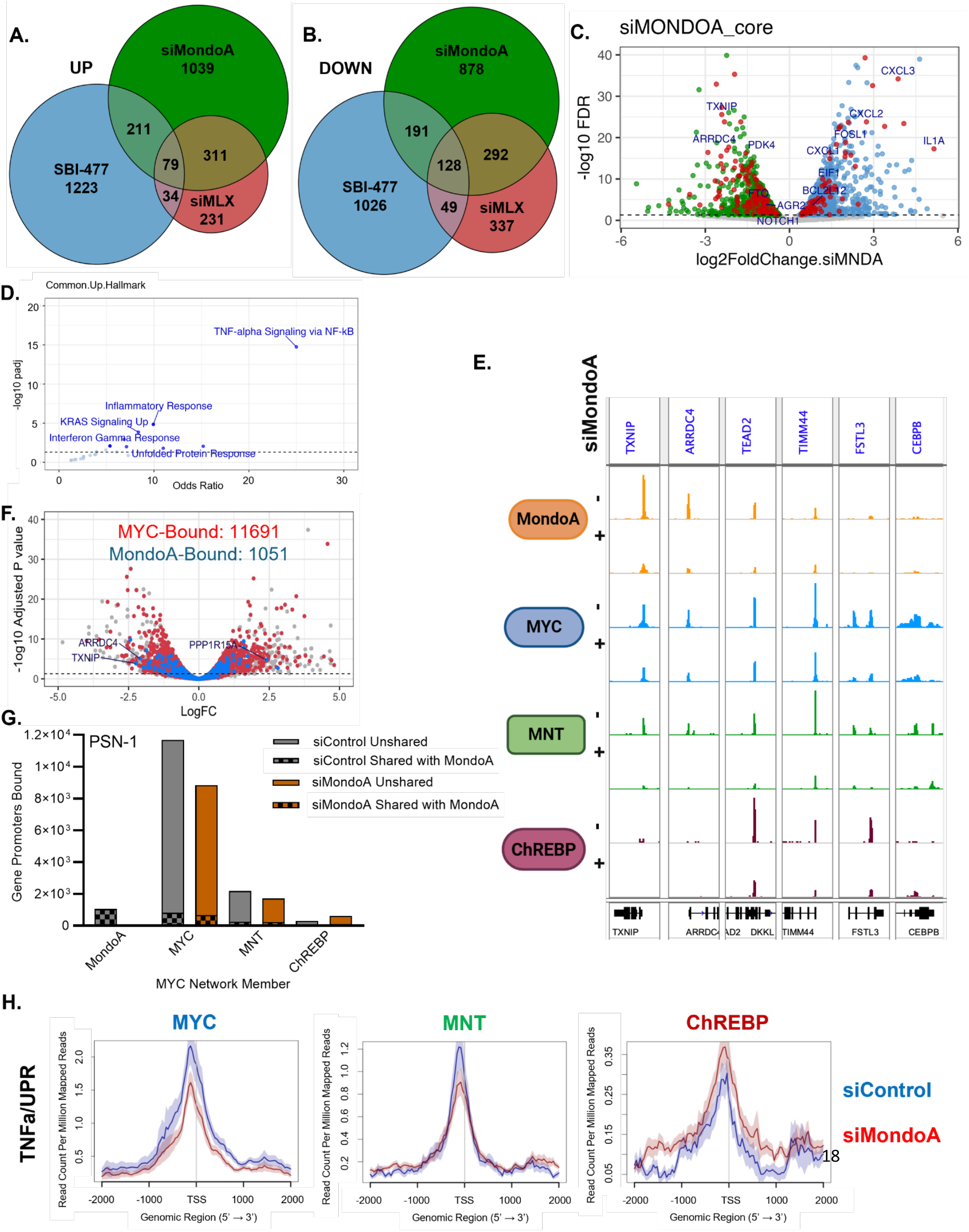
MondoA and MYC network genomic engagement and transcriptional output in PDAC. (A) Venn diagram of differentially expressed genes (DEGs) up-regulated or (B) down-regulated after 72 hours of siMondoA, siMLX, or SBI-477 (20µM) treatment compared to siControl or DMSO. (C) Volcano plot of DEGs after siMondoA treatment. Up-regulated genes in blue, and down-regulated genes in green, with shared DEGs across all three conditions (siMondoA, siMLX, SBI-477, denoted as ‘core’ gene-list) indicated in red. (D) Hallmark pathway analysis of core DEGs up-regulated in ‘core’ gene-list. (E) BigWig tracks for selected genes that lose MondoA and/or exhibit altered MYC network binding with siMondoA treatment. MondoA binding in orange, MYC binding in blue, MNT binding in green, ChREBP binding in maroon. (F) Volcano plot of DEGs after siMondoA treatment with genes bound at the promoter by MYC denoted in red and genes bound by MondoA in blue. Genes labeled are co-bound by both MYC and MondoA. (G) Total count of promoter bound genes for MondoA, MYC, MNT, and ChREBP with siControl (gray) or siMondoA (orange) treatment. Solid bars denote unique targets, checker box denotes shared gene target with MondoA. (H) MYC, MNT, and ChREBP binding over TNFα/UPR target promoters, siControl in blue, siMondoA in red.

We identified sets of “core” up- and down-regulated genes in response to MondoA-MLX inhibition that were changed in all three datasets (Fig. 3A-C, Fig. S3A-B, Supplemental Tables 1-2). Importantly, these core signatures correlate with *MLXIP*, encoding MondoA, within the TCGA PAAD dataset with the downregulated core positively correlating with MondoA (R=0.42, p=6.2e-08) and the upregulated core negatively correlating with MondoA (R=-0.29, p=2.6e-4) (Fig. S3C). The 79 up-regulated “core” DEGs were enriched for inflammatory pathways, including TNFα signaling and the unfolded protein response (UPR) (Fig. 3D), while the 128 down-regulated “core” DEGs were enriched for the DNA damage response and interferon signaling (Fig. S3D). We previously observed similar gene enrichment affecting metabolic genes and stress effectors upon loss of either MondoA or MLX in tumor cell lines [3] or male germ cells [16], and the UPR was targeted by SBI-477 in MAX-null neuroendocrine tumor cells [4]. Among the differentially expressed genes are transcription factors (CEBP, NFκB, FOS) and genes that regulate metabolism (PFAS, FASN, SREBF1), proteostasis (ATF4, PPP1R15A, encoding GADD34), and apoptosis (CDKN1A encoding p21, GADD45A, H2AX) (Figure S3E, Supplemental Tables 1-2). Many of these components of the MondoA-regulated program are critical determinants of homeostasis and cell fate and are also important for MYC-dependent tumorigenesis.

To determine direct versus indirect transcriptional effects of MondoA perturbation upon MYC network members in PSN-1 cells, we investigated the genomic occupancy of MondoA, MYC, MNT, and MondoA paralog, ChREBP, in the presence (siControl) and absence of MondoA (siMondoA) by CUT&RUN. In PSN-1 cells, MondoA is associated with 1051 gene promoters (Fig. 3E-F). MondoA occupancy was detected at previously identified target promoters, such as *TXNIP* and *ARRDC4*, as well as newly identified targets, including *TEAD2* and *TIMM44* (Fig. 3E). Upon siMondoA treatment, its binding was diminished at these gene promoters as expected. Genes identified as both up- and down-regulated in our RNA-seq analysis (including decreased *TXNIP* expression and increased *GADD34* expression) were occupied by MondoA, supporting a dual role for MondoA as both transcriptional activator and repressor.

Interestingly, genes whose promoters were occupied by MYC were likely to be differentially expressed upon MondoA depletion (KS stat 10e-14, comparing RNA-seq data from untreated vs MondoA depleted cells), consistent with the dependence of these cells on MYC-driven transcription. This occurred even as only a small subset of MYC bound genes were occupied by MondoA (1051 of 11691 genes) in these cells (Figure 3F). Thus, MondoA appears to act as a substantial, albeit indirect, effector in the MYC-driven transcriptional program.

We have previously shown that altered expression of individual MYC network members can shift the genomic occupancy of other members [4, 16]. Since knockdown of MondoA alters stress responsive targets of the UPR and TNFα-NFκB pathway, we plotted MYC, MNT and ChREBP genomic coverage in the presence (siControl) and absence of MondoA (siMondoA) over the promoters of genes comprising these pathways. siMondoA resulted in decreased MYC binding (∼25%) at MYC target loci and TNFα/UPR pathway genes compared to control cells (Fig. 3G-H). We also observed a decrease (∼20%) in total MNT binding over these same genes (Fig 3G-H). By contrast, ChREBP exhibited a gain (∼two-fold) in occupation of these gene promoters upon MondoA loss (Fig. 3G-H). This included increased binding at the *CEBPB* promoter, which is in both the TNFα and UPR Hallmark categories (Fig. 3E). Altered binding by MYC, MNT, and ChREBP occurred at both up- and down-regulated gene promoters. This suggests that perturbation of MondoA leads to shifts in genome occupancy by multiple network factors, thereby contributing to dysregulation of both unique and shared targets, including the UPR.

### The UPR is involved in the response to MondoA inhibition

Because induction of the UPR has been shown to inhibit global protein synthesis, we surveyed translation rates in relation to MondoA and MYC in our MYC-low versus MYC-amplified PDAC lines. We used puromycin incorporation to measure possible alterations in protein synthesis upon MondoA knockdown. As expected, the highest levels of puromycin incorporation were observed in the two MYC-amplified cell lines, and MondoA loss decreased puromycin incorporation in all 4 lines by ∼50-65% (Fig. 4A). To examine sensitivity to UPR-induced inhibition of protein synthesis, we tested the dose-responsiveness of these lines to a UPR-inducing drug, Tunicamycin (Tm). We found that the MYC-amplified lines exhibited increased sensitivity to Tm treatment relative to the MYC-low lines (Fig. 4B), suggesting that high MYC expression poises the UPR for activation. This is likely due to increased proteostatic stress from enhanced biosynthetic demand driven by MYC [2].

**Figure 4.**
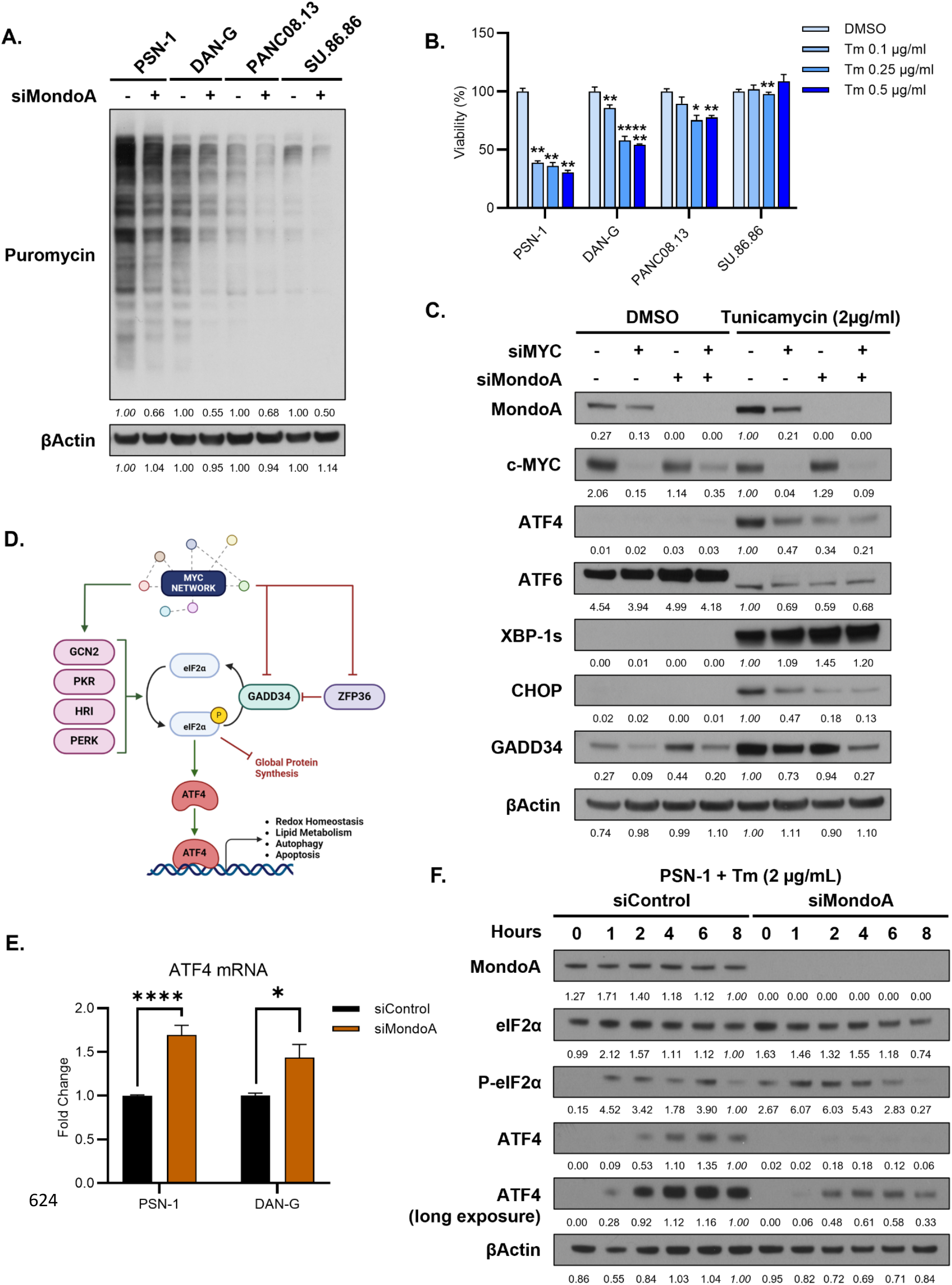
Crosstalk between MondoA, MYC and proteostasis via the UPR/ISR. (A) Puromycin incorporation assay for PDAC cell line panel treated with siControl or siMondoA for 48 hours. Relative abundance of puromycin incorporated into newly synthesized proteins as detected by anti-puromycin western blot shown in relation to siControl treated cells and normalized to βActin within each cell line. (B) Cell viability expressed as percent of DMSO treatment after 48 hours for a panel of PDAC cell lines treated with indicated concentrations of Tunicamycin. Significant differences (n=3, one-way ANOVA) expressed as compared to DMSO treatment within lines. (C) Western blot surveying the three primary effector arms of the unfolded protein response in PSN-1 cells with siMondoA, siMYC, or combined, under baseline conditions or with UPR activation with Tunicamycin for 6 hours. Relative protein abundance expressed in relation to tunicamycin treated siControl cells and normalized to βActin. (D) Schematic of MYC Network interactions, as determined by promoter occupancy, with the Integrated Stress Response and ATF4. Green lines indicate activation, red lines repression. (E) ATF4 mRNA levels by RT-qPCR with and without siMondoA in MYC amplified PDAC lines PSN-1 (n=11) and DAN-G (n=7). Data expressed as mRNA fold-change, significant differences (two-tailed t-test) as compared to siControl within each line. (F) Time course western blot for ATF4 and its upstream activators under activation by Tunicamycin (2ug/mL) with and without MondoA in PSN-1 cells. Relative protein abundance expressed as compared to maximal tunicamycin-induction at 8 hours in siControl treated cells and normalized to βActin. All data expressed as mean±SEM and p-values expressed as ns= not significant; * p≤0.05; ** p≤0.005, *** p≤0.0005, **** p≤0.0001.

To further examine the MYC dependence on MondoA and the UPR, we assessed the three major UPR pathways in PSN-1 cells subsequent to Tm treatment and knockdown of MYC and MondoA, alone or in combination. While Tm induced all three expected downstream UPR transcriptional effectors (ATF4 protein translation, ATF6 cleavage, and XBP-1 splicing, reviewed in [8] and [12]), ATF4 alone required the combined presence of MYC and MondoA for maximal expression, even with Tm induction (Fig. 4C). This combinatorial requirement was also observed for the ATF4-downstream target CHOP, whereas GADD34, which is also an ATF4 target, was repressed directly by MondoA as seen between the first and third lanes (Fig. 4C). This data supports a coordinated output of the MYC network (shown in Fig. 4D), via the ATF4 arm of the UPR and the ISR, where MondoA plays an obligatory role in the induction of ATF4 under MYC-initiated stress.

To further test the function of the ISR in this system, we determined the effects of knocking down DEGs down-regulated (ARRDC4, TXNIP), or upregulated (ATF4, PPP1R15A; encoding GADD34) following MondoA-MLX perturbation (see Fig. 3) to gauge their effect on viability (Fig. S4A). siRNAs against ARRDC4, TXNIP, or GADD34 had only non-significant or borderline effects on cell growth, while siRNA targeting ATF4 markedly compromised viability in PSN-1 cells to a similar extent as siMondoA. The ATF4 dependence of these cells is consistent with the increased reliance on the ISR when MYC is amplified. Taken together, our expression and biological data indicate that MondoA-MLX functions within the ISR by both repressing and activating ISR components to regulate metabolic adaptation to stress.

### Loss of MondoA leads to decreased ATF4 protein expression

As ATF4 is the primary downstream stress responsive transcription factor cooperatively affected by perturbation of MYC and MondoA (Fig. 4C, Fig. S4A), we further characterized their crosstalk with the ISR, which regulates ATF4. The ISR responds to a variety of cellular stressors and, through stress-induced downstream phosphorylation of eIF2α, inhibits global protein synthesis while permitting preferential translation of the master-regulator ATF4 through its normally inhibitory uORF2 [19](Fig. 4D). **I**n both the PSN-1 and DAN-G cell lines, we found ATF4 mRNA upregulated ∼1.5-fold upon siMondoA treatment (Fig. 4E). Furthermore, eIF2α phosphorylation is readily induced following Tm treatment in both siControl and siMondoA-treated PSN-1 cells (Fig. 4F). However, ATF4 protein levels were significantly reduced after Tm challenge in siMondoA cells relative to siControls (Fig. 4F, replicate quantification Fig. S4B). To test whether this phenomenon contributes to the diminished viability phenotype seen with MondoA loss, we measured cell confluency with either individual siRNA knockdown of MondoA or ATF4, overexpression of ATF4, or a combination of MondoA knockdown and ATF4 overexpression (Fig. 5A). While loss of ATF4 leads to a similar decrease in growth as MondoA knockdown, ectopic overexpression of ATF4 was sufficient to rescue growth of siMondoA-treated cells (Fig. 5A). Rescue was also observed in a clonogenicity assay in ATF4 overexpressing PSN-1 cells treated with MondoA inhibitors SBI-477 or SBI-993 (Fig. 5B-C). Together, these data indicate that proper induction of ATF4 protein and its subsequent downstream program is a significant driver of MondoA dependency in the PSN-1 cells.

**Figure 5.**
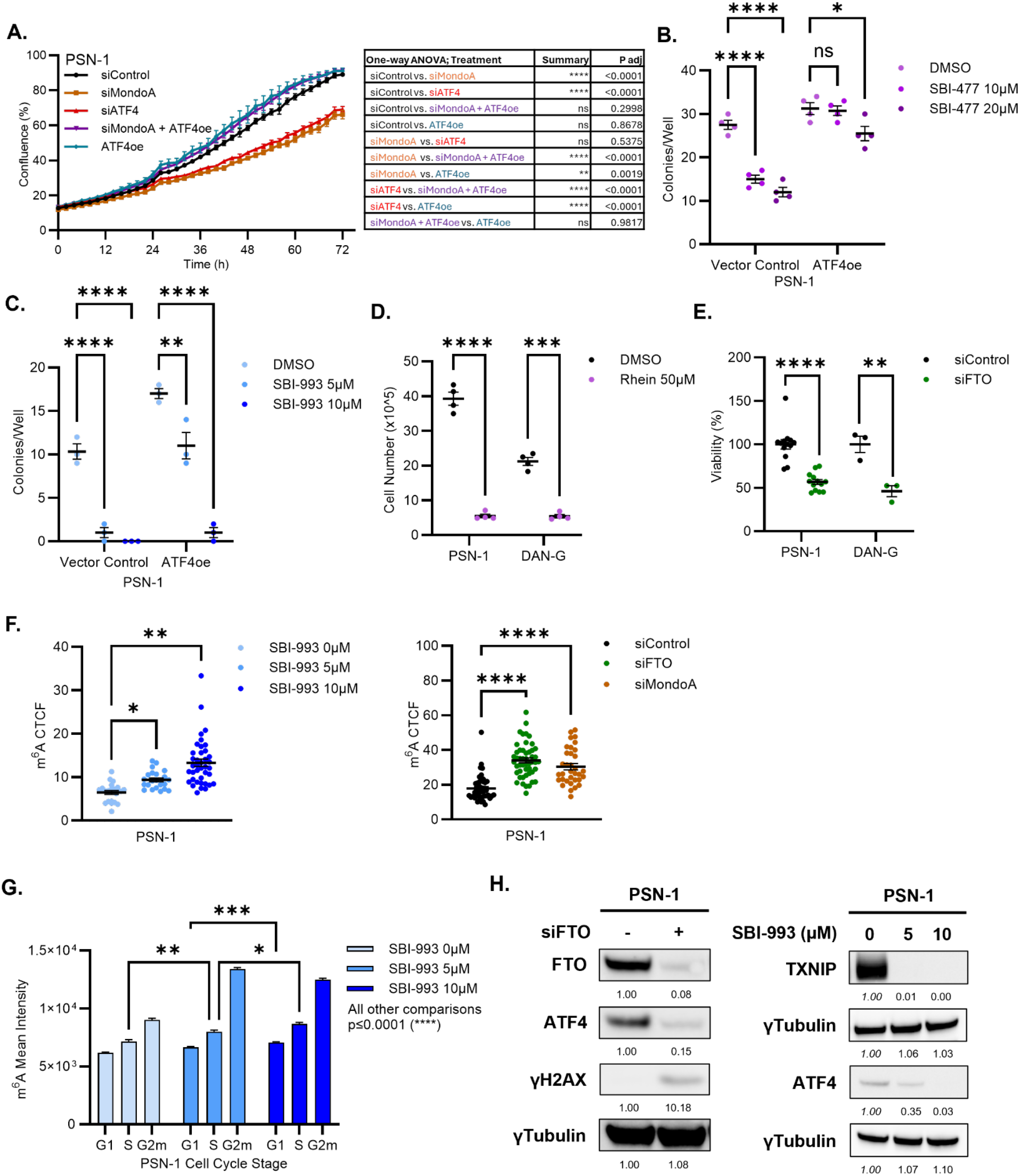
Assessing the role of ATF4 as a critical downstream effector of MondoA. (A) Incucyte growth curves for PSN-1 cells with siMondoA, siATF4, combination siMondoA and ATF4 overexpression vector (ATF4oe) (all n=8), or just ATF4oe (n=7) over 72 hours. Data plotted as mean percent plate confluency. Significance (one-way ANOVA) denoted in accompanying table. (B) Clonogenicity assay for PSN-1 cells with a control or ATF4oe vector seeded at 100 cells after 12 days and treated with indicated amounts of SBI-477 in four wells or (C) with indicated amounts of SBI-993 in three wells. Data expressed as colonies per well, significant differences (one-way ANOVA) expressed as compared to DMSO treatment within lines. For 5B, comparison between vector control and ATF4oe at both 10 and 20µM p<0.0001. For 5C, comparison between vector control and ATF4oe at 5µM is p<0.0001. (D) Viability assay for PDAC cells treated with DMSO or 50 µM rhein for 72 hours. Data expressed as mean cell number, significant differences (n=4, one-way ANOVA) expressed as compared to DMSO treatment within lines. (E) Viability assay for PSN-1 (n=13) or DAN-G (n=3) cells with siControl or siFTO treatment for 72 hours read as percent of siControl cell count. Significant differences (two-tailed t-test) expressed as compared to siControl within lines. (F) Relative global RNA m^6^A modification level as determined by immunofluorescent corrected total cell fluorescence (CTCF) for PSN-1 cells treated with siControl, siFTO, siMondoA, or indicated amounts of SBI-993 for 72 hours. Significant differences (one-way ANOVA) expressed as compared to siControl or DMSO treatment. (G) Mean m^6^A intensity by flow cytometry in PSN-1 cells treated with SBI-993 (0, 5, 10 µM) for 72 hours and called for cell cycle stage by DAPI stain. Significant differences (not labeled p≤0.0001, 2-way ANOVA) expressed between all doses and cell cycle stages. (H) Western blots for treatment target readout in PSN-1 cells treated with either siFTO or SBI-993 for 72 hours. Relative protein abundance expressed in relation to control treatment (siControl or DMSO) and normalized to γTubulin. All data expressed as mean±SEM and p-values expressed as ns= not significant; * p≤0.05; ** p≤0.005, *** p≤0.0005, **** p≤0.0001.

In order to gain mechanistic insight into the connection between ATF4 dysregulation and MondoA loss, we surveyed our RNA-seq signatures for perturbation of pathways potentially regulating ATF4, focusing on m^6^A-dependent RNA methylation which has been previously implicated in ATF4 translation[20]. N6-methyladenosine (m^6^A) modification of RNA is regulated by a methyltransferase complex (writers), demethylase enzymes (erasers), and various RNA binding proteins (readers). Dysregulation of this machinery has been shown to impact every stage of RNA processing in a wide range of tumor cells, including pancreatic cancer where dysregulation of the m^6^A pathway is linked to tumor progression and maintenance (reviewed in [21, 22]). As shown in the box plots in Figure S4C, multiple components of the m^6^A pathway are altered upon inhibition of MondoA-MLX via siRNA and/or SBI-477, most notably down-regulation of the primary mRNA demethylase FTO in all three conditions. Inhibition of RNA demethylases ALKBH5 and FTO by the inhibitor rhein (Fig. 5D) and siRNA knockdown of FTO (Fig. 5E) led to a significant loss of viability in the PSN-1 and DAN-G cells. We also observed a global 1.5-2-fold increase in m^6^A RNA modification by immunofluorescent staining in cells with siRNA knockdown of either FTO or MondoA, as well as a dose-dependent increase in cells treated with SBI-993 (Fig. 5F, representative images Fig. S4D). As m^6^A levels in RNA have been shown to vary across the cell cycle[23], we surveyed global m^6^A modification via flow cytometry and observed a dose-dependent increase in RNA m^6^A modification independent of cell cycle stage when treated with SBI-993 (Fig. 5G).To test whether perturbation of m^6^A machinery was sufficient to cause ATF4 protein loss, we determined ATF4 levels in PSN-1 cells and observed a significant loss of ATF4 protein both with siFTO and, in a dose-dependent manner, with SBI-993 treatment (Fig. 5H, S4E-F). We conclude that MondoA is necessary for the induction of the major ISR effector ATF4 at the protein level likely through crosstalk with the machinery of the mRNA m^6^A modification network (aka epi-transcriptome).

### Targeting MondoA in patient-derived organoids

PDAC patient-derived organoids (PDOs) recapitulate many features of the original disease and are suitable for testing personalized medicine and drug sensitivity [24]. Compared to normal pancreatic organoids, PDAC PDOs exhibit a gene expression signature that includes activation of MYC and E2F targets, concomitant with suppression of the TNFα pathway [25]. As these pathways are affected by MondoA inhibition in the PSN-1 cells, we investigated the feasibility of targeting MondoA via SBI-993 in PDAC PDOs. Six of the seven tested patient-derived organoids are sensitive to SBI-993 treatment (Fig. 6A, all images Fig. S5A), with faster growing PDOs exhibiting higher sensitivity to MondoA inhibition. Representative insensitive and sensitive organoid images and growth curves are shown in Figure 6B-C and S5B. Importantly, we confirmed target engagement in one of the sensitive lines, as the MondoA target TXNIP is decreased in a dose-dependent manner by SBI-993 (Fig. 6D).

**Figure 6.**
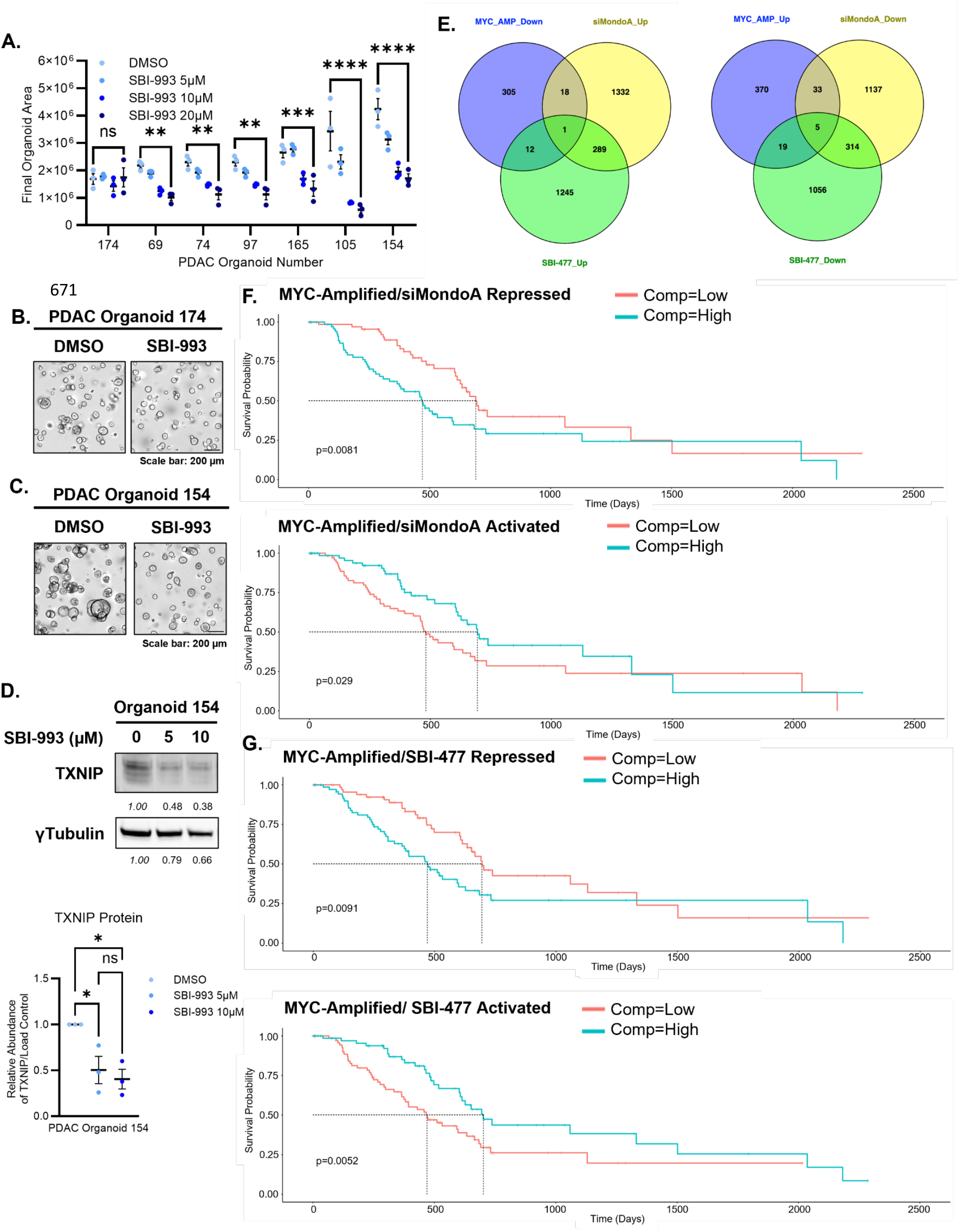
Effect of MondoA inhibition in patient-derived organoids and relationship to TCGA patient data. (A) Final organoid area µm^2^±SEM for PDAC Patient Derived Organoids (PDOs) after indicated doses of SBI-993 for 7 days. Significant differences (two-way ANOVA) expressed for 20µM dose as compared to DMSO treatment within PDOs. (B) Representative Incucyte final images for SBI-993 insensitive PDAC PDO 174 and (C) SBI-993 sensitive PDAC PDO 154 with 20µM of SBI-993. Scale bar 200µm. (D) Western blot to confirm SBI-993 target engagement as determined by TXNIP expression in PDAC PDO 154 with SBI-993 for 3 days. Relative protein abundance expressed in relation to DMSO treatment and normalized to relative abundance of γTubulin. Normalized mean protein abundance for TXNIP, significant differences (n=3, one-way ANOVA) expressed compared within doses of SBI-993. (E) Intersection between TCGA signature of DEGs in MYC-amplified PDAC and RNA-seq datasets following siMondoA or SBI-477 treatment as generated in Figure 3A-C. (F) Kaplan Meyer survival curve comparing patients with low (red) versus high (blue) expression of the combined MYC-amplification activated and siMondoA/SBI-477 repressed gene set generated in Fig. 6E. (G) Kaplan Meyer survival curve comparing patients with low (red) versus high (blue) expression of the combined MYC-amplification repressed and siMondoA/SBI-477 activated gene set generated in Fig. 6E. All data expressed as mean±SEM and p-values expressed as ns= not significant; * p≤0.05; ** p≤0.005, *** p≤0.0005, **** p≤0.0001.

### Translational Relevance of MondoA in PDAC clinical data

To further establish the relevance of MondoA-regulated pathways and targets in MYC-amplified PDAC, we queried the TCGA for expression data in patients with MYC amplification. We juxtaposed our RNA-seq datasets of either siRNA against MondoA, or chemical inhibition of MondoA in the PSN-1 cells, against the significantly enriched transcripts up-or downregulated with MYC amplification in the patient data (Fig. 6E). This yielded signatures for MondoA-dependent putative MYC-regulated targets of interest: those of which are upregulated with MYC-amplification, but downregulated by MondoA inhibition, as well as those repressed by MondoA, and enriched in the non-MYC-amplified patients (Supplemental Table 3). These “signatures” were then assessed for survival differences in the patient data of the pdacR database [26]. As shown in Figures 6F-G (Kaplan-Meier plots), these 4 signatures predict overall survival, with high expression of MondoA/MYC-amplified positively regulated genes correlating with poor survival, and high levels of the repressed genes correlating with better survival. This supports a role for MondoA in maintaining the MYC-amplified transcriptional program by enabling proper activation and repression via MYC in patients.

## Discussion

In this report, we have characterized functional coordination between MondoA and other components of the MYC network which directly contribute to MYC-driven tumorigenesis and survival in PDAC. We have found that MYC amplification or ectopic expression is synthetically lethal with loss of MondoA-MLX. Suppression of MondoA expression, or abrogation of MondoA transcriptional activity by means of a small molecule inhibitor, leads to decreased protein expression of the critical stress responsive transcription factor ATF4, altered MYC-ATF4 shared target gene expression, and cell death. Importantly, viability is rescued by the enforced expression of ATF4 in cells lacking MondoA expression or transcriptional activity. This work highlights a critical point of intersection and dependency between MYC, MondoA, and the ATF4-driven ISR.

Our initial report of MYCN-MondoA cooperation in neuroblastoma suggested that MondoA would be broadly required for MYCN function at a large proportion of shared target genes, many of which regulate metabolic processes [3]. Our present study using PDAC cells implicates both direct coordination at a limited subset of shared promoters (e.g. TXNIP, GADD34) as well as a dependency upon MondoA for proper genomic engagement of MYC and its extended network, even at promoters not normally bound directly by MondoA. This is consistent with a report of overlapping MYC and MondoA genomic binding in B-ALL, in which loss of MondoA profoundly affects MYC genomic occupancy, despite normal occupation of MondoA at only a fraction (<20%) of total MYC-bound promoters [27]. Such indirect regulation could be due not just to the loss of MondoA but also to the lack of ATF4 or other MondoA-regulated proteins that act as transcriptional cofactors with MYC at shared targets [9]. In addition, we observed a reconfiguration of MNT and ChREBP genomic occupation in response to loss of MondoA, changes that may be either secondary to, or directly contribute to altered MYC binding.

This work supports a model in which the MLX arm of the MYC network modulates MYC transcriptional output, mediating both hypo- and hyper-activation of MYC targets. MondoA maintains both MYC and MNT co-occupancy of shared and non-shared targets, possibly by heterodimerizing with MLX and thereby controlling the availability of MLX for dimerization with MNT and ChREBP. The balance between MLX- and MAX-containing heterodimers may represent a network restricted source of context-dependent cooperation or antagonism to MYC-MAX. As PDAC plasticity is driven by the ability to modulate MYC levels, both at the protein as well as genomic ecDNA copy number [28], functional interactions within the network may explain part of the inherent plasticity at the genomic level where the ability to temper MYC output can act as a survival advantage. In the case of MYC amplified cells, proteostatic stress must be curbed by the remedial ISR to accommodate diminished capacity or lack of metabolic sufficiency. Thus, MondoA’s role in fine-tuning the MYC network output and properly inducing ATF4 for the remediation of stress becomes necessary for cell survival. A diagram summarizing these roles for MondoA in MYC-amplified PDAC is shown in Figure 7.

**Figure 7.**
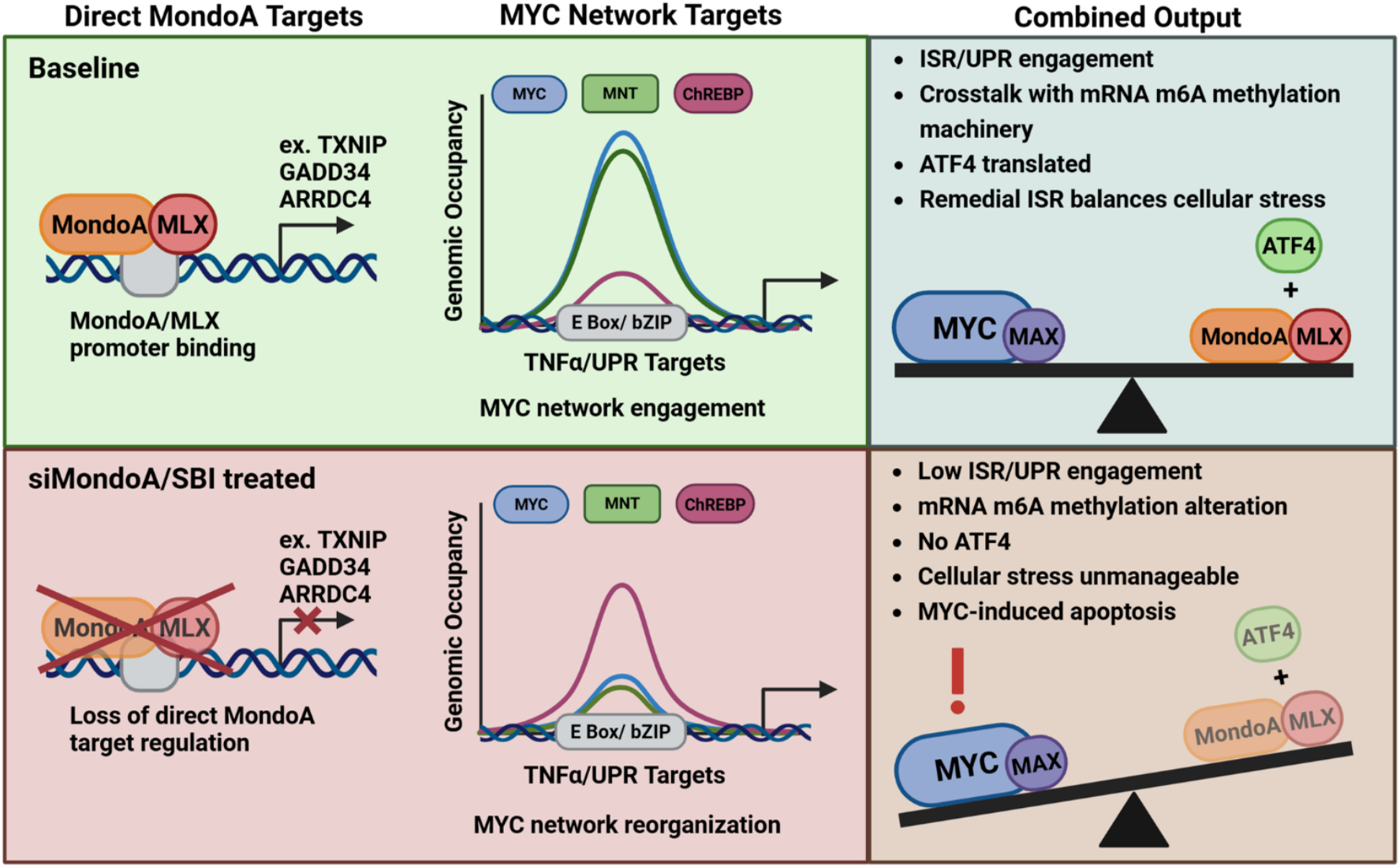
A model for MYC network transcriptional and genomic coordination with ATF4 in MYC amplified PDAC. Loss of MondoA leads to alterations in expression of direct MondoA-bound targets, as well as secondary effects on the genomic engagement of MYC network members, even at non-bound promoters. MYC and MNT are decreased, whereas the MondoA-paralog ChREBP, binding at UPR/TNFα pathway targets is enhanced. This co-occurs with loss of ATF4 protein, which may contribute to lack of MYC and MNT engagement at critical shared targets. This ultimately unbalances the tumorigenic and apoptosis-promoting effects of amplified MYC, resulting in non-remediated stress and cell death.

Our findings raise the possibility that inhibition of MondoA by means of a chemical inhibitor such as SBI-477/SBI-993 might be used to arrest the growth of PDAC and other neoplasms assessed as being addicted to high levels of MYC. However, a precision medicine approach to MondoA inhibition would require a deeper understanding of what levels and in what contexts MYC protein abundance activates the ISR to induce dependence on MondoA and its downstream effectors. MYC protein is present in both classical and basal PDAC as well as in normal pancreatic epithelium. More detailed studies will be required to determine the threshold above which MYC imposes sensitivity to MondoA loss. A related approach would be to target factors downstream of MondoA critical to the maintenance of dysregulated growth of tumor cells. Indeed, we find that a MondoA-regulated gene signature correlates with patient outcome. Moreover, we have shown that the FTO/ATF4 axis is engaged in PSN-1 cells by the MYC network via MondoA.

FTO is an important metabolic regulator in obesity and diabetes [29], as well as a modulator of chemotherapy-response and bZIP transcription factor activity in cancer[30]. FTO is a targetable oncogene in PDAC, where loss of function by knockdown or chemical inhibition impairs growth *in vitro* and *in vivo* [22, 29, 31]. These studies did not directly address the role of MYC in the sensitivity of PDAC cells to loss of FTO expression or activity, as none of the lines investigated exhibited amplification of the *MYC* locus, our work would suggest that MYC-amplification does not lead to resistance to such insults. The work presented here also implicates ATF4 as an important downstream effector of FTO in PDAC. It joins a host of other factors associated with adaptive response and cellular plasticity such as hypoxia, EMT, and pro-survival pathways mediating tumor progression and therapy resistance, all targeted by FTO and m^6^A in pancreatic cancer[22, 31].

ATF4 protein function is multifaceted and tightly controlled at multiple levels of regulation, culminating in either a remedial or terminal UPR [32]. This includes context- and cell-type-dependent regulation involving transcriptional, epi-transcriptional (mRNA methylation/demethylation via m^6^A) [20], translational (selective translation via uORF), protein stability (protein turnover/half-life), as well as availability of heterodimerization partnerships with other bZIP TFs required for the downstream transcriptional program of the ISR. MondoA coordinates these steps in MYC-driven PDAC. Global m^6^A levels across the transcriptome increase upon either loss of MondoA expression or MondoA-MLX targeting by a small molecule inhibitor. This is a profound phenotype for a transcription factor without demethylase activity to elicit in a cell upon loss of function. Future work will determine the specific nucleotides differentially modified by m^6^A, as well as their functional significance in mRNA stability or protein translation mediated by MondoA, potentially providing other therapeutic targets for existing, or yet to be discovered, inhibitors.

## Materials and Methods

All mice used in the study were housed and treated according to the guidelines provided by the Fred Hutch Institutional Animal Care and Use Committee (IACUC) protocol number: PROTO201900024. Complete Materials and Methods are included as a Supporting Information file (SI Appendix). Standard molecular biology and cell culture techniques were used, and kits and reagents were utilized following the manufacturer’s recommendation. Statistical comparisons and the tests for significance used for each experiment are indicated in the corresponding figure legends.

## Acknowledgments

We are grateful to Faiyaz Notta (Ontario Institute for Cancer Research) for his kind gift of patient-derived organoids, Daniel Kelly (University of Pennsylvania) for SBI-477 and discussions about MondoA inhibition, and to Phillp Norwood for technical assistance with aspects of this research. This work was supported by grants NIH/NCI R35 CA231989 (to R.N.E) and R37 CA241472 (to S.K.), an American Cancer Society Postdoctoral fellowship (PF-24-1196662-01-RMC) and Walter Benjamin Fellowship (465590102) from the German Research Foundation (to S.D.). Genomics and Bioinformatics analysis was funded by the Fred Hutch/University of Washington/Seattle Children’s Cancer Consortium (P30 CA015704), as well as scientific computing infrastructure supported by the Office of Research Infrastructure Programs (S10OD028685). We acknowledge the work of Fred Hutch Comparative Medicine (RRID:SCR_022610).

This manuscript is the result of funding in whole or in part by the National Institutes of Health (NIH). It is subject to the NIH Public Access Policy. Through acceptance of this federal funding, NIH has been given a right to make this manuscript publicly available in PubMed Central upon the Official Date of Publication, as defined by NIH.

