## Supplemental Info, Figs S1-S5 for "MondoA mediates transcriptional coordination between the MYC network and the integrated stress response in pancreatic ductal adenocarcinoma"

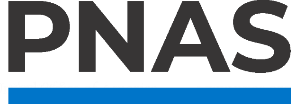

**Supporting Information for**

MondoA mediates transcriptional coordination between the MYC network and the integrated stress response in pancreatic ductal adenocarcinoma.

Paste the full author list here Erin L. Ramsey^1^, Stephanie Dobersch^2^, Brian Freie^1^, Nan Hyung Hong^1^, Xiaoying Wu^1^, Sita Kugel^2^, Robert N. Eisenman^1,^*, Patrick A. Carroll^1,^*

^1^Basic Sciences Division, Fred Hutchinson Cancer Center, Seattle, USA

^2^Human Biology Division, Fred Hutchinson Cancer Center, Seattle, USA

*Corresponding authors Robert N. Eisenman and Patrick A. Carroll

**This PDF file includes:**

Supporting Materials and Methods

Figures S1 to S5

SI References

Supporting Materials and Methods

Animal Use Statement:

All mice used in the study were housed and treated according to the guidelines provided by the Fred Hutch Institutional Animal Care and Use Committee (IACUC) protocol number: PROTO201900024.

Cell Lines and Culture Conditions

The following cell lines were obtained from ATCC and routinely screened for mycoplasma: PSN-1 (RRID:CVCL_1644), DAN-G (RRID:CVCL_0243) PANC08.13 (RRID:CVCL_1638), SU.86.86 (RRID:CVCL_3881) and HEK293FT (RRID:CVCL_6911). PDAC lines were cultured in RPMI (Gibco) supplemented with Pen/Strep (Gibco) and 10% FBS (Gibco), while the HEK293FT cells were cultured in DMEM (Gibco) with Pen/Strep and 10% FBS. Cells were washed with PBS (Gibco) and passaged by treatment with Trypsin (Gibco). Cell counting and Viability was monitored by Trypan Blue (Gibco) exclusion assay and cell counting via hemocytometer.

Chemical Stock Solutions

The following chemicals dissolved in the indicated solvents were used to treat cultured cells in this study: Dimethyl sulfoxide (DMSO) tissue culture grade (Santa Cruz Biotechnology), Doxycycline (Research Products International) was dissolved in water (1mg/ml stock) and added fresh every 2 days, Geneticin G418-Suifate (Gibco) was purchased as a 50mg/ml stock, Puromycin (Fisher Scientific) was dissolved in water (1M stock), Rhein (R&D Systems) was dissolved in DMSO (5mM stock), SBI-477/SBI-993 (MedChem Express) was dissolved in DMSO (10mM stock) and Tunicamycin (Cell Signaling) was dissolved in DMSO (5mg/ml).

Knockdown with siRNA

Cells were plated and transfected using Lipofectamine RNAiMAX (Invitrogen) following manufacturer recommendation for reverse transfection. Each experiment was carried out with a gene specific mixture of 4 siRNAs purchased from Qiagen and detailed information for all is included in table S4. Cells were incubated at standard conditions described above after transfection for 72 hours before cell counting and harvest.

Incucyte time-lapse imaging and quantification

The IncuCyte SX3 or SX5 live cell imaging and analysis system (Keyence) was used for tissue culture growth curves (confluency) as well as for patient-derived organoid growth (organoid area) quantification following the manufacturer’s recommendations.

Western Blotting

Cells were lysed normalized to cell number in complete RIPA buffer (20 mM Tris-HCl, 150 mM NaCl, 1% NP-40, 1% NaDOC, 0.1% SDS, 1 mM DTT, 1 mM EDTA, complete protease inhibitor (Roche) and phosphatase inhibitor (Bio-Rad)) at 100uL/1x10^6 cells and incubated with 4X Loading Buffer (Invitrogen) plus 1/10^th^ volume Beta-mercaptoethanol (BioRad). Samples were run on 4-12% Bis-Tris Gels (Invitrogen) in MES buffer (195.2 mg/mL MES (Millipore), 121.2 mg/mL Tris (ChemCruz), 20 mg/mL SDS (Sigma Aldrich), 6 mg/mL EDTA (Sigma Aldrich)) alongside a broad range protein ladder (ThermoScientific). Blots were transferred in Transfer Buffer (3 mg/mL Tris (ChemCruz), 14.4 mg/mL Glycine (RPI), 5 µl/mL 20% (v/v) SDS (SigmaAldrich), 200 µl/mL Methanol (Fisher Chemical)), onto a 0.2 µm Nitrocellulose membrane (BioRad) overnight at 40V 4C before ponceau stain (Fisher Scientific) and blocking with 5% non-fat dry milk (Apex) in TBST (1.21 mg/mL Tris (ChemCruz), 8.76 mg/mL NaCl (Fisher Bioreagents), 1 µL/mL Tween20 (SigmaAldrich)). Primary and secondary antibodies were diluted in 5% non-fat dry milk in TBST and incubated either overnight at 4C or at RT for 1 hour respectively. Antibody details can be found in table S5. For development, blots were incubated for 2 minutes at room temperature with Femto ECL (Thermo Scientific) and imaged on a ChemiDoc MP (BioRad).

Western Blot quantification

Images were uploaded into ImageJ[1] for quantification using the protocol from Hossein Davarinejad at York University[2]. To normalize protein quantity to load control, net protein of interest density was divided by net load control density. Further normalization to a control sample was performed for each blot by dividing net density of each sample by the control sample, effectively setting the control sample to a relative quantity of 100% and expressing relative quantity of all other bands as a percentage compared to control sample.

Lentiviral Production

HEK293FT cells were transfected with Lipofectamine 2000 (Invitrogen) according to the manufacturer’s recommendations with the following packaging constructs: psPAX2 (Addgene plasmid #12260), and pCMV-VSV-G (Addgene plasmid #8454) in combination with one of the expression vectors. For Inducible shRNA system, SMARTvector inducible shRNA lentiviral system (Dharmacon), with either piSMART-hEF1a-TurboGFP-NTC (shNTC control), or piSMART-hEF1a-TurboGFP-shMondoA (target sequence AAGTTTGCTGGAGTCAACA) with puromycin selection were used. Lentivirus was then transduced into the PSN-1 cells with polybrene (Sigma-Aldrich) to generate the Dox-inducible shNTC and shMondoA lines. For the Inducible MYC system: a cDNA encoding human MYC was subcloned into the vector pCW57.1-T2A-GFP (a kind gift from Dr. Stephen Tapscott) with the addition of an N-terminal 3xFLAG tag to generate, pCW57.1-3xFLAG-MYC-WT. Lentivirus was produced in HEK293FT cells, then transduced into the SU.86.86 cell line to generate the Dox-inducible MYC line.

Apoptosis Assay

Cells were seeded at equal number in a 96 well plate via reverse transfection of the indicated siRNA. Apoptosis was monitored using the Caspase 3/7 System (Promega) normalized to cell number using Cell-Titer Glo (Promega).

Clonogenic Foci Formation Crystal Violet Stain

For siRNA treatment, cells were reverse transfected following the above protocol with either siControl or siMondoA 24 hours before seeding and then plated at 100, 200, or 500 cells per well in a 6-well plate. For SBI-477 and SBI-993 treatment, cells were plated at 100 cells and treated small molecule inhibitor. Plates were incubated at 37°C with 5% CO2 for 12 days with media and/or treatment replaced every 3-4 days. Foci formation was assessed by crystal violet stain (Millipore Sigma).

Xenograft

Female NSG (NOD.Cg-Prkdc^scid Il2rg^tm1Wjl/SzJ) mice, aged 6–8 weeks, were obtained from the Translational Research Model Services (TRMS) Core at Fred Hutch and housed in a pathogen-free facility under standardized conditions (12-hour light/dark cycle, ambient temperature of 24 °C [75 °F], and 30–70% relative humidity). Mice were pre-treated with 200g/L doxycycline in their drinking water for 1 week and for the duration of the study. For xenograft implantation, 1.3 × 10⁶ cells were suspended in 100 μL of a 1:1 mixture of cold Dulbecco’s phosphate-buffered saline (DPBS) and growth factor-reduced Matrigel (Corning) and injected subcutaneously into the left and right flanks of each mouse. PSN-1 cells expressing either dox-inducible shNTC or shMondoA were injected subcutaneously into the flanks of NSG mice (left flank shNTC, right flank shMondoA) for pair-wise comparisons. Tumor growth was monitored three times per week using digital calipers for up to 38 days post-implantation. Tumor volume was calculated using the ellipsoid formula:

$$\boldsymbol{Tumor Volume}\left( \boldsymbol{mm}^{\boldsymbol{3}} \right)\boldsymbol{=0.5 \times length\times}\boldsymbol{width}^{\boldsymbol{2}}$$

where length refers to the tumor’s head-to-tail axis and width to the front-to-back axis.

Mice were monitored closely for signs of distress or tumor-related morbidity. Humane endpoint criteria included tumor ulceration, necrosis, bleeding, infection, impaired mobility, difficulty accessing food or water, or disruption of vital physiological functions (e.g., respiration, excretion). As per institutional guidelines, maximum allowable tumor sizes were 2.0 cm in average diameter for unilateral tumors, or 1.5 cm per tumor for bilateral implants. Mice were euthanized immediately upon meeting any of these criteria. Upon completion of study, tumors were removed and weighed for final tumor weight quantification.

RNA-Sequencing

Total RNA was extracted with Trizol (Thermo-FIsher) and purified with the Direct-zol RNA miniprep Kit (Zymogen) following manufacturer recommendations. RNA integrity was monitored on a TapeStation 4200 (Agilent). TruSeq libraries were prepared from 500ng of RNA and paired-end NGS sequencing was carried out by the Fred Hutch Genomics Core on an HiSeq 2500 (Illumina). Raw files were aligned to human genome Hg38 with TopHat and analyzed with DESeq2 [3]. Volcano plots and Heatmaps were generated with ggPlot2 in R or NGS plot [4]. Pathway analysis was done with Enrichr [5].

Genomics (CUT&RUN)

Genomic binding was monitored by CUT&RUN [6] run on an automated platform through the Fred Hutch Genomics Core [7]. Samples from PSN-1 cells were prepared 48 hours post-siRNA transfection (1x10^6 cells per condition), and incubated overnight with antibodies against MondoA, MYC, MNT, ChREBP or control immunoglobulin (IgG). Duplicate samples were merged to obtain higher coverage datasets. Peaks were called using the MACS2 package in R. Library normalized Bigwig files were visualized as tracks in Integrative Genomic Viewer (IGV) [8].

TCGA Data Analysis

Patient data was analyzed either through The Cancer Genome Atlas (TCGA) cBioPortal [9] (<https://www.cbioportal.org/>) to sort patients on MYC-amplified versus non-amplified status, or the pdacR database [10] (<https://pdacr.bmi.emory.edu/>) to generate correlation plots and Kaplan-Meier survival curves.

Puromycin Incorporation Assay

Cells were treated with 1μM puromycin (Fisher Scientific) and incubated at 37°C for 15 minutes prior to collection. Cell lysates were prepared as described above for western blotting and probed with anti-puromycin.

Quantitative RT-PCR

Reverse transcription was performed using Superscript First-Strand Synthesis Kit (Invitrogen) and Quantitative PCR was performed using SYBER Green iTaq Supermix (BioRad) on an Icycler (BioRad) with the following primer pairs: ATF4 Forward: CCAAGCACTTCAAACCTCATG and ATF4 Reverse: ATCCATTTTCTCCAACATCCAATC; RPL0 Forward: GGCTGCGTCTATGGTCATGA and RPL0 Reverse: CGAGACTTTGGGTACGGCTT. Data was processed by the delta-delta cT method for relative fold-change quantification comparing siControl to siMondoA conditions.

Plasmid Transfection

Either the pRK5 empty vector (ATCC) or pRK-ATF4 construct (a gift from Yihong Ye, Addgene plasmid # 26114,[11]) were transfected into PSN-1 cells using Lipofectamine 3000 (Invitrogen) following the manufacturer’s recommendations. Cells were selected with Geneticin G418-sulftate (Thermo-Fisher) to generate the pRK5 vector and pRK5-ATF4-OE cell lines.

Immunofluorescent Staining and Quantification

Immunofluorescence was performed on cells fixed with 4% paraformaldehyde (PFA) in PBS. Cells were treated with a blocking and permeabilization buffer (B&P buffer) (4% g/mL IgG-free BSA, 0.2% Triton X-100, in PBS) before incubation with DAPI nucelar counterstain (Sigma-Aldrich, 0.1ug/mL), primary, and secondary antibodies diluted in B&P. For preserving fluorescence and imaging, cover slips were mounted on slides with Mounting Solution (ThermoFisher) and let dry overnight. Fluorescence images were obtained using a Leica DM IL LED microscope with a Leica K3C microscope camera at 20x/0.40 magnification. Antibodies used can be found in table S5.

FIJI ImageJ [1] was used to balance brightness and contrast of images, with identical adjustments across all images for each primary antibody. The FIJI freehand tool was used to select each cell, and capture the following measurements: cell area, mean gray value, and integrated density. Reported Integrated Density is the product of selection area and mean gray value. Five background readings were also recorded for normalization. Corrected Total Cell Fluorescence (CTCF) was then calculated to adjust for cell size and background variability using the following formula:

$$\boldsymbol{CTCF=Integrated Density-(Area\times Mean Background Fluorescence)}$$

Flow Cytometry

Intracellular staining was carried out on methanol-fixed PSN-1 using intracellular fixation buffer according to the manufacturer (eBiosciences) and co-stained with DAPI (Sigma-Aldrich, [1:20]). Antibody details can be found in table S5. Cells were run on a Fortessa X-50 Flow Cytometer (BD Biosciences) and cell-cycle stages were called by the DAPI cell cycle feature in FlowJo™ v10.10.0 Software (BD Biosciences).

Human Patient-Derived Organoids

Human pancreatic ductal adenocarcinoma (PDAC) organoids were generously provided by Dr. Faiyaz Notta [12]. Characterization can be found in table S6. Organoid culture was carried out following the standardized procedures outlined in the Tuveson Laboratory Human Organoid Protocols[13]. For passaging, organoids were enzymatically dissociated into single cells using TrypLE Express (Gibco). Cells were subsequently embedded in growth factor-reduced Matrigel (Corning) and plated as domes. Organoids were maintained in Human Feeding Medium, composed of Advanced DMEM/F12 (Invitrogen) supplemented with 1× HEPES, 1× Glutamax, 1× penicillin/streptomycin, 1× B27 (all from Invitrogen), and 1 mg/mL Primocin (InvivoGen). The medium was further enriched with 200 μg/mL N-acetyl-L-cysteine (Sigma), 20% (v/v) Wnt3a-conditioned medium, 30% (v/v) R-spondin1-conditioned medium, 0.1 μg/mL mouse Noggin (Peprotech), 50 ng/mL human EGF (Peprotech), 0.21 μg/mL human Gastrin (Tocris), 0.1 μg/mL human FGF10 (Peprotech), 0.045 mg/mL nicotinamide (Sigma), and 0.21 μg/mL A83-01 (Tocris). For SBI-993 treatment, cells were seeded in the indicated dose of SBI-993 or DMSO and allowed to grow for 8 days.

Statistical Analysis

Statistical analysis was done either using R based scripts or using Prism 10.2 (GraphPad). Student’s T test was used to compare groups of 2, whereas ANOVA was used for groups of 3 or more.

Network and Pathway Figure Creation

Explanatory figures and protein icons were generated using BioRender. Publication Licenses below:

Figure 2E and protein icons in 3E: Created in BioRender. Ramsey, E. (2025) <https://BioRender.com/s18l7yo>

Figure 4D: Created in BioRender. Ramsey, E. (2025) <https://BioRender.com/kxnmxay>

Figure 7: Created in BioRender. Ramsey, E. (2025) <https://BioRender.com/v7mt905>

**Supplemental Figures**

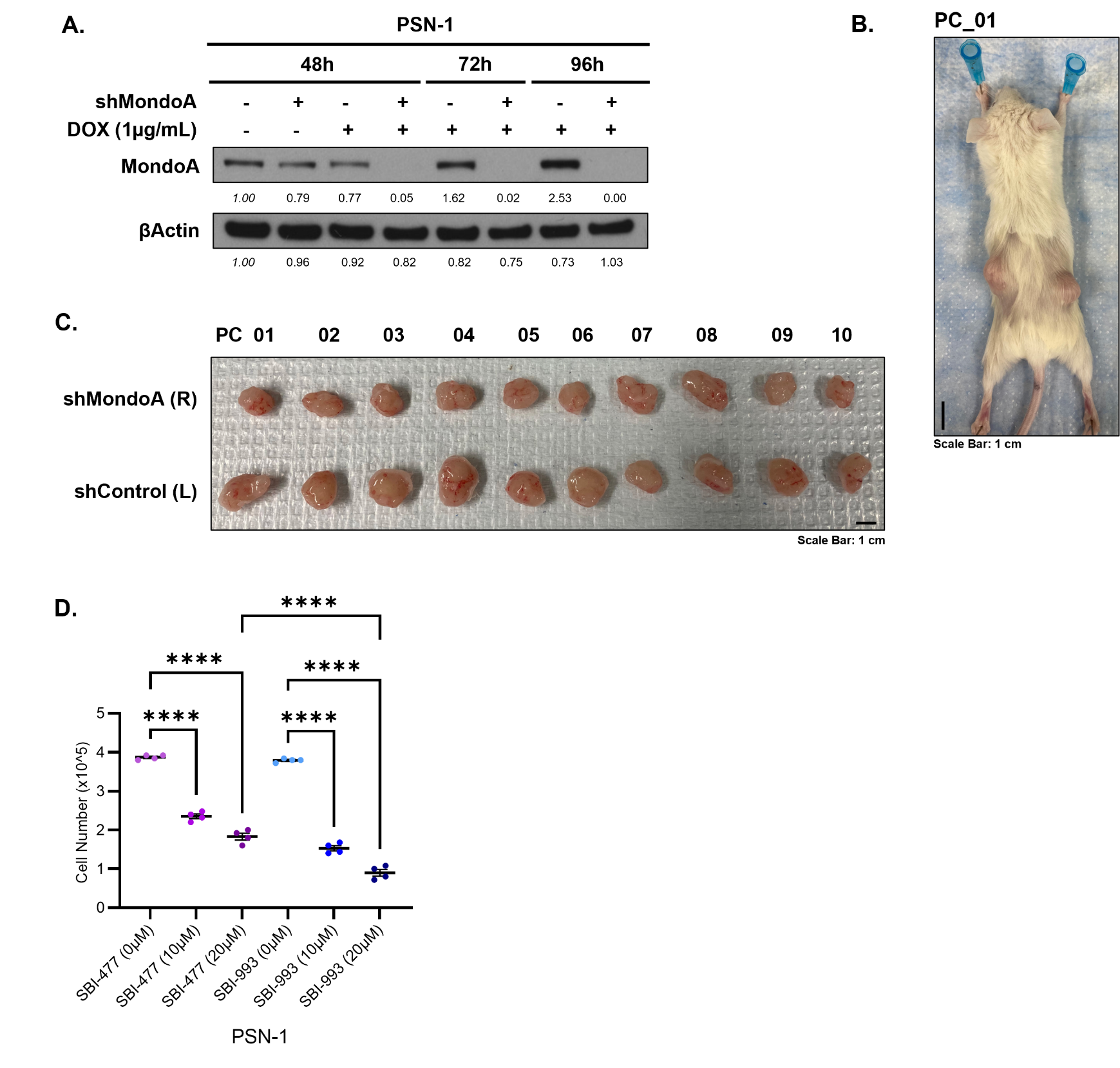
Figure S1. Targeting MondoA *in vitro* and *in vivo* in PSN-1 cells. (A) Western blot confirming DOX-induced expression of shMondoA and knockdown of MondoA protein in engineered PSN-1 cells at 48, 72, and 96 hours. Relative protein abundance expressed in relation to shControl without DOX at 48 hours and normalized to relative protein abundance of βActin. (B) Representative image of a mouse injected with PSN-1 subcutaneous xenograft containing shControl (left flank) or shMondoA (right flank) after 38 days. Scale bar 1 cm. (C) Images of final tumor size for PSN-1 subcutaneous xenograft with either shControl or shMondoA after 38 days. Scale bar 1 cm. (D) Viability assay for PSN-1 cells treated with 0, 10, 20 µM of either SBI-477 or SBI-993 for 96 hours. Data expressed as mean cell count, significant differences (n=4, one-way ANOVA) expressed as compared to DMSO treatment within each inhibitor and between inhibitors for 20 µM dose. All data expressed as mean±SEM and p-values expressed as ns= not significant; * p≤0.05; ** p≤0.005, *** p≤0.0005, **** p≤0.0001.

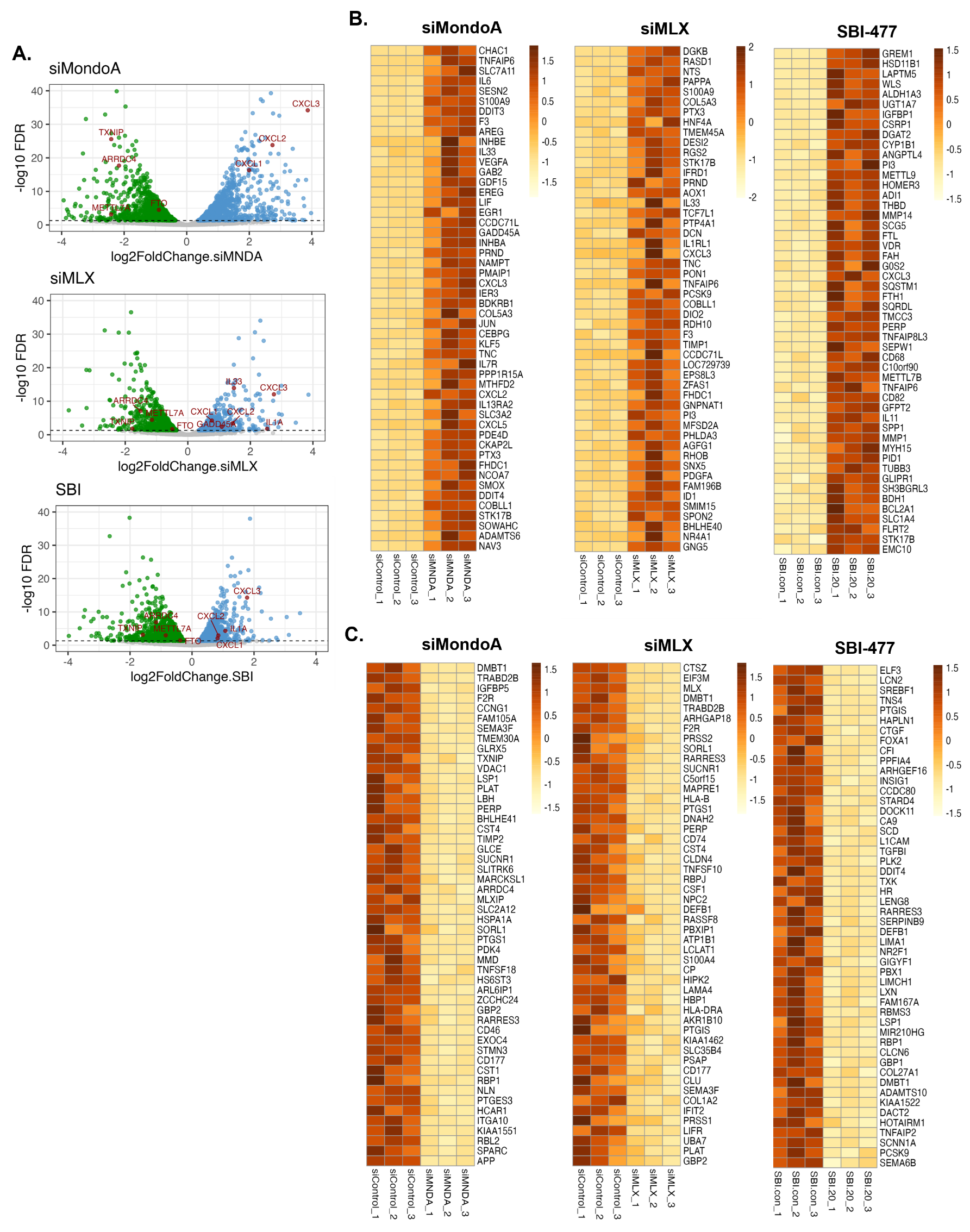

Figure S2. Transcriptomic analysis of MondoA-MLX knockdown and chemical inhibition. (A) Volcano plot of DEGs after siMondoA, siMLX, or SBI-477 treatment. Up-regulated genes in blue, down-regulated genes in green, and select ‘core’ DEGs highlighted in red. (B) Heatmap of the top 50 up-regulated DEGs across all three biological replicates for siMondoA, siMLX, and SBI-477. (C) Heatmap of the top 50 down-regulated DEGs across all three biological replicates for siMondoA, siMLX, and SBI-477.

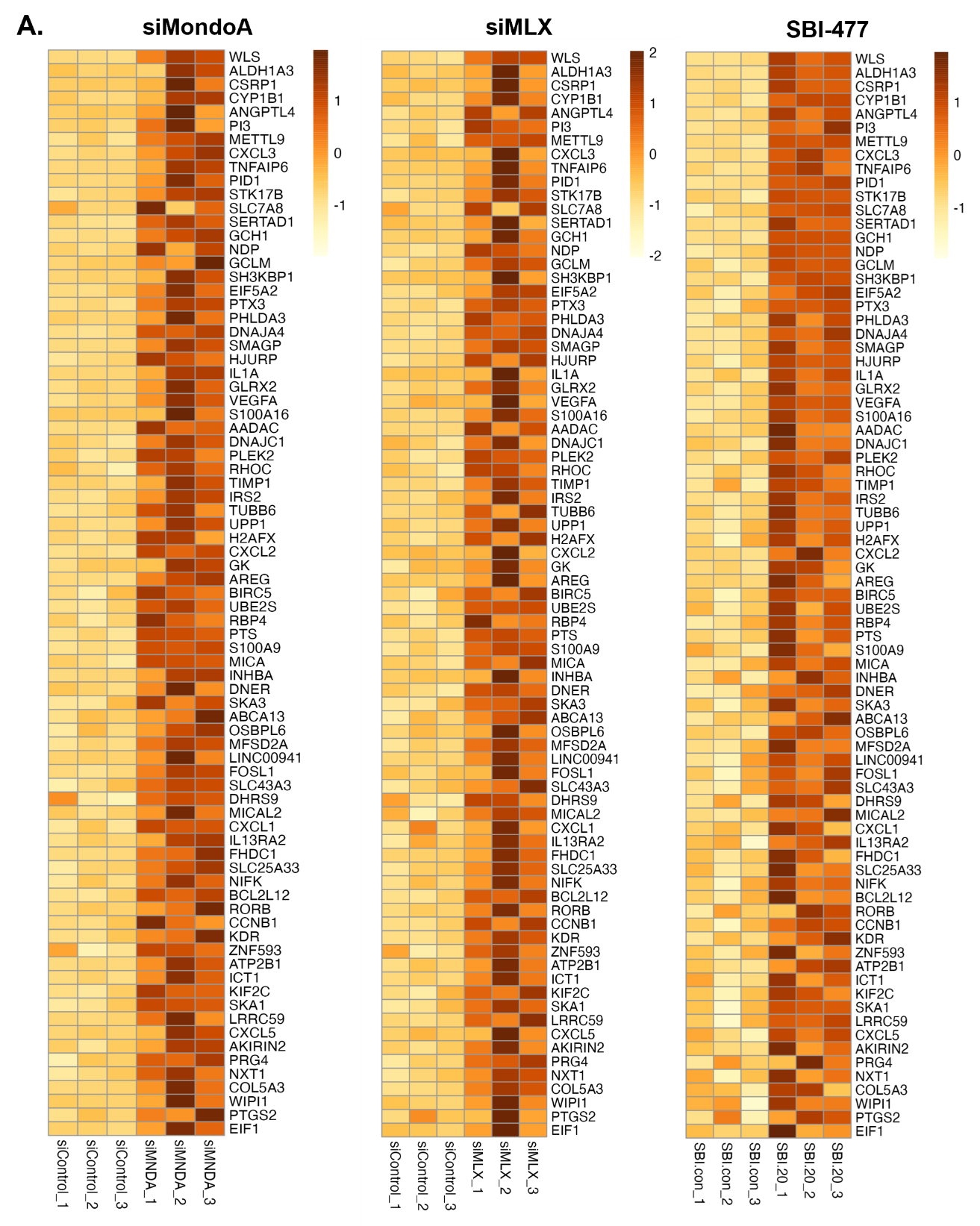

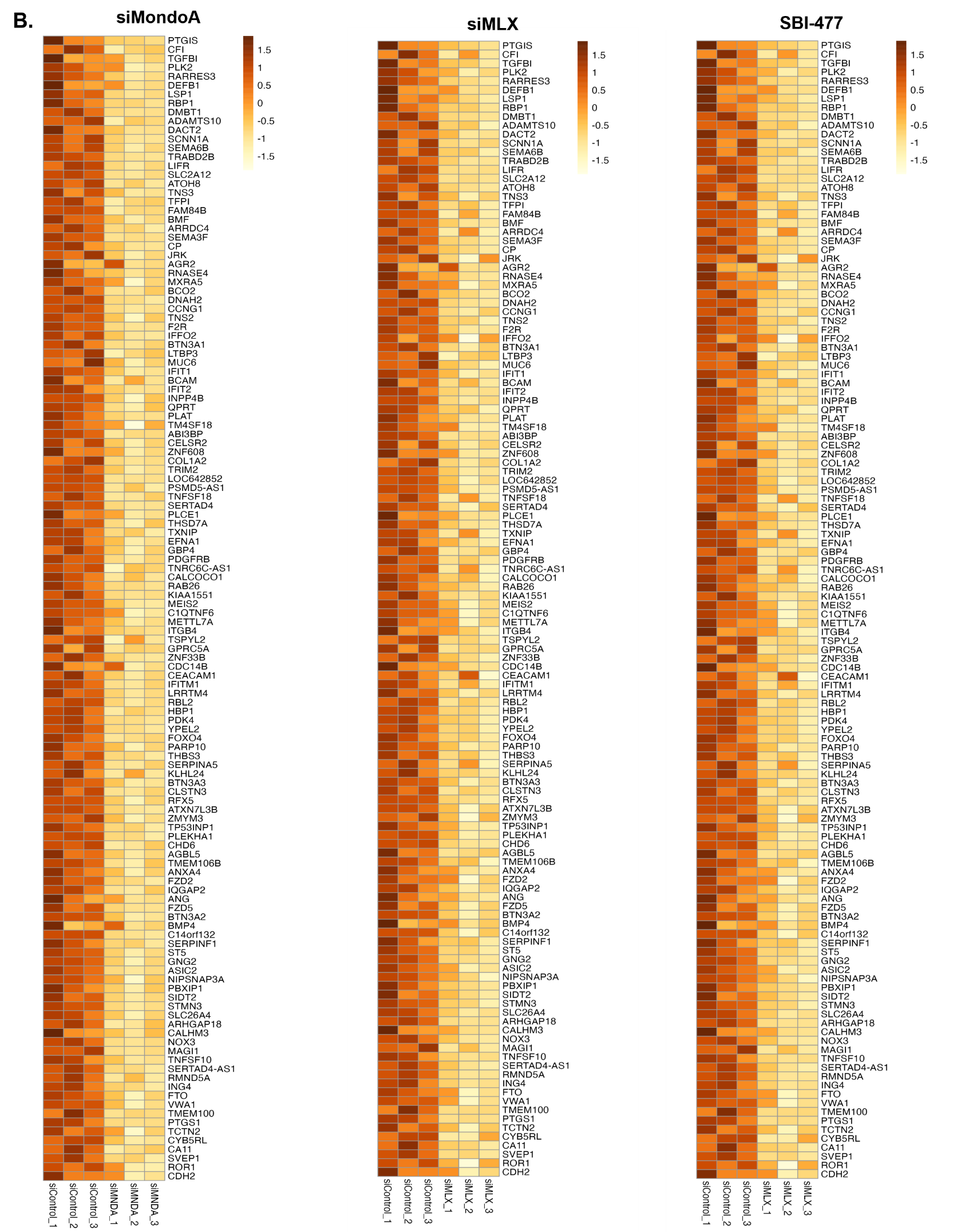

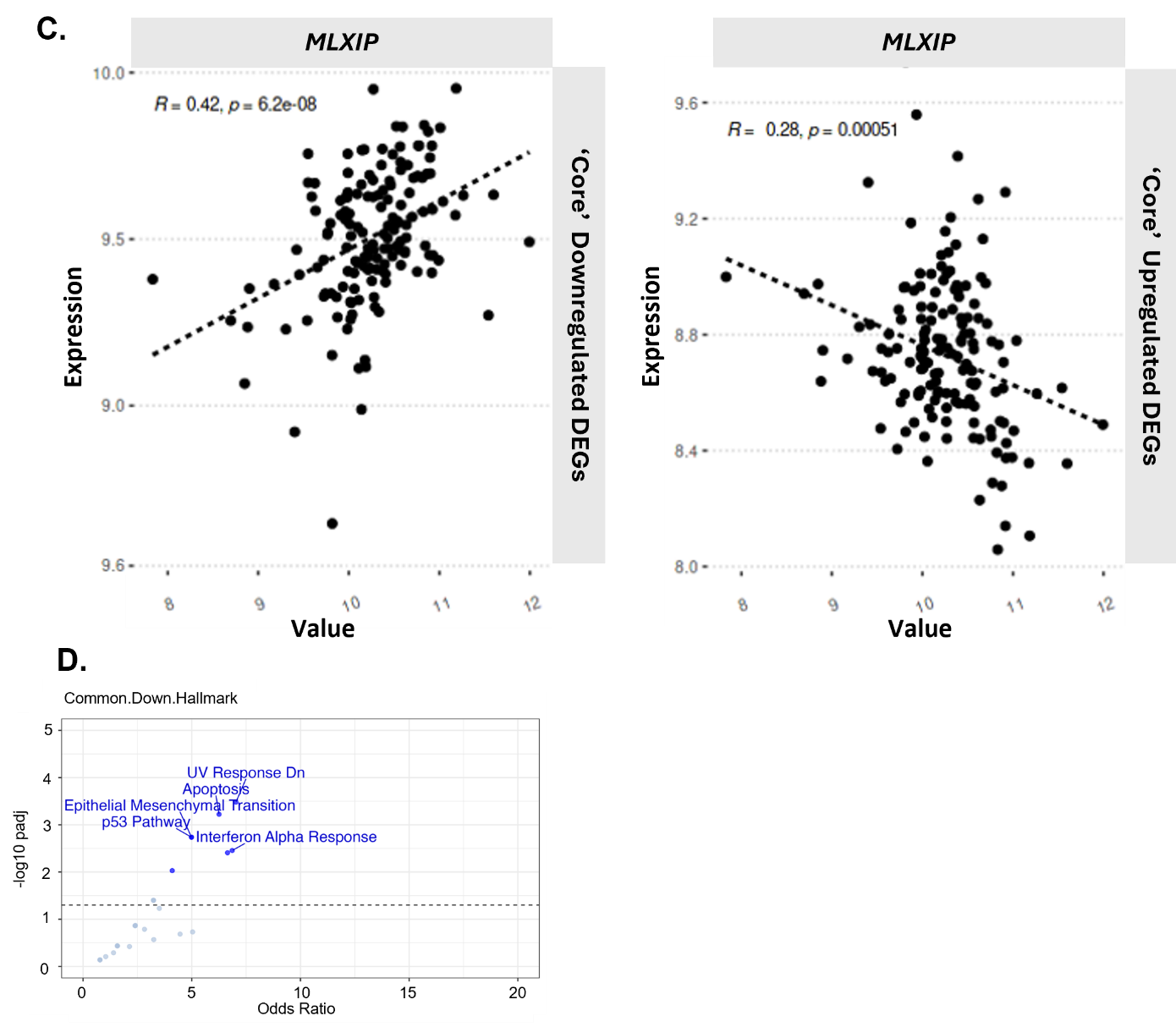

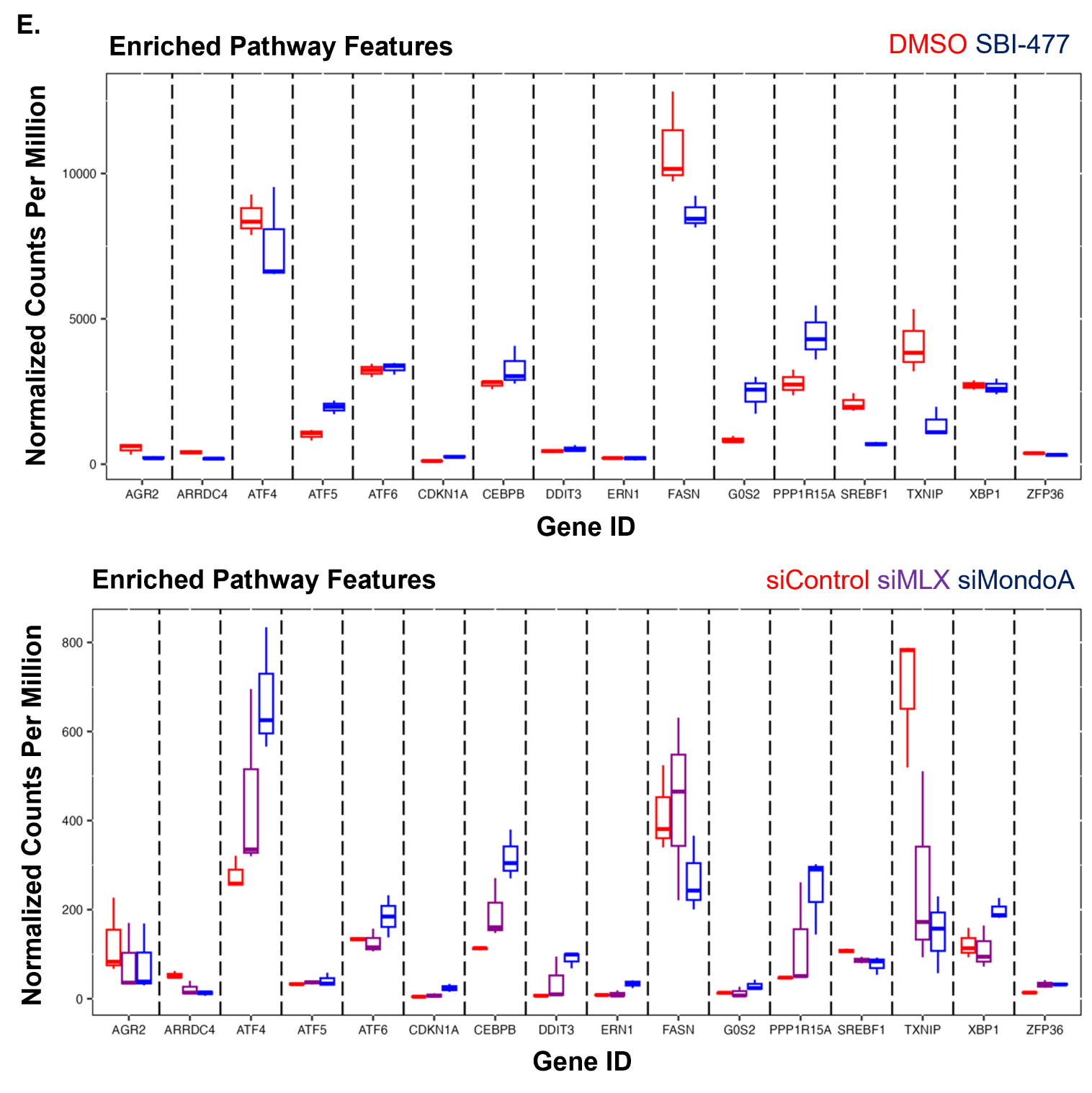
**Figure S3.** Comparative RNA-seq analysis identifies “core” MondoA regulon. (A) Heatmap of the ‘core’ up-regulated DEGs across all three biological replicates for siMondoA, siMLX, and SBI-477. (B) Heatmap of the ‘core’ down-regulated DEGs across all three biological replicates for siMondoA, siMLX, and SBI-477. (C) TCGA expression correlation between MondoA (MLXIP) and ‘core’ up and down regulated DEGs. (D) Pathway analysis of DEGs down-regulated in ‘core’ gene-list. (E) Box plot for key target pathway gene expression in RNA-seq datasets regulated by siMondoA, siMLX and SBI-477.

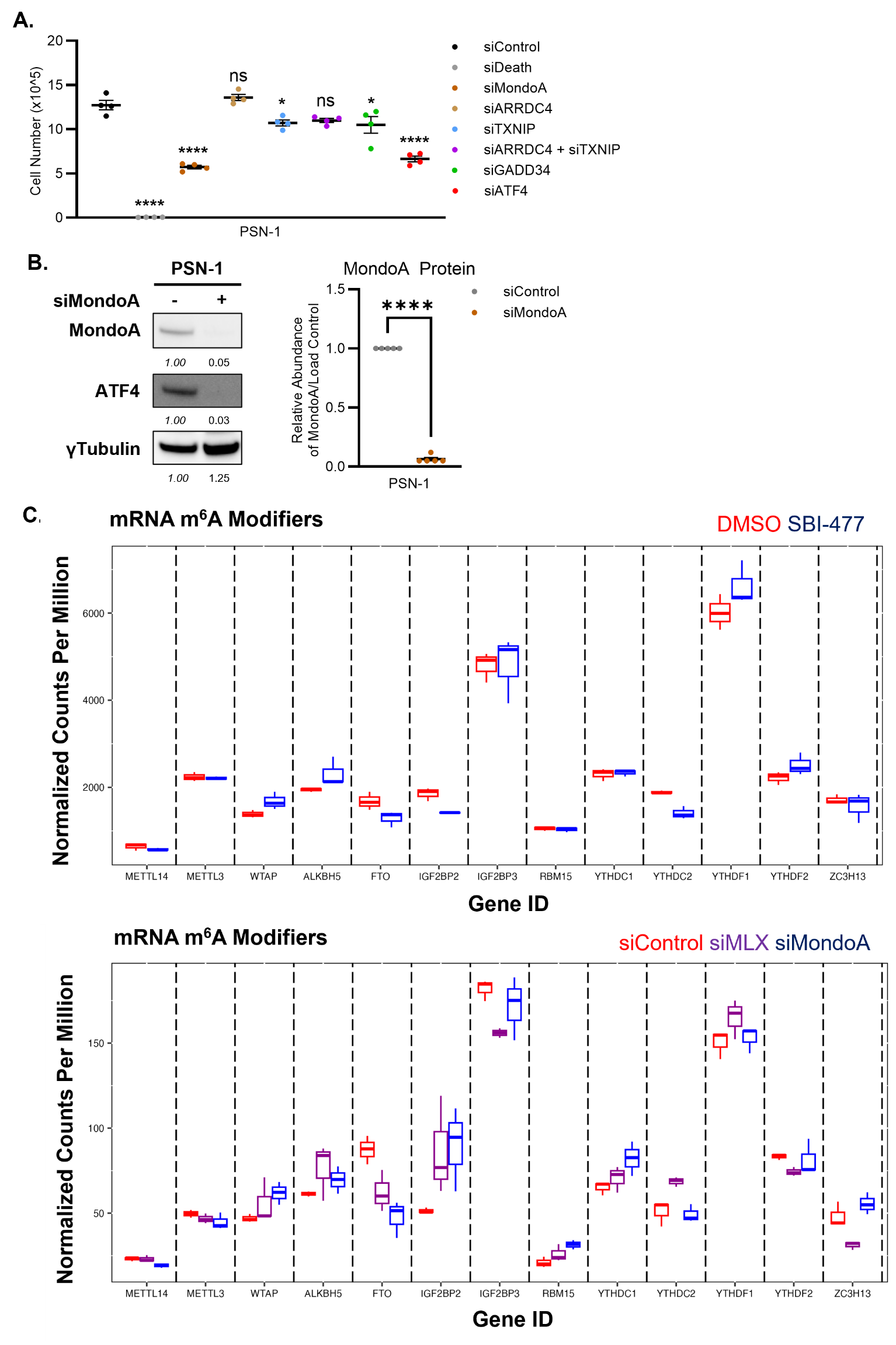

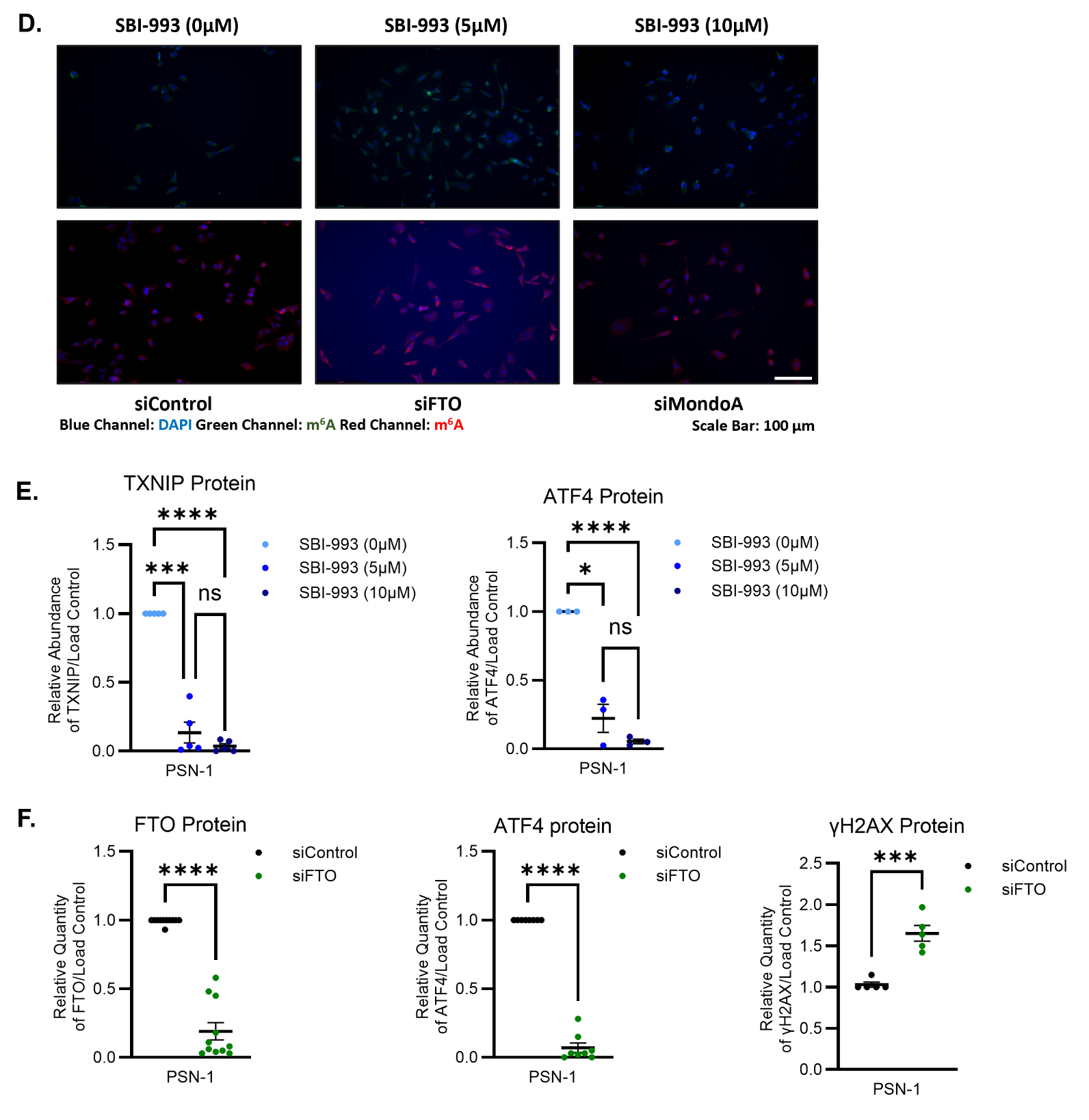
Figure S4. MondoA regulation and functional test of the UPR/ISR and m6A pathways in PDAC. (A) Viability expressed as mean cell number for PSN-1 cells treated with an siRNA screen of UPR effectors significantly changed in RNA-seq data from Figure 3 alone and in combination for 72 hours. Significant differences (n=4, one-way ANOVA) expressed as compared to siControl treated cells. (B) Representative western blot and quantification for MondoA and ATF4 protein in PSN-1 cells after 72 hours of siControl or siMondoA. Relative protein abundance expressed in relation to siControl treated cells and normalized to relative abundance of γTubulin. Significant differences (n=5, two-tailed t-test) for five expressed as compared to siControl treatment. (C) Box plot for key m^6^A modification pathway gene expression in RNA-seq datasets. (D) Representative images for immunofluorescent staining against DAPI (blue) and m^6^A (green and red) in PSN-1 cells treated with siFTO, siMondoA, or SBI-993 (0, 5, 10 µM) for 72 hours. Scale bar 100µm. (E) Western blot quantification of relative protein level for TXNIP (n=5) and ATF4 (n=3) in PSN-1 cells treated with 0, 5, or 10µM of SBI-993 for 72 hours. Data expressed as relative protein abundance in relation to DMSO treatment and normalized by relative abundance of load control. Significant differences (two-tailed t-test) expressed between all doses. (F) Western blot quantification of relative protein level for FTO (n=11), ATF4 (n=8) and γH2AX (n=5) in PSN-1 cells treated with siControl or siFTO for 72 hours. Data expressed as relative protein abundance in relation to siControl and normalized by relative abundance of load control. Significant differences (two-tailed t-test) expressed in relation to siControl. All data expressed as mean±SEM and p-values expressed as ns= not significant; * p≤0.05; ** p≤0.005, *** p≤0.0005, **** p≤0.0001.

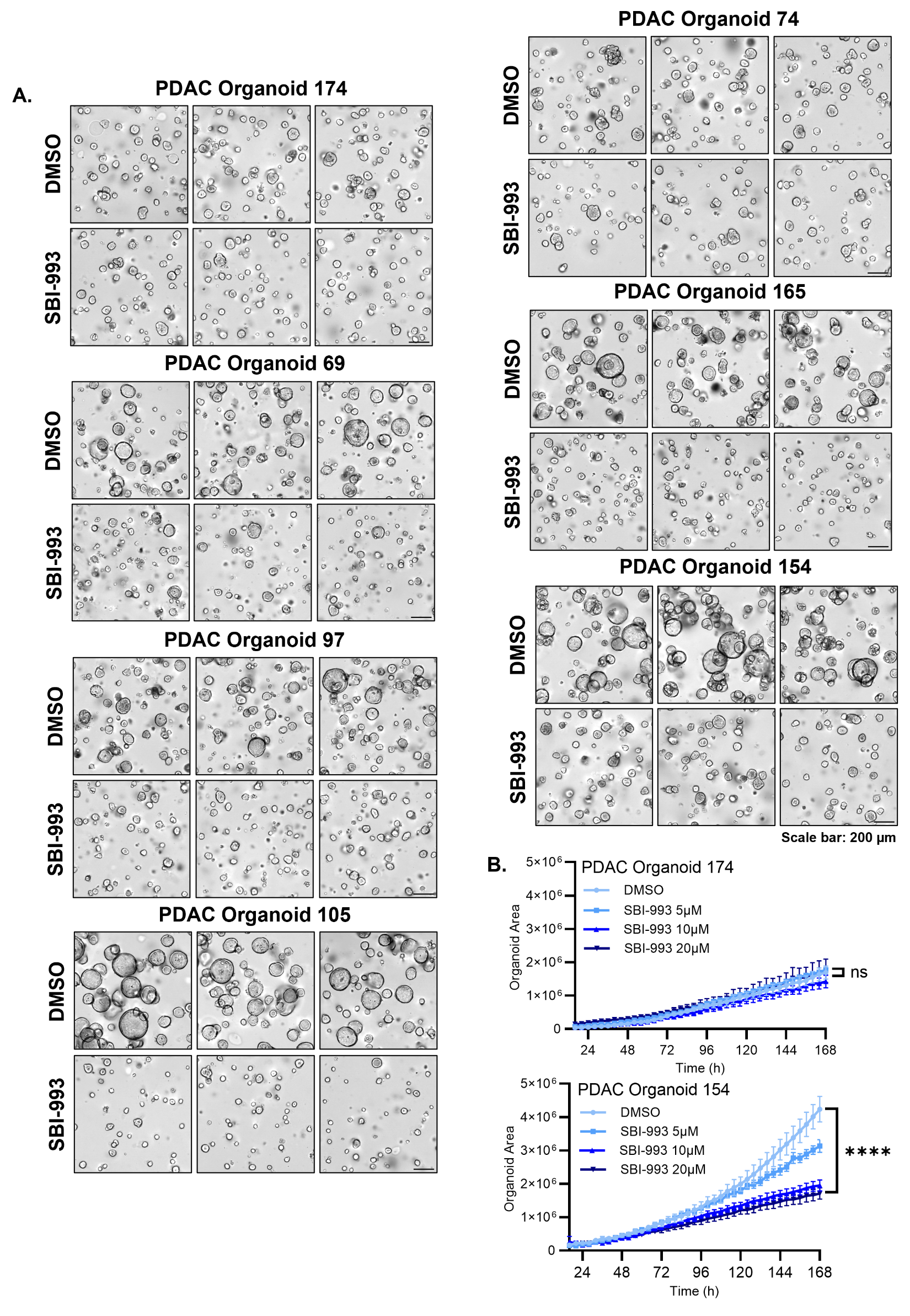

Figure S5. Targeting MondoA via SBI-993 in Patient-Derived Organoids. (A) Final Incucyte images for SBI-993 treated PDAC PDOs with 20µM of SBI-993. Scale bar 200µm. (B) Incucyte growth curve data for a representative unresponsive (PDO 174) and responsive (PDO 154) PDAC PDOs treated with 0, 5, 10, or 20 µM SBI-993 for 168 hours. Data expressed as mean incucyte area units, significant differences for final area (n=3, two-way ANOVA) expressed between 20µM of SBI-993 and DMSO treatment. All data expressed as mean±SEM and p-values expressed as ns= not significant; * p≤0.05; ** p≤0.005, *** p≤0.0005, **** p≤0.0001.
