## Supplemental Table 1 for "MondoA mediates transcriptional coordination between the MYC network and the integrated stress response in pancreatic ductal adenocarcinoma"

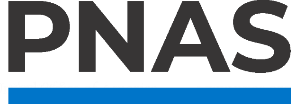

**Supporting Information for**

MondoA mediates transcriptional coordination between the MYC network and the integrated stress response in pancreatic ductal adenocarcinoma.

Paste the full author list here Erin L. Ramsey^1^, Stephanie Dobersch^2^, Brian Freie^1^, Nan Hyung Hong^1^, Xiaoying Wu^1^, Sita Kugel^2^, Robert N. Eisenman^1,^*, Patrick A. Carroll^1,^*

^1^Basic Sciences Division, Fred Hutchinson Cancer Center, Seattle, USA

^2^Human Biology Division, Fred Hutchinson Cancer Center, Seattle, USA

*Corresponding authors Robert N. Eisenman and Patrick A. Carroll

**This PDF file includes:**

Supplemental Table 1

**Supplemental Table 1: Upregulated DEGs from RNA-seq**

| **siMondoA Up** | **siMLX Up** | **SBI-477 Up** | **Core Up DEGs** |
| --- | --- | --- | --- |
| C20ORF194  C20ORF196  MCEMP1  BTBD10  PRKAB2  LARP6  MTHFD2  FEZ1  FEZ2  TMEM126A  OPTN  ATF3  ATF1  C3AR1  PTPDC1  LURAP1L  GMDS  WLS  ATG3  CREBBP  JAG1  TSLP  SECISBP2  BAZ1B  CTAGE5  C19ORF25  KDM7A  ATF4  POP7  MUS81  CDCA8  GATA6  PDGFA  BAZ2B  PPFIBP1  ISM1  TSKU  OPLAH  FAM110A  NTMT1  RNF215  ZC3H12A  EPB41L2  ZC3H12C  LOH12CR1  PIP5K1A  SUPT20H  ATG7  DUSP5  DUSP6  JUN  DUSP1  DUSP2  FUCA2  RAB3IL1  GMIP  DUSP8  SLC7A5  CENPE  CENPF  SLC7A7  MMP14  SLC7A8  COL3A1  FXYD5  CCDC9  BHLHE40  LOC100507065  PI4KB  CCDC6  RAPGEF1  MITD1  ZNF473  LOC101927482  SLC7A1  SHB  PEAR1  ZNF48  BBC3  MED10  DLGAP1  DLGAP4  LIN54  MBD4  ATAD2  MBD6  SLF2  DGAT2  MBD1  MBD2  WNT7B  SLC39A14  NOC3L  WTIP  OAF  ADAM17  TAOK3  PPARG  NCAPD2  PPARD  SNX11  CENPA  RICTOR  GATSL3  ZSWIM6  THAP9-AS1  OLR1  ZSWIM4  SEPT10  MAP3K2  MAP1LC3B2  ATP11B  FBXO11  CDC25B  CDC25C  ATXN2L  PLOD1  DMTF1  AJUBA  PKP1  ARL8B  KCNG1  ALDH1L2  ROCK1  CRCP  ROCK2  PLOD3  CASC3  IL1RAP  EGFR  RCHY1  GTPBP10  LINC00116  SLMO1  ODF2  NEK7  HPS3  FST  FBXO32  NDC80  FBXO30  ARPC4  ZNF419  LCN2  BMP2K  SLC27A4  BCAT1  YRDC  ZNF410  INO80D  CD274  INO80E  TMEM167B  KCNE5  NAB2  FBXO28  RIF1  EIF4EBP1  CREM  PKMYT1  ARHGAP6  NHSL1  NEK2  TNFRSF9  ACTR3  ACTR2  ITGA5  BOD1  ITGA2  MUC5AC  RABGGTB  TNFRSF1B  ARL4D  NRG4  ERBB2IP  ITGA6  LINC01411  ACO1  SNAP25  DLX2  DLX3  SAR1B  SHMT2  GMPS  FAM206A  NRF1  UAP1  BICD1  MLLT4  DRAM1  PPP6R1  KIAA1524  ADAMTS18  TRIM47  SMIM15  TSTD2  FEM1C  MID1IP1  LENG1  MYO1D  MYO1B  EIF3J  SMIM20  TRIM38  NAGS  KCTD13  LINC01433  ALAS1  ECE1  LENG9  FAM208B  MALT1  PTER  LINC01426  IGF2BP2  TRIM25  PGM3  ITGB8  YAF2  ITGAV  SPDL1  FBXO8  TRIM24  MFSD12  MED13L  MIS18A  TRIM15  TRIM16  CCDC82  RIN3  CCDC68  TAGLN  HERC4  HERC3  RACGAP1  E2F1  E2F2  KDR  TCEB1  EGR1  EGR2  COX8A  EGR3  XBP1  RAB27B  SNHG17  SNHG15  TLCD1  GRB2  FBXL5  MON1B  SLC45A4  LPAR2  LDLRAD3  CCDC91  SLC9A7P1  ANPEP  IL11  AGTRAP  GAB2  VEGFA  SYN1  IL1A  IL1B  RAP2C  FAM193A  SLTM  DEPDC1  ARHGEF2  SYNC  GGNBP2  CSRNP1  DDR2  CSRNP2  RRAD  SDC4  RAP1B  MAP1LC3B  HTATIP2  PHGDH  NFIL3  PCK2  LOC101926963  IL33  SPRYD3  IL32  PNPLA8  ITCH  SH3RF3  TCEA1  TACC3  PNPLA1  DYNC1H1  NRP1  ECM1  ECM2  MAMLD1  IL26  IL24  PRICKLE2  GNAI1  HERPUD1  SH3RF1  AKAP13  AKAP12  ENPP3  TBC1D22B  MFSD2A  SPINK4  UBE2C  DENND4B  HSD3B1  HSD3B2  TMEM251  UBE2S  WDFY2  AMOTL1  TBC1D23  WDFY3  AMOTL2  UBE2M  YAP1  RELA  PRDM8  CARS  CRTC3  ZFAND5  CRTC2  UBXN2B  LOXL4  PRDM1  ZFAND3  INPPL1  NAPG  RELB  LOXL2  RELN  ASAP1  FAM196B  FGF21  DENND5B  MSX2  RELT  MORC3  YARS  IGF2R  TGIF1  BORA  NABP1  TIMM44  GEM  ADGRF4  MXD1  MKRN1  MKRN2  TCF7  LINC00941  IL7R  SRP19  PKDCC  PYROXD1  WHSC1  COX19  SLC41A1  TACO1  SLC41A2  ETS1  ETS2  ADGRG3  MUC13  PIBF1  ADGRG1  QSER1  RPP38  SUN2  URGCP  ARFGEF1  SHROOM3  NAV2  NAV3  TNIP1  SGOL1  EPN1  EPN2  TTC17  ELF1  TNIP2  PLEC  VIMP  TNFAIP6  TNFAIP3  SLC3A2  TNFAIP2  AGPAT2  C4ORF32  IRAK2  LLPH  ADGRE5  LPXN  TPBG  DAPK3  QRSL1  ERCC1  SNTB1  TBX15  GREM1  GALE  MYO15B  KRAS  PTX3  TRIB3  TRIB1  RND3  NR3C1  CBL  IL1RL1  SGOL2  OXCT2  PTK2B  CASC19  BNIP1  HUS1  EVA1A  WDR74  GTPBP2  ATG12  POLR3C  POLR3D  GTF2IRD1  CDA  NUP205  RRN3  RALB  ANKLE2  SGMS2  RALA  XYLT2  RNF6  FAM169A  APOL6  GTPBP1  KNSTRN  GNG5  TFG  ZMIZ2  FLRT1  CEP95  PXDC1  ST3GAL1  PRSS3  NAA50  PHC2  PRKCD  PLK1  PLK3  ELL2  COQ10B  EHD1  FOSL2  FOSL1  GJB2  TCN1  SNAI2  RNF216P1  CFB  PLEK2  RRP1  EPRS  SMC4  ANAPC11  ELK4  PLAC8  ZNF280C  CDK5RAP1  CSNK1A1  IL10RB  TUBB2A  FAM126B  FAM126A  NCOR1  ZEB2  MTF1  CLCF1  EXOC5  PLEKHM2  RARA  GARS  BTG3  GMEB2  GRAMD1A  PEA15  TIMP4  TIMP1  ORAOV1  TEAD1  CEP55  MBNL2  CDKN2B  MOCOS  G0S2  NR1H2  DHRS9  NFIB  LINC00869  FOSB  MAP1B  SMOC1  TOM1L1  ANGPTL4  PLIN4  PLEKHO2  PLIN2  SMG5  SMG1  CDKN1A  SETD8  LYST  NAMPT  AIFM2  ADAMTSL4  ZNF329  GPAT3  SETD2  DEDD  CSGALNACT2  RAB2A  TPM4  TPM3  IPMK  USP3  DIO2  VRK2  FAM91A1  RAB31  CNIH1  PSAT1  ZNF319  ADM2  PHIP  CNIH3  RAB1A  CSF3  CEP120  FAM129A  TNC  ABLIM1  RAB23  RAB21  ICT1  RAB24  SLC39A7  USP1  TXNL4B  ZNF787  LDLR  MB21D1  KCMF1  OSBPL6  FRMD4B  FANCE  MIR100HG  KIFC3  KIFC1  MAP3K10  ZNF777  LOC729739  RAB5A  BCL2L2  ZNF773  BRSK1  FAM49A  RAB3B  FGF2  DUSP5P1  FGF7  VPS72  PI3  C1QTNF1  SRGAP1  LINC01348  IL13RA2  COA4  RFC4  WIPI1  FAM219A  LOC103021296  DNAJC1  DNAJC7  LOC374443  LYPLAL1-AS1  BCL3  SLC26A9  MCM6  NEMP2  C8ORF76  NEMP1  TNRC6C  PLEKHF2  NUFIP2  VPS4B  UBR4  ZC3H8  RNPC3  CYR61  TNPO1  ORC2  DNAJB5  P2RY2  PHACTR4  CEP170  TNPO3  MCL1  RALGAPA1  CKAP2L  SMOX  ASCC2  CCDC59  HAUS6  FBXL13  DNAJA2  EVI2A  VAPA  EVI2B  DNAJA4  DNAJA3  SNRNP48  CSNK1G1  CCDC71L  CEBPB  TSSC4  WIPF2  CEBPG  PIK3C2A  ADAMTS6  LOC100130899  FIP1L1  POLR2C  TLK2  ADAMTS9  CCDC50  UGT1A7  IL1R1  IFRD1  AP1AR  YOD1  CTH  SIK3  UHRF1BP1  SIK1  DCP1A  LUCAT1  BCYRN1  AQP9  STXBP2  AQP3  TIAL1  PDCD10  NOCT  SUSD4  SUSD6  JAZF1  LONP1  TANGO6  COG3  CBARP  TPX2  CDCP1  SMTN  RBAK  SLC22A4  CASZ1  FAM89A  KLHDC7B  GBA2  KAT5  TNRC18  ZNF800  TP53BP2  LIF  PDCD1LG2  LHFPL2  SCNM1  P4HA2  NFX1  COPG1  TLR5  SDCCAG8  AVPI1  RAB5C  CITED2  ZFAND2A  ZFAND2B  ZFAS1  FXR2  SH3PXD2B  SPTBN1  UBXN7  TMED5  PARP6  METTL4  MPP1  SLC35G2  PARP8  PDP1  XPOT  AIDA  HEIH  SAA1  ATP6V0D2  NUPL1  F2RL3  HIP1  CSTA  PTGER2  MKI67  BEGAIN  C17ORF51  PTS  SIPA1L1  DNER  METTL5  METTL9  TBC1D10A  GABRE  CPSF2  LMO4  CPSF4  TRPV3  NUPR1  MYADM  XPR1  TYMP  ZCCHC8  ITPRIP  ZSCAN9  PRUNE2  RFNG  PVR  CWC27  LMNA  NT5E  EFNB2  RASD1  GAS2L3  HIST1H2AC  HMHA1  EIF2S2  MESDC1  DIAPH2  HGS  TMEM2  CAPRIN2  PRPF3  FCMR  PXK  RPS19BP1  SPPL2A  SEPSECS  PMAIP1  KDM4A  TIPARP  IRF2BP2  PYCR1  PYCR2  AXIN1  RBKS  DNLZ  MYH9  GPSM2  NOP2  IRF2BPL  HK2  FAM83H  FAM83G  FAM83F  HRH1  FAM83A  MYEOV  LONRF3  LONRF1  KDM6B  TCF7L1  PPP2R3A  PPP2R3C  PLAUR  AIM2  KANSL2  DDIT4  ALDH1A3  ALDH1A2  DDIT3  PFDN2  KIAA0907  HIST1H2BD  SOWAHC  HIST1H2BK  C2ORF69  HIST1H2BJ  THBS1  SLC4A5  OASL  HLX  SPRED1  PPP2R1B  MAPK6  BUB1  STARD8  ZNF282  ZNF281  DAW1  HPCAL1  RUFY2  DDX39A  PSMC4  RBP4  TMEM79  BEND3P3  NGEF  CGGBP1  USP6NL  CHIC2  ZNF274  LIN9  C4BPB  MTMR6  CLDN1  FAM132B  PSMD7  SRPX2  MICALL1  VTI1A  HES1  UBASH3B  HES4  ZNF267  C5ORF34  PROSER2  PROSER1  ZBTB7A  ABCA1  PMM1  KLF10  VPS37A  EIF1  LATS2  EIF5  SETX  EIF6  EZH2  MCPH1  PID1  IL20RB  MRPL18  FAM135A  KIAA0020  GRPEL2  LIPH  AMZ1  TMEM8A  TRMT10B  SIRT6  TMC7  RBSN  LETM2  UBAP1  PIGH  AVL9  FLII  RILPL1  IQGAP3  AREG  ATXN2  ATXN7  ATG101  SH3BGRL3  RIPK2  CFAP97  MEF2D  CSNK2A2  STAT4  F3  SH3RF3-AS1  MRPL50  TMEM56  TIPRL  AADAC  FLJ22447  CKS2  PELI1  SMAGP  ZNF217  ZNF697  SRPR  FSTL3  MYPN  PTGS2  BTAF1  FOXQ1  RABGEF1  ZNF207  IKBKG  ZNF202  SEC24A  AGTPBP1  GK  SH3GLB1  BRPF3  ZBTB17  SARS  TICAM1  S100A7  SLC25A33  SEC24D  S100A9  S100A8  PVRL1  MAD2L2  OSBP2  PPP1R15B  TMF1  PPP1R15A  UBE2D2  ZBTB21  HR  SLC9A1  RNFT1  BCL2L12  SH3BP1  SH3BP4  SLC16A7  FLNB  YKT6  PITX1  SLC16A3  KPNA4  SH2B3  SLC16A6  C11ORF86  CPA4  SLC38A2  KIRREL  SLC38A1  GOT1  FTSJ1  TNFRSF10B  FOXN2  NFKB1  LRRC42  NFKB2  MFHAS1  FRMD6  STK24  NT5C3A  PIGBOS1  MLXIPL  CDC42EP3  LINC01239  BIRC5  CDC42EP2  TROAP  MDM4  CDC42EP1  BIRC2  R3HCC1L  BIRC3  MDM2  LRRC59  GPR87  RORB  CYP19A1  GLIS2  STK10  NIPBL  GLIPR1  SESN2  NELFE  GAREM  PIM3  ROS1  PPP1R12A  GFPT1  KSR1  GFPT2  RAD23A  NPC1  FOXL1  HAX1  AAK1  NFE2L2  FBLIM1  RASGRF2  FOXK2  ME1  DSP  DST  ZBTB38  ARNT  FOXJ2  PPM1B  CCAT1  DBF4  P4HB  PABPC1L  MAK16  LINC00341  NXT1  ARHGAP11A  PPM1M  COL15A1  FURIN  ZBTB43  SQRDL  MFF  ZBTB41  PPP1R13L  LRRC8D  LRRC8B  LRRC8C  UPP1  SLC25A25  CCNJL  SLC12A4  SFMBT1  EREG  MAPK13  FNDC4  ZNF714  RTCA  MXRA7  CCM2  ZNF710  EPHA2  CALM2  EPHA4  TMEM87A  STX16  CCNK  CCNJ  SATB2  CCNH  PPP1R3B  BAIAP2L1  EXOSC8  NCLN  RAB11FIP1  RAB11FIP2  ZNF706  DHX38  PAQR5  EAF1  XRCC3  PRG4  SNW1  C7ORF43  PPP1R1C  INTS6  PPP1R18  GCC1  IFI35  THUMPD2  GCC2  TMPO  RASSF1  PELO  PCBP4  RASSF6  MIER3  MICAL2  HIVEP2  KIF21A  HIVEP1  KIF21B  CEP170B  SCN1B  CCR1  NCOA3  AKIRIN2  WARS  FOXD1  NCOA6  TNFRSF12A  NCOA7  ENAH  NR5A2  PTPRF  ANXA2P2  AARS  INTS12  DOT1L  KIF14  BRCA2  SLC7A11  PTPRG  PTPRH  SBNO2  SOCS2  ERI1  CTSL  STK39  SNX9  PDRG1  ZFR  GCH1  DOHH  STAMBPL1  PER2  STK17B  PER1  NEDD4  FOXD2-AS1  ERRFI1  CD63  SLC43A3  AKR1B1  HEATR1  NUBP1  DERL1  SPTLC3  RSRC2  EP300  SERTAD1  REV3L  CD55  SCARA5  MOK  ERN1  TANC2  COL5A3  CD68  DCLRE1C  INPP1  CD82  AURKA  SLC1A4  SLC1A5  ATAD2B  NDRG1  AURKB  CLGN  SH3TC1  HSF1  CCRL2  PPFIA1  MICA  PRNP  CCDC137  ARHGEF18  BUD31  INHBA  DDX55  DDX52  KIF23  SLC30A1  INHBE  SSH1  CREB1  H1F0  FAM175B  KIF2C  TAB2  CCDC130  GCLM  CREB5  CCDC124  MIDN  LRRFIP1  WWC1  WWC2  GXYLT1  PSEN1  MYDGF  PRND  CCNB2  CCNB1  CYP11A1  NDEL1  CAMTA2  SH3GL1  VDR  RBM15  MMP1  MMP2  GALNT3  SLC52A2  HMGA1  CLK3  CLK1  IRF1  NUP50  IL6  CLCN5  SCN4A  RBM24  TMEM45A  TMEM45B  PSMD11  MAST4  BCL2A1  LAMC1  FUT1  LAMC2  AFF4  CHD2  FUT4  PDLIM4  BCL7B  H1FX  APOPT1  PDLIM7  ATP2B4  LMTK2  GLRX2  THAP1  ATP2B1  SOD2  TAF9  DAAM1  CCDC174  DDX27  GOSR1  LAMA3  GOSR2  RPGR  CLMP  CLMN  EPS8L3  RBM33  RBM34  MED1  ACOT9  UNC13A  WSB2  SERPIND1  UPF3B  MED8  DYNC1LI1  VIM  ASF1A  NHLRC2  CPEB4  CDK16  PCED1B-AS1  CDK17  EHF  MAML2  SERPINE1  SLAIN2  BACH1  GABPB1  ARIH2OS  MILR1  CREB3L2  CREB3L3  CHST11  RPLP0P2  CYP1B1  S100A16  S100A11  DNAH14  ST14  ZNF140  CIDECP  FHDC1  SLC6A9  HOXB2  HOXB9  PRR13  OGDH  PTP4A1  CDC42SE2  CDC42SE1  C9ORF72  NEDD4L  MYSM1  ABCA13  GLDN  ZXDB  PPP3R1  ADNP2  IER2  PNO1  IER5  JUNB  IER3  SLC19A2  ANXA1  ANXA2  ITPK1  CCL20  MAD2L1BP  TCF12  BRAF  PAWR  RNF146  RNF149  JDP2  ZNF598  SKA1  C9ORF91  ZNF593  ZNF592  ABHD5  ELL  SKA3  C16ORF52  TRIM8  CSRP1  ZNF106  BRD4  SERPINB5  SERPINB2  IFNGR1  SERPINB1  GPR1  AEN  LOC101927374  SERPINB8  MZT1  TMEM154  TMEM156  RYBP  RNF126  CLIP1  RLIM  HBEGF  C16ORF87  PLCB1  PHLPP2  HSPA13  PRSS22  C16ORF72  GPR132  NUF2  RIOK3  ZNF569  ZNF562  AOC2  PHLDA1  AOC3  SEMA4C  GPT2  GTF2F2  PNRC1  RALGAPB  MELK  STIM1  CNOT3  ZYX  DGKH  PHLDB2  UHRF1BP1L  GPS2  DGKB  HJURP  AHR  GTF2E2  BEST1  TUBB6  ZFP36  NACC1  RNF19B  CDH1  C1ORF109  GRB10  MISP  ZUFSP  PHLDA3  WHSC1L1  IGFBP1  ESRRA  PNISR  BTN2A1  KREMEN1  LSM4  RCAN1  MAFF  RNF168  MOB1B  METTL23  MAFG  ERF  SPIRE1  IARS  MAFK  XAF1  CEMIP  NOTCH1  CALCOCO2  NOTCH2  IRS2  ANKRD12  AK6  VTN  BAG2  BTBD3  ABL2  NDP  ANKRD11  EFHD2  ABL1  SEMA7A  TAPT1  REXO1  C18ORF8  ASNS  CFLAR  AIM1L  GPX8  PTPN12  MARCH7  REXO4  MARCH3  CDK8  CDK6  ZNF516  NF1  SERINC2  SOS2  TRMT61A  NUMB  NFAT5  PTPN21  PTPN22  TARS  KLC2  DUSP16  KLC1  C15ORF48  PTBP2  RREB1  SKIL  WASF2  PGPEP1  PATL1  SPAG9  RNF24  ADRM1  KLF5  KLF4  ZFP91  KLF3  KLF6  PAPD7  BOLA3  SMPDL3A  RP9  CBLL1  SEMA3A  LUZP1  SART1  NHS  ACSL1  RUNX1  ACSL5  ACSL4  JMJD6  RUNX2  LACC1  TMCC3  EIF5A2  ANTXR2  ARNTL  AKAP2  RGAG1  CHAC1  TIPIN  NUS1  SNHG1  SIAH1  IL17RA  GADD45B  GADD45A  CMC2  PAPPA  AMMECR1  STAM  C8ORF4  SNHG8  NUAK1  ARFGAP3  CEP57L1  MEIS3P1  EZR  PTGES  CREBRF  GOLGA4  GOLGA5  RUSC2  CMYA5  GOLGA2  AGFG1  SOX5  CDT1  TSC22D2  TSC22D1  TSC22D3  NMI  RHOF  RHOG  TUBE1  IGFN1  KRT10  RHOB  RHOC  POC5  TFF2  RHOU  CC2D1B  SGK1  IGSF8  CLIC4  NUTM2B-AS1  C10ORF88  STC2  VLDLR  HINT3  PCF11  ZMYM5  PRKCDBP  GNPNAT1  TFE3  MAP7  ANKRD1  ARNT2  CDV3  TRAIP  RANBP9  TJP1  DLC1  B3GNT5  MARS  WAC  H2AX  TOMM40  GDI1  FAF1  TOR4A  CILP2  MAP4K4  TGFB3  GDF15  ZFC3H1  ARID3A  PPP4R3B  ARID3B  PPP4R3A  CGRRF1  TMEM133  TBL1XR1  ARHGAP31  EOGT  RELL1  ULBP1  STAM2  RAI14  RGS17  SLFN5  RGS16  ZFP36L1  ARHGAP42  MYH15  EP400  REPS1  ATP6V0C  PSAPL1  DESI2  STIL  ARID5B  CDYL  SDCBP  CKAP5  ACBD3  DOCK6  AKNA  DOCK5  GOLT1B  SH3KBP1  SLC2A1  SLC2A3  LTBP1  SEL1L3  SLC2A6  NR4A1  PLCXD1  POGZ  PDZD2  DTWD1  COBLL1  MARK3  PSPH  HSPA9  IFNLR1  ARAP2  NFKBIB  DDX19B  MAN2B2  NFKBIE  CXCL8  B4GALT4  ESRG  TSEN15  CKS1B  DTX2  CXCL2  LINC01094  CXCL1  NID2  CXCL3  CXCL6  MOSPD1  U2AF1L4  STK4  DPP4  CXCL5  SGPL1  DPP9  BDKRB1  AOX1  BDKRB2  HIST1H1C  SPEN  PDE4DIP  PDE4D  LCA5  KCTD5  KCTD9  FMNL2  PALLD  LTBR  FMNL1  LRIF1  PALM2  PKD1  KRT80  TUFT1  USP12  PRRG4  KYNU  LRIG2  NIFK | MED1  STARD4  UNC13A  HOMER2  GATAD2A  DAW1  PRKAB2  C12ORF75  GTPBP4  L1CAM  COL1A1  MEX3C  DEPDC1B  RBP4  FDX1  TBPL1  C19ORF48  NGEF  CDK17  ATF1  PON1  RALA  COL12A1  PON2  NCAPG  ADARB1  FAM169A  KNSTRN  ZMIZ2  GNG5  CYP1B1  S100A16  UBASH3B  WLS  HES4  PRSS3  EPSTI1  DNAH14  NAA50  PRR11  PRKCI  PRPF38B  C12ORF56  EIF1  CCSAP  FOSL2  GIGYF1  FOSL1  EHD4  EIF5  FHDC1  LGMN  NOX5  RAD18  SNAI1  HOXB9  CHST6  ATF4  PTP4A1  POP7  PID1  FLG  CDCA2  CTBP2  PLEK2  CDCA8  GATA6  SMC6  PDGFA  SMC4  ABCA13  FAM135A  ISM1  NTMT1  GJA1  PLAC8  GRPEL2  EPB41L2  ZC3H12C  ZNF280C  PIP5K1A  IER2  MYBL2  CEP78  SLC19A3  PPAPDC1A  BARD1  CCL26  CENPV  ANXA3  ITPK1  CCL20  SPHK1  LYZ  DCBLD1  PAWR  DUSP8  CENPE  LETM1  SLC7A7  ZEB2  SLC7A8  CENPK  BHLHE40  LOC100507065  EXOC5  ADGRL2  C5ORF51  MGLL  SKA1  ZNF593  YTHDC2  RILPL1  IQGAP3  AREG  TIMM10  PEAR1  SKA3  ACAT2  CSRP1  RRAS2  TIMP1  RIPK2  TEAD1  CEP55  TEAD4  BRD4  ATAD2  MBNL2  WTIP  NFIA  SERPINB8  F3  AGMAT  DHRS9  EBNA1BP2  MRPL50  FOSB  AADAC  FLJ22447  ANGPTL4  PELI1  SMAGP  GOPC  STAG3L4  PHLPP2  GLI2  CENPA  FSTL3  GLI1  PTGS2  FAM102B  GPR132  NUF2  ADRBK2  SBDSP1  ZNF569  ZSWIM6  THAP9-AS1  ZNF202  CSGALNACT2  SEC24A  CSGALNACT1  GK  PRRX2  DIO2  VRK1  CDC25A  VRK2  APOA1BP  CDC25B  CDC25C  MELK  RAB32  SRSF4  SLC25A33  S100A9  KCNG1  S100A8  MAD2L2  PLXND1  DGKB  UBE2D2  HJURP  TNC  HR  IL1RAP  METRNL  HOTAIRM1  SLC9A1  TSPAN33  RNFT1  SBSN  ABLIM1  TUBB6  ZFP36  BCL2L12  ICT1  HAS2  C1ORF109  USP1  MACROD1  PITX1  GRB10  SLC39A8  LZTS3  PHLDA3  KPNA4  HMGN1  KCMF1  SH2B2  C11ORF86  IGFBP2  PNISR  SLC38A1  OSBPL6  ODF2  FRMD4B  FOXN2  SLC9A3R1  NDC80  LOC100128317  MAFF  NT5C3B  FBXO30  STK24  NT5C3A  TRAF5  KIFC1  KIAA1755  BIRC5  TROAP  MDM4  BMP2K  LOC729739  BCAT1  BIRC2  MDM2  LRRC59  NOTCH3  RAB3D  NOTCH2  INO80E  IRS2  RIF1  EIF4EBP1  RORB  PKMYT1  ARHGAP6  WISP1  CYP19A1  C6ORF223  FGF7  BAG2  NDP  PI3  NEK2  ROS1  EDIL3  IL13RA2  SLC2A4RG  RFC4  GFPT1  KSR1  DNMT3B  WIPI1  FAM207A  YDJC  PTPN12  FAM219A  NPC1  FOXL1  ARL4D  DNAJC1  NRG4  RTN4RL2  BCL3  SLC26A9  ZNF516  CDR2L  PSMG3  TOP1  ARL4A  MCU  DLX2  GTF2A2  DLX3  GMPS  UAP1  FAM109A  DUSP16  CCDC85C  KIAA1524  DNAJB5  RDH10  P2RY2  WASF3  MCL1  SMIM15  PPP1R14A  DSP  PPP1R14B  PLEKHG2  UNC5B  CKAP2L  SMOX  CCDC59  SPAG5  KIAA1549L  KLF5  PTBP3  MYO1E  KPNB1  CCAT1  DBF4  RAB15  MYO1B  SH3D21  DNAJA4  PAPD7  TSPAN18  SNRPA1  SLC25A10  CSNK1G3  SLC25A15  CCDC71L  CEBPB  NXT1  ARHGAP11A  TRIM31  WDR4  PRR5L  MFF  USH1C  ADAMTS6  LRRC8D  IGF2BP2  SPDL1  UPP1  IFRD1  DDIAS  AP1AR  ACSL4  TM4SF5  GCAT  RUNX2  MIS18A  SLPI  ZNF714  TRIM15  ANP32E  ZNF711  EIF1B  EIF5A2  CCDC68  BCYRN1  CCNH  CCDC102B  TIAL1  STRIP2  RP9P  PDCD10  RAB11FIP2  CDC27  CPNE2  KDR  APBB2  TIPIN  SRGAP2B  FER1L4  EGR2  GADD45A  XRCC3  CMC2  COMMD3  PAPPA  RASSF9  RAB27B  C8ORF4  PRG4  SNHG3  SNHG17  SNHG15  TPX2  KLHDC8A  BANK1  STAR  SDC2  NOL12  PTGES  RCOR2  RBAK  SPON2  SPARC  ANP32B  C9ORF142  CASZ1  FAM89A  ECI2  GFI1  DCUN1D5  LPAR2  KLHDC7B  TMPO  DMBX1  SLC9A7P1  MICAL2  PPAT  KIF21B  SOX7  AGFG1  CDR2  AKIRIN2  STRADB  GAB2  NME4  RHOB  VEGFA  RHOC  SYN1  ENAH  IL1A  NR5A2  PTPRF  CMSS1  P4HA2  ID1  DEPDC1  ALDOC  SGK1  AHCTF1  MRPS15  ZFAND2A  ZFAS1  KIF15  PCSK9  VLDLR  PTPRH  SNX5  ERI1  HLTF  C1QBP  HNF4A  GNPNAT1  MAP7  NFIL3  LYAR  LOC101926963  IL33  CDV3  GCH1  RANBP1  SUSD2  GAL  SH3RF3  GPAM  XPOT  STK17B  NEDD4  TACC3  NUPL1  CD44  POPDC3  H2AX  NRP2  ECM1  MAMLD1  SLC43A3  HEATR1  SLC20A1  MKI67  BEGAIN  C17ORF51  PTS  VMO1  SCML1  SIPA1L2  NPW  CILP2  RSRC2  DNER  SERTAD1  METTL9  MFSD2A  DCPS  CD55  SPINK5  TFAP2E  DIMT1  OCRL  ARID3A  KAZN  LMO7  PPP4R3A  TANC2  DOK4  DKC1  UBE2S  KIF4A  COL5A3  UBE2M  YAP1  INPP1  CRTC3  CHRNA4  AURKA  CHRNA9  PVR  ATP1A3  CWC27  LOXL2  HOXA11  TIMM50  EFNB2  RELN  RASD1  ASAP1  FAM196B  PTTG1  S1PR2  BANF1  MICA  MORC3  AASDHPPT  AZI2  FDPS  MGAM  TGIF1  ACBD6  RPP40  DESI2  STIL  ARID5A  INHBA  TIMM44  MESDC1  CKAP5  SYT7  LOX  CAPRIN2  TCF7  KIF2C  TCF4  TCF3  GCLM  CREB5  LINC00941  MIDN  WWC1  ARHGEF26  SH3KBP1  GXYLT1  SLC2A3  NTS  NR4A1  PLCXD1  CAMSAP3  PRND  CCNB2  PDZD4  NME1-NME2  ADGRG3  CCNB1  RGS2  CYP11A1  DTWD1  COBLL1  MUC13  NDEL1  SACS  MARK3  HSPA9  SRGN  HMGA2  HMGA1  TNIP1  SGOL1  GPCPD1  EPN1  ACVR2A  IL6  CLCN5  IRF3  MAP6D1  TMEM45A  SMIM3  NID1  NOP2  TNFAIP6  CKS1B  TRPS1  CXCL2  CXCL1  AFF4  CXCL3  CHD1  DPP4  CXCL5  FAM83H  FAM83F  CYP26B1  SNAPC2  IRAK3  BDKRB1  FAM83A  AOX1  BDKRB2  CA9  LONRF1  PDLIM7  FGB  RRM2  SPEN  TCF7L1  FGG  MIIP  PDE4D  DDX10  GLRX2  SSRP1  ANXA10  PPP2R3B  ATP2B1  DCN  AIM1  DDIT4  ALDH1A3  LRIF1  PFDN2  S100P  ZP1  PTX3  ZPR1  SIGIRR  RBM28  RBM26  SOWAHC  C2ORF69  DDX21  THBS1  DMKN  SLC4A5  CBL  SH3YL1  IL1RL1  RPS6KA6  SGOL2  CLMP  OXCT1  OXCT2  POLD2  EPS8L3  NIFK  IFT57  RBM6 | TEX30  WDR83OS  CKMT1A  C12ORF75  NECAP1  NECAP2  FEZ1  C19ORF43  MTHFD1  C19ORF48  ORAI1  SKAP1  HCP5  C3AR1  PON2  TPRG1L  TFPT  C19ORF53  WLS  TOE1  HLA-DRB5  ATP6AP1  PBX3  TBL2  JTB  PKIB  C19ORF25  CHST2  SFXN2  C19ORF24  CHST3  ATF5  HLA-DRB1  TSTA3  TARSL2  MEPCE  TFRC  CDCA5  POP5  POP4  LYPD3  CIAO1  HTR7  NKX3-1  ATG7  GSTM4  GSTM3  JUN  GSTM1  DUSP1  FUCA2  JOSD2  TOMM40L  MMP15  MMP14  SLC7A8  FXYD5  C12ORF29  CENPM  CENPN  CENPP  CCDC3  ERGIC2  ERGIC1  ELOF1  C11ORF1  NOC4L  LOC101927482  MED17  ECHS1  ATIC  MED10  SLC18A2  DGAT2  GSTO1  MBD3  KIAA2013  CNPY2  RABIF  COX6B1  SNX17  ADAM15  SDHAF2  CINP  PTRH1  TSR3  TSR2  GSTP1  PRKAG1  SMPD1  TSPO  BID  NUDT14  FBXW5  FAM118B  CDC25A  CORO2B  BFSP1  HNRNPAB  APOA1BP  PLOD1  NAAA  UQCRQ  LY6E  CDKN2AIPNL  PKP1  ARL8B  AGRN  KCNG3  ODC1  PDCD6  DLST  MPV17L2  APEH  SLC9A3R2  SPP1  STX3  LINC01444  NOP16  UROS  RFXANK  ARPC5  MED27  MINPP1  ARPC4  DRAP1  ARPC2  FTL  YRDC  ZNF410  INO80C  COL17A1  OLFML2A  EIF5A  TMEM255B  ITGA3  FAM207A  FUZ  AHSA1  TMBIM1  ARL4C  ADORA2B  RN7SK  TMBIM4  ETF1  KCNK6  KHDRBS3  UFD1L  SAR1B  SPR  CCDC85B  C6ORF203  SNAP29  CRSP8P  SMIM10L1  NFKBIL1  NAGA  EXOG  NAGK  MYO1D  EIF3K  STUB1  GAPDH  NOP56  TRIM35  MTCH2  ALAS1  ECE2  ETFB  SRM  FOXRED2  CYSRT1  PRADC1  PGLS  FBXO4  ITGB8  RNMTL1  ATP6V1E1  MFSD12  CDC20  PEX16  MT1A  CRISPLD2  MT1E  MT1F  RIN1  CERK  SSNA1  HM13  IL27RA  CPNE2  KDR  TCEB3  RILP  E2F7  RTFDC1  COX8A  ALG6  ALG3  TALDO1  ALG1  AIMP2  MSRA  RNF144A  TLCD1  CSNK2B  NDUFAF6  NDUFAF3  SDC1  NDUFAF4  MON1A  SUGP1  APOBEC3B  ATP6V1B2  ZMYND19  ECI1  ICAM3  ZBTB2  MT1X  LPAR3  RCBTB2  GSPT2  MT2A  LGALS3  NUDCD3  LGALS1  ANPEP  EPC1  KCNN4  NKAIN1  IL11  SLC10A3  AGTRAP  VEGFA  IL1A  IL1B  RAP2B  UFC1  CALR  SDC3  MRPS12  SDC4  MRPS11  ACTR1A  NEU1  HTATIP2  ATP6V1G1  SPRYD4  MRPS26  SPRYD7  APRT  TPST2  BRMS1  FAM195A  CARS2  PNPLA1  DHRS13  NRP1  COX5A  SYNM  GBA  MRPS33  NUDT1  PRICKLE2  CAPG  RAB22A  COX5B  NANS  ENPP1  SDF2  XDH  RAC2  MFSD2A  HSD3B7  UBE2I  UBE2C  UBE2F  AKR7A2  EEF1A1  BOP1  UBE2Z  SPOCD1  UBE2S  UBE2T  GNG12-AS1  POLE3  GPRIN1  POLE4  PPIF  RPP25  DEGS1  NAPA  NDUFB10  EEFSEC  TTC38  DENND2D  CHCHD10  GEM  ADGRF4  MXD1  UBA1  MICU2  FBN1  LINC00941  IL7R  TXN2  UBIAD1  ACTG1  HTRA1  ACTG2  HIGD1A  PTRF  COX14  IL6R  ADGRG1  URGCP  GALK1  POLA2  EEF1D  ACOX3  TNIP2  ACOX2  ACOX1  RER1  CUEDC2  IFT46  VIMP  CHI3L1  TNFAIP6  DHRS7B  SLC3A2  AGPAT2  PLD3  TMEM201  SHARPIN  LASP1  IRAK2  LLPH  IRAK1  TMEM208  LPXN  FAHD1  TIMM8B  RRM2  EYA2  RRM1  ERCC1  C19ORF60  TBX15  GREM1  GALE  PTX3  OTUB2  RND3  USE1  C19ORF70  GAMT  PRELID1  HOMER3  AP1S1  UBE2MP1  KIAA1462  EVA1A  HNRNPM  OSTM1  POLR3C  TOLLIP  GLA  VPS25  POLR3K  CDA  PI4K2A  RALB  C14ORF142  ARPC5L  ADARB1  GPATCH4  FLRT2  RRP36  ST3GAL1  RALY  C2ORF82  VPS18  ICMT  NDFIP2  PRKCD  MCAM  RMDN3  EHD1  FOSL1  GJB2  ITPA  FARSA  DANCR  PLEK2  RRP1  WDR46  LRPAP1  ANAPC11  C14ORF169  WDR45  BLOC1S2  MLST8  CDK5RAP2  HMOX1  FLJ23867  TIMM17B  HMOX2  DHRS4L2  TPI1  TUBB2A  NSA2  ADI1  PQLC2  ARPC1A  ARPC1B  PEA15  WDR20  SDSL  DHRSX  PBK  TIMP4  TIMP1  RRS1  MGMT  MOCOS  G0S2  NR1H2  ELP6  AGMAT  TUBB4B  DHRS4  FDX1L  MAP1A  DHRS9  HADHA  MAP1B  AHRR  ANGPTL4  NUCB1  PLP2  NDUFA12  CDKN1A  NDUFA13  PYGB  NDUFA10  NDUFA11  RARS  FAM127C  FCGRT  GPNMB  PBDC1  RRP12  GPAT4  TMEM38A  GPAT3  RBPMS2  YIPF1  TPM3  GSS  PPP1CA  COPRS  RAB31  CNIH1  RAB32  COPS6  RAB38  COPS3  STAMBP  CSPG5  SNRNP25  TCTA  SNRPE  FKBP4  COPS8  TCEAL1  FAM129A  ABLIM3  ICT1  LAMP2  DDRGK1  TGFBR3L  CFL1  ZNF787  SIAE  HMGN2  TRAPPC3  TRAPPC4  MAP2K2  TRAPPC1  EIF2B3  OSBPL6  TRAPPC5  GTF3A  FRMD4A  REEP5  PGP  TPP1  MIR31HG  N6AMT1  ZNF771  SLC38A10  HSPBP1  KRBOX4  MPC1  MPC2  PI3  PLXNA2  RAB4A  IL13RA2  COA3  RFC2  ANGPT1  LANCL2  EDF1  WIPI1  DNAJC1  MTFR2  DNAJC9  MIR3682  ORC6  DNAJB2  P2RY8  PLTP  RRP7A  ORC3  PLEKHG4  FKRP  CPT1A  IDH3G  ASCC2  HAUS3  PKM  DNAJA1  P2RX4  TST  DAD1  PPAP2C  DNAJA4  RAB13  PPAP2B  DRG1  POLR1E  TSSC4  ADPGK  KIAA0101  BPHL  TPRN  THBD  POLR2C  POLR2F  POLR2J  POLR2K  CCDC51  IDH3A  ILKAP  POLR2L  ADAMTS8  UGT1A7  CD320  ESCO2  P3H3  P3H4  MAPK8IP2  MTHFD2L  GSAP  TLN2  ATP6V0D1  SLC35B2  CISD3  HS6ST3  PNP  SUSD6  TMED9  SLC35C1  TANGO2  HLA-C  HLA-A  HLA-B  VASN  HLA-E  WBSCR16  SMTN  IFI27L2  SLC22A4  ISCA1  FAM89B  SLC35D1  PITPNA  KIAA0930  FTH1  RPS10  SIL1  RPS12  ATP5SL  STRADB  SLC35E4  TUBB  MLF2  HIST3H2A  COPG1  RAB7A  RAB7B  COL18A1  GK3P  RAB5C  DTYMK  SLC35F6  ZCCHC17  FXR2  SH3PXD2A  DGCR6L  TMED3  TMED1  ATP6V0E1  CBR1  ATP6V0E2  METTL1  SUSD1  MPP1  QPCT  ZDHHC12  SLC35G2  ZDHHC18  ZDHHC16  ATP6V0D2  NUPL2  CSTB  LINC01272  ZDHHC24  CST7  PTS  CST3  SIPA1L2  DNER  METTL8  METTL9  UBTD1  GINS3  PHYH  SLC31A2  UQCC3  UQCC1  NOSIP  SHISA4  ACYP2  ATP1B3  CDK2AP1  CHMP4A  MPHOSPH6  ALDH16A1  TYMP  DAP3  ITPRIP  ZWILCH  SEPW1  COMT  CNDP2  LMNA  NT5E  EMC3  BMPER  TRAPPC12  EMC7  HIST1H2AC  C17ORF89  TRAPPC2L  IDH2  PRPF6  TMEM5  TMEM2  COL6A1  SPECC1L  DYNLRB1  FGFR1  THOP1  SELT  RASEF  RPS19BP1  ZMAT2  COL4A3BP  UBB  MLKL  HADH  TIPARP  C14ORF119  SRGN  PARVB  PARVA  GRHPR  COQ4  DAP  RABEP2  RFT1  DNLZ  SMKR1  TERF2IP  COPE  AVEN  STOML1  RPAP2  EMG1  H2AFJ  MFSD5  MFSD6  LONRF2  ATRIP  H2AFZ  ANXA10  PLAUR  MLPH  GAP43  RPS6KB2  ALDH1A3  PITHD1  PFDN1  HIST1H2BC  HIST1H2BD  HIST1H2BK  HIST1H2BJ  RABEPK  OASL  HLX  GNA11  RPS6KA2  BUB3  POC1A  B3GALNT1  STARD7  STARD8  APCDD1L  C20ORF27  HPCAL1  EML1  HSPE1  SCAMP4  SCAMP3  MYL6  DDX39A  PSMC4  RBP4  PSMC5  PSMC2  ETHE1  PSMC3  UQCRC1  RPL22L1  RETSAT  NCAPG  PSMD9  LSM10  SRPX2  PSMD8  KRTCAP2  PREX1  PSMD4  DNTTIP1  VTI1B  PSMD2  AP2S1  RABAC1  NGFR  SIGMAR1  DNTTIP2  SNF8  EIF1  FERMT1  EIF6  UCK1  PRKAR1B  TMEM9B  VAMP3  PID1  HACD1  TTYH2  UXS1  MRPL17  TMEM97  MRPL15  MRPL12  MRPL20  HSPH1  MRPL21  FAM134B  NETO2  GRPEL1  ZBTB7C  PUSL1  CYB5R1  IKBIP  FGFBP3  NSUN5  EMP3  MRPL27  MRPL24  SUMF1  FAHD2A  SIRT2  MZT2B  GPRC5C  TMEM106C  PIGH  MBLAC1  C11ORF31  FPR1  MRPL37  AREG  PQBP1  MRPL36  MRPL34  TRMT112  AACS  MRPL41  ATG101  FLVCR2  SH3BGRL3  RSU1  RIPK1  C11ORF24  HSD11B1  DYNLL1  UTP6  ARG2  MANEAL  DYNLL2  F5  FAM225A  PSMA6  PSMA5  SPSB1  FABP5  BDH1  AADAC  CKS2  HLA-DPB1  SMAGP  SLC29A1  ZNF697  PIGT  LAD1  HEXB  TMEM54  MRPL55  PTGS2  PSMB7  PSMB8  TUBA1C  HLA-DMB  PSMB5  PSMB4  UCN2  EFCAB2  HLA-DPA1  HLA-DMA  GK  AKR1E2  SRD5A3  MUL1  TUBA4A  OLFM1  MRTO4  SLC25A39  SLC25A38  HAT1  MAPKAPK3  S100A6  SLC25A33  NPTXR  SQSTM1  S100A9  RNASEH2C  UBE2D4  PPP1R15A  NTPCR  SLC9A2  CARD8-AS1  BCL2L12  HARS  SAP18  HAS3  YKT6  ZNF668  SH2B3  SLC16A6  KPNA2  LRRC47  GOT1  FTSJ3  TMEM18  TMEM11  BIRC5  RNASEH2A  ST6GALNAC4  PIN1  LRRC59  TUBGCP2  ARL2  GPR87  RORB  PORCN  GLIPR1  THEM4  NELFE  SRD5A1  SLC38A5  GFPT2  APCDD1L-AS1  NDUFC2  BAIAP2  NPC2  CDR2L  PPP1R12C  PSMG1  L3MBTL2  SLC25A4  NDUFB7  GTF3C6  NDUFB2  C6ORF1  PMVK  ATP5H  DSEL  ME2  ME3  ZNF622  PPP1R14A  PPP1R14B  NDUFA8  NDUFA6  LRRC8A  MBOAT7  NDUFA2  LYSMD2  RPL23A  MCOLN2  TPD52L2  ATOX1  P4HB  ANAPC5  DTL  SLC25A11  KDELR3  SLC25A10  PCOLCE2  NRROS  EPHB6  DRAXIN  NXT1  SLC47A1  FHL2  SQRDL  CXXC5  PPM1G  SLC25A29  SUMO2  SUMO3  SLC25A20  LRRC8E  PCBD2  UPP1  AAMP  NOMO1  PCBD1  EPHB2  SLC25A25  ENTPD3  IMP3  IFITM10  SLC12A2  EREG  GCAT  DCLK1  FNDC4  CD2BP2  RTCA  MXRA7  CCM2  PREB  CALM1  KDELR2  KEAP1  PLEKHB2  UBE2L3  STX12  UXT  NCLN  EXOSC4  AAR2  ARSB  SURF2  SURF1  PAQR5  SURF4  XRCC1  COMMD4  COMMD1  PRG4  COMMD7  RPE65  INTS5  TAGLN3  INTS7  LOC100507600  LRR1  TXNDC12  PTTG1IP  MRPL1  IFI30  PELO  C7ORF50  MRPL9  PCBP4  RASSF5  MICAL2  ZBED1  TSPAN3  SCN1B  CTSA  FOXD4  AKIRIN2  SSSCA1  TNFRSF12A  NCOA5  UBE2G2  MRPS6  ID2  ANXA2P2  CRELD2  FOXC1  SUV39H1  SWI5  SUPT4H1  GPAA1  NEDD8  PCSK6  AATF  IDS  KIF1C  CTSD  CTSC  PDRG1  FAM58A  CCT3  BCAP31  DBNL  GCH1  RPA2  UBE2E3  CREG1  DOHH  GPAM  STK17B  CCNE1  CD47  CD44  FKBP10  CD63  SLC43A3  FAM57A  NUBP1  MALSU1  PEPD  NUBP2  SAMM50  SPTLC3  HYAL2  CAMK2N2  ENC1  NCS1  SERTAD1  BCAP29  CD59  CCT7  CD74  PDGFRA  G6PD  ATAD3A  ATAD3B  KLHL21  DKC1  FAM173B  TMX2  SSU72  COL5A3  KIF4A  ACTL10  CD68  RPL26L1  MPG  CD82  GEMIN4  SLC1A4  KLHL35  SLC1A5  FUOM  PAK1  CCND3  GPC1  PERP  PES1  CCRL2  CAPN2  NDUFV3  MICA  NDUFV2  LYNX1  ADPRHL2  OST4  PRNP  CCDC137  DDX56  INHBA  CYBA  CREB3  QDPR  CCNA2  PROCR  STT3A  KIF2C  GCLM  VCP  CCNB1  EBP  FAM174B  FAM174A  ACAA1  VDR  MMP1  GALNT3  MMP3  HMGA2  LRWD1  SLC52A2  RCN3  NDUFS8  ARL2BP  FAM96B  NDUFS3  CYCS  ATG4B  QRICH2  MCUR1  DDX49  YWHAB  BCL2A1  LAMC2  PDLIM4  BCL7B  BCL7C  TAF15  HSDL2  NAA10  TAF12  TAF10  ATP2B4  GLRX2  ATP2B1  EEPD1  PLA2G15  PPDPF  CPTP  TAF9  FAM50A  CHURC1  ZP3  MEA1  DCTN3  DDX24  GOSR2  EMC10  ATP2A3  CLN6  CYGB  SFR1  SCN5A  DLEU1  SCARB1  SERPIND1  VIM  HSPA1A  CYP1A1  RPL28  ACOT7  ASF1B  ASB2  SF3B5  SF3B4  RPL13  SERPINE2  TPGS1  ASAH1  CHST12  CHST10  CHST11  RPL15  S100A13  CYP1B1  S100A16  CSRP2BP  S100A11  FZD1  F8A1  ST13  FN3KRP  NUDT16L1  TMEM199  FIBP  FHDC1  DUS1L  C9ORF78  ABCA13  RPL9  VPS26A  PRDX6  GPR176  PRDX5  PTPMT1  PRDX2  PRDX1  RPL36  PPAPDC1A  GSDMD  NUDC  BLCAP  ANXA2  H3F3C  ANXA5  H3F3B  SSR4  TIMM23  TIMM22  CDC6  FAM102A  RNF149  C16ORF62  ITPKA  TRMT6  SKA1  ZNF593  RPN1  ENDOD1  CLTB  CLTA  ADO  CYB561D2  TMEM164  SKA3  CSRP1  PNPO  LAPTM5  TM9SF1  TMEM158  KIAA1279  NMNAT1  HYLS1  LAMTOR1  HBEGF  TCF25  LPCAT1  TMEM141  TMEM147  IMPA2  ENSA  MPV17  AGA  LDLRAP1  METTL7B  TMEM14C  TOR1A  TOR1B  RPS7  C1ORF122  GTF2F2  YIF1A  TEX2  DISP1  STIM2  ITGA11  ADAM8  CHPF  GPS2  HJURP  GLB1L2  SBSN  CDH6  TUBB6  TUBB3  LZTS3  PHLDA3  PHLDA2  MPDU1  IGFBP2  IGFBP1  CYB561A3  LSM1  ABHD15  METTL23  SCCPDH  PCYOX1  MAD1L1  RNF167  GLYR1  CSF1R  NOTCH1  CD151  IRS2  BRI3  AK1  AK5  AK6  MAEA  C1ORF53  C1ORF50  ABHD12  BAG3  APMAP  BTBD1  NDP  EFHD2  GYG1  SEMA7A  GPX1  GPX3  SLC37A2  GPX4  GPX7  C18ORF8  ASMTL  REXO4  MARCH3  PPP5C  CDK5  CDK4  VSTM2L  CDK1  SERINC2  TRMT61A  GTF2A2  MAPKAP1  PTPN22  FAM109B  FAM109A  DUSP14  METTL13  DUSP10  CNGA3  RNF25  SPAG7  ADRM1  MND1  RANGAP1  ENOX2  LAGE3  MGAT4B  BOLA3  TYRO3  RP9  UBA52  CLEC2D  SRXN1  SEMA3C  SEMA3G  RN7SL1  HIRIP3  GM2A  TRAP1  ACSL5  MRPS18A  ASS1  TMCC3  FAM64A  EIF5A2  SLC46A1  TP53RK  TTLL11  UQCR11  MYL10  NTSR1  AAMDC  B2M  FIS1  PINLYP  NUAK1  TUBG1  ARC  NCEH1  GRWD1  TTLL12  TEX264  ARL6IP1  HGH1  GDE1  LMAN2L  MGST1  PTDSS2  GNG11  ASUN  ASL  SKP1  TSC22D1  IGFN1  RHOC  RAD51C  PRPF19  RPUSD1  CEACAM6  TFF3  TFF2  CAMK1  TFF1  ALDOA  UMPS  TOMM5  TOMM6  TOMM7  FASTKD1  MAOB  STC1  CD99L2  STIP1  HINT2  NKIRAS1  PITRM1  DPH3  PRKCDBP  FADD  TNFAIP8L3  CC2D2A  C10ORF90  NQO1  RANBP1  TRAIP  TOMM34  NOV  TTPAL  MAPRE1  HRSP12  H2AX  FEN1  ATP5EP2  TOMM40  GDI2  RDH8  NPL  OLA1  HCCS  GUK1  SERP2  CXCR4  GDF11  EPDR1  B3GAT3  APLP2  FAH  BATF  DPM2  TP53I3  ATG16L1  SPNS2  PMP22  SCG5  STEAP3  ULBP2  ABCG2  FAIM  TMEM59L  SLFN5  GALNT18  RGS16  IFI6  CIB1  EFTUD1  TOMM22  MYH15  FOPNL  ABCF3  ABCF2  ATP6V0C  PSAPL1  ATP6V0B  EPHX1  DESI1  TBCD  TBCB  HIST2H2BE  NMRAL1  CKAP4  ALDH3A2  TMEM115  ALDH3A1  TRNP1  SRPRB  CHMP7  ATP6V1D  SH3KBP1  LOC101927746  LTBP1  SEL1L3  TMEM104  MAN2A1  ATP6V1F  JAGN1  HSPA6  HSPA8  TGFBR1  NFKBIB  ITGB1BP1  MAN2B2  PRTFDC1  CXCL8  CXCL2  CXCL1  CXCL3  DPP3  MOSPD1  CXCL5  DPP7  HEBP1  C9ORF16  NRIP3  LMNTD1  PDE4DIP  DCXR  KRT79  KRT78  KCTD5  KRT75  FGR  B4GALT7  B4GALT5  ZPR1  AURKAIP1  ACADVL  POLDIP2  TM2D2  USP11  USP12  CAMK2B  CHCHD5  PRRG4  NT5DC2  IGFBPL1  ALDH2  CDIPT  NIFK  SCYL1  GUSB | AKIRIN2  GK  WIPI1  VEGFA  RHOC  IL1A  DNAJC1  SLC7A8  UBE2S  RBP4  COL5A3  SLC25A33  S100A9  EIF5A2  SKA1  ZNF593  TNFAIP6  HJURP  CXCL2  CXCL1  AREG  CXCL3  CXCL5  SKA3  TUBB6  BCL2L12  ICT1  CSRP1  KDR  CYP1B1  TIMP1  S100A16  WLS  PHLDA3  MICA  OSBPL6  GCH1  INHBA  GLRX2  PRG4  ATP2B1  EIF1  FOSL1  FHDC1  DHRS9  STK17B  ALDH1A3  DNAJA4  AADAC  BIRC5  ANGPTL4  SMAGP  KIF2C  PTX3  GCLM  LINC00941  PID1  LRRC59  H2AX  NXT1  SLC43A3  IRS2  PLEK2  SH3KBP1  RORB  ABCA13  PTGS2  PTS  CCNB1  DNER  MICAL2  SERTAD1  PI3  NDP  NIFK  UPP1  METTL9  MFSD2A  IL13RA2 |
