## Supplemental Table 2 for "MondoA mediates transcriptional coordination between the MYC network and the integrated stress response in pancreatic ductal adenocarcinoma"

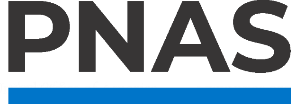

**Supporting Information for**

MondoA mediates transcriptional coordination between the MYC network and the integrated stress response in pancreatic ductal adenocarcinoma.

Paste the full author list here Erin L. Ramsey^1^, Stephanie Dobersch^2^, Brian Freie^1^, Nan Hyung Hong^1^, Xiaoying Wu^1^, Sita Kugel^2^, Robert N. Eisenman^1,^*, Patrick A. Carroll^1,^*

^1^Basic Sciences Division, Fred Hutchinson Cancer Center, Seattle, USA

^2^Human Biology Division, Fred Hutchinson Cancer Center, Seattle, USA

*Corresponding authors Robert N. Eisenman and Patrick A. Carroll

**This PDF file includes:**

Supplemental Table 2

**Supplemental Table 2: Down-regulated DEGs from RNA-Seq**

| **siMondoA Down** | **siMLX Down** | **SBI-477 Down** | **Core Down DEGs** |
| --- | --- | --- | --- |
| ENDOG  B3GALNT1  NIPSNAP3A  LUM  HEXIM1  STARD7  C5ORF15  MCTS1  ZFYVE21  KIAA1586  TMEM80  PRKAB1  SCAMP1  EML4  PTRHD1  EID1  COL1A2  RBP3  LAS1L  TFR2  RBP1  RFX5  UQCRC2  ORAI2  CYB5RL  LETMD1  SKAP2  CHIC1  MTMR1  JRK  MTMR4  HPGD  IFT172  C4BPA  CLDN3  SLC5A3  TFPI  ACCS  TPRG1L  SNX30  LSM11  SPDEF  DPYSL2  SAMD15  TMEM68  CAPNS1  TMEM69  SNCG  LINC00998  PPARGC1A  GBP2  MUC6  ENPP7P13  GBP4  ABCA2  ABCA3  GSTK1  JAG2  EDEM3  LRBA  NR2F1  LOC642852  NR0B2  FAM136A  AIF1L  HDHD2  PBX1  DHFR  FERMT1  MOV10  FAM111B  VANGL1  HSPG2  TBL3  QARS  FASN  SFXN1  PRKAR1A  LYPD6B  PSME1  SFXN3  DZIP1  C5ORF30  VAMP5  DZIP3  VAMP3  NOX3  LXN  TDRKH  GABARAP  C5ORF24  TTYH3  COL13A1  TMEM97  MYLK  AAAS  SULT1A1  GRIN2A  CYB5R4  NETO2  SCP2  SMCO4  ARPIN  LTA4H  RHBDL3  TMEM8B  ZNF488  GSTM2  FGFBP3  LYPLA1  GSTM1  JUP  KIAA1143  TMEM42  EMP3  IFT122  CP  DUSP9  SUMF1  LMBR1  DUSP7  LRRC20  DCBLD2  VWA5A  GPRC5A  PIGC  MBLAC2  GPRC5C  LYRM7  C4ORF3  PXMP2  BHLHE41  PXMP4  AGO1  TMEM106C  PIGM  MDFI  TMEM106B  ISPD  ERGIC1  SLBP  ARF5  ELOF1  SLC18B1  C11ORF1  ZC3HAV1  ACAT1  IQGAP2  MRPL36  SLC7A2  AACS  EXTL3  SHH  PPP2R4  EXTL2  TMEM25  PCMTD1  CGNL1  EGLN3  MAMDC2  STAT1  IFT140  MANEAL  TMEM64  RNASE4  CNPY4  LRG1  HEXDC  CASD1  NGFRAP1  ABCB10  ETNK1  MYRF  EBAG9  LGR4  TRUB2  OAT  ERO1B  QPRT  MPV17L  WRB  GSTP1  CIPC  TMEM53  TUBA1A  UNG  PTGS1  NUDT12  SCAMP5  GLRA3  HLA-DMB  ZFYVE28  KIAA1147  HLA-DPA1  AGR2  ZNF445  ZBTB18  SLC16A2  CORO2A  FBXW9  RASL10B  CTNNBIP1  ATP11C  MSH2  FOXP1  SLC9A7  MSH3  TTC30A  TTC30B  MAPKAPK3  PDCD4  S100A4  ROR1  TARDBP  MAD2L1  TRABD2B  PTPN4  SLC25A36  KANK2  SLC27A1  PDCD7  RASL11A  PDCD6  SRSF2  FAM117A  FOXO6  CASC4  CASC1  FOXO4  DEFB1  FAM117B  TSEN2  APEH  C1ORF233  LOC100129534  GGA2  LAPTM4B  LOC728554  BCL2L11  SH2B1  HAS3  PMPCB  ADORA1  PMPCA  PSKH1  ZNF664  KCNH2  NEK9  BCAS4  CPA1  TNFRSF10D  METRN  HPS6  EEF2  MED28  FOXN3  FBXO33  LRRC45  MED20  SMO  FSCN1  SMS  BIRC7  TMEM19  OSBPL1A  PFKFB3  FTO  COL17A1  PFKFB4  KCNE3  C11ORF95  LOC102724312  TACSTD2  MSI2  OLFML2A  HNMT  FBXO21  SNN  KIAA0895  DHRS4-AS1  UROD  THEM4  CNTNAP3B  G3BP2  EIF4EBP2  THEM6  BMF  ATP6V0E2-AS1  TRIM66  SYBU  PSMD5-AS1  CYB5B  HSP90AA1  BDNF  SPTB  TNFRSF1A  IMPACT  ARL4C  PIK3CB  NPC2  PCCA  IMPAD1  HLA-DRA  LOC257396  IREB2  CNTNAP3  SHC3  SHC4  FOCAD  POMT1  NREP  ADAMTS10  FBXO41  ADAMTS15  MDK  RABGGTA  ADAMTS14  ALAD  DRAM2  TKFC  KDSR  DSEL  ACP6  D2HGDH  BOK  B4GALNT4  B4GALNT3  KCNH3  MARCKSL1  NDUFA7  NDUFA5  RNASET2  TP53INP1  NDUFA2  MBOAT2  VPS33A  SORL1  TBX3  MYO1F  TSPAN15  SYT13  TXNIP  TRIM37  KCTD12  EIF3D  EIF3E  MMADHC  LINC00346  CTXN1  SPG21  EPHB6  ITGB4  FHL1  ITGB3  KIAA1551  SRI  ARRB2  PBXIP1  CXXC5  TNFSF13B  ZBTB42  PPM1H  ZNF608  ARRB1  SUMO3  UBQLN2  SLITRK6  PCDH7  CCL2  EPHB3  SLC25A23  EIF2D  CTHRC1  EIF2A  HNRNPA1  EPHA7  LINC00094  PLEKHA2  SFMBT2  FBXO2  PLEKHA1  ASRGL1  MYO7A  PPA1  PEX12  DHX40  MAPK14  GSTZ1  SFRP5  ST5  MT1A  CCDC80  MGP  LAP3  PPA2  SPATA13  ITGBL1  PEX11G  LRRN1  COL9A3  MXRA5  RIN2  C22ORF39  CALM3  TP53  AHCYL1  TMEM87B  ITIH4  CERK  KDELR2  CCDC69  PLEKHB1  PEX11B  LRP5  LRP3  CYB5D1  CYB5D2  STS  CDC23  ZNF709  C1GALT1C1  EXOSC5  CHORDC1  BSG  ADGRA3  PEG10  CPNE1  CPNE3  SLC12A8  RIBC1  SLC12A9  AAR2  TNFSF18  TNFSF15  ST6GAL1  ALG8  IQSEC2  FAM78B  ALG6  XRCC4  RASSF9  ANK3  ALG1  PTK7  IGF2  ANK1  PRG1  IFFO2  AIMP2  EWSAT1  RNF144A  PPP1R1A  WNK2  MYO5C  NET1  FBXL2  INTS5  APOBEC3C  LPGAT1  ZBED8  MTCL1  NRARP  ZBED5-AS1  TXNDC15  TXNDC16  RCBTB1  HYKK  LGALS2  C14ORF1  NUDCD2  C7ORF55  ARSK  TSPAN8  ANKH  PIEZO2  TSPAN6  RCC2  ACAD10  ZBED1  APOE  TNFSF10  DIRC2  TSPAN1  RIMKLA  TMEM150A  BBS2  BBS1  ZHX3  NXPE3  AKR1A1  FLJ10038  KLHL4  MRPS6  BST2  PTMA  ALDH6A1  TOX2  ID2  KLHL9  CCNG1  UFC1  CRELD1  CALR  ATPAF1  PTPRU  RABGAP1  MRPS16  HIBADH  CETN2  CCDC103  LINC00284  OMA1  RTF1  PCSK9  PTPRK  ACTR1A  ACACB  SEC14L1  SEC14L2  BABAM1  SH3BGRL  ARV1  NEU1  IDS  ORMDL3  CTSH  SOCS7  CTSF  PCK1  CAMP  CCT3  NPM1  MMD  SORT1  NUBPL  STARD3NL  TTC3  CACYBP  AKR1C3  KAZALD1  C9ORF116  APRT  DPY19L1  ETAA1  NOL4L  NUDT9  SEC23IP  ZADH2  NLRP11  KIF26A  CCNE2  TCEA2  CD47  CD46  FKBP11  FOXA1  DHRS11  AHCY  SYNM  DIRAS3  DERL3  PRICKLE4  PRICKLE1  CRIP2  COX5B  ZDHHC4  SPAG16  SERTAD4  SPTLC2  ZFP14  POFUT1  PNMA2  HYAL2  ENC1  HYAL3  NCS1  SUFU  DMBT1  FAM3C  MGAT5  THSD7A  MGAT3  PDGFRB  UBE2H  CD74  UBE2I  RARRES3  RARRES2  MRE11A  RPGRIP1L  KLHL24  AKR7A2  SDHC  SDHA  TMEM254  KLHL21  ALDH4A1  GPRIN3  SLFN12  GPD2  YPEL2  SIDT2  PLAA  YPEL3  IVNS1ABP  NDRG4  UBE3D  MPI  SCNN1A  CPOX  PEBP1  UBE3B  PRKX  ASB13  ASAP3  CELSR2  SLC1A7  NDRG2  CARF  ATOH8  TMEM245  TMEM246  RAB40B  GNPTAB  CAPN7  FAM172A  CAPN8  KIF5C  CAPN5  PERP  TBC1D14  PAK3  TBC1D17  SREBF1  TTC37  TCAF1  ARHGEF16  DSTN  FAM199X  TMEM230  TMEM231  GNB1  GEMIN5  GNB5  MXD4  VCL  FBN2  VIL1  PYROXD2  PODXL2  HTRA3  NAT14  PLAT  HTRA1  RASGRP3  FAM198B  HCAR1  CALHM2  EEF2K  CALHM3  SCN9A  PALM  KIF3C  COX15  MUC16  BCKDHB  BCKDHA  PAICS  NCBP2  CACFD1  TTC12  MSN  HNRNPA1P10  PEX1  PEX2  TTC19  GULP1  SSTR5  TAPBP  BMP4  RCN2  C7ORF60  MMRN2  PEX5  CPS1  EEF1G  PEX6  SPOPL  FBP1  PAOX  DNAH5  RTN1  DNAH2  YWHAE  UBA7  PSIP1  CHD6  HMGB1  PIK3R2  PLD2  CHD3  PHTF2  PHTF1  SIRPB1  AGPAT5  SYNGR2  MARVELD1  SEP15  CPT2  SARAF  WNT3  WNT4  LOC100506127  DACT2  CCDC176  MUC5B  GLRX5  DYNLT3  RRM1  DAPK1  PDPK1  TBC1D9  BISPR  YWHAZ  SUOX  SOD1  ADGRB1  ABHD17A  CHDH  ERCC8  GALM  TRIB2  NUP37  SNTB2  PRR7  CASC10  LDB1  TEX101  MVP  MSMO1  CLCN3  RND1  SAMHD1  ADD3  CYGB  CLN3  ACOXL  MAT2B  ABHD16A  MAT2A  SPATA24  IFT52  MAN1C1  DLEU1  EARS2  DNAAF3  AHNAK2  GLCE  RPL23  SYTL2  KIAA1462  SOX12  IFT81  IFT80  SYTL5  DKK3  WDR77  DYNC1LI2  MARCKS  ASB9  WDR81  GPR161  GPD1L  RPL26  HSPA1A  POLR3G  GLA  TARS2  RPL29  HSPA1B  CYFIP1  ZNF395  RPL11  SERPINE2  RPL12  PPOX  FNBP1L  GNG2  NWD1  GNG4  CREB3L4  MYB  NTHL1  CHST12  HENMT1  MYC  WDR70  ABI3BP  PLCE1  ST3GAL5  HOXA3  PIP4K2B  SERPINF1  S100A13  RPL14  ZNF561-AS1  CEP97  CHST15  VPS16  CSRP2BP  NCSTN  FDFT1  HOXA5  FZD2  EDARADD  TSPYL1  ICMT  FZD4  TSPYL2  LINC00673  AFAP1L2  RPL41  FZD5  FZD7  PLK2  EPT1  TYSND1  WDR54  RMDN3  FN3KRP  KIAA0391  EHD2  SQLE  SLC6A6  BACE2  EHD3  BCAM  WDR60  SSPN  HBP1  CEP68  RPL5  HSPA4L  CFD  RPL31  RPL4  DANCR  CFI  TRIL  RPL34  PKI55  STMN3  VPS26B  ATXN7L3B  ARPP19  IPO11  WDR47  RRP9  FAM122B  GCSH  PRDX3  ALCAM  GJA3  STMN1  LINC01503  AP1S3  MCMBP  TMEM200A  RPL39  NUDC  ZNF362  ANXA4  NDC1  TET1  NUTF2  TYW1B  WDR35  H19  WDR34  TAF6L  FAM210B  PAX9  VMA21  EXOC4  RNF141  PCYOX1L  ZNF358  GAS7  GAS6  GPI  SERPINA3  BHLHB9  PHF20  BTG2  ENDOD1  TRIM2  SERPINA5  SERPINA6  FAM171A1  RNF157  TMEM168  LDHA  NDNL2  RHPN2  TIMP3  PBK  TIMP2  TK2  TEAD2  TMOD2  SF3A3  SLC2A12  MGMT  WDR19  SLC2A13  ELP2  UBE4B  DHRS1  FAM213B  ELP6  FAM213A  TUBB4B  SYNJ2BP  LZTFL1  MAP1A  ZNF33B  AGBL5  ZNF33A  PLCB4  LPCAT4  COL4A5  IVD  LPCAT3  LAMTOR5  C16ORF89  MSRB1  SETD5  PFAS  RARG  NDUFA10  LPCAT1  TCF21  GRAMD4  SLC8A1  PRSS21  ADAMTSL2  ZSCAN30  UBL5  ING4  IMPA2  TCTN3  TCTN2  TCTN1  GCNT3  METTL7A  CKB  ANXA9  TMEM38A  CHML  DEPTOR  TMEM14A  GSN  BCHE  MREG  TLE2  SEMA4F  LINC00493  SEMA4G  THNSL1  C1ORF123  NRBP1  COPS4  ITGA10  RAB37  LIFR-AS1  LRFN1  BAIAP2-AS1  CSPG4  ANKRD36C  FKBP7  CHRM3  VKORC1L1  POGLUT1  CD177  GTF2I  FKBP2  CSF1  TUSC3  TCEAL1  GLT8D1  GLB1L2  FAM214A  FAM105A  PAPSS2  TCEAL8  RAB26  CDH2  TCEAL4  RBBP4  TPM1  RBBP9  RBBP7  RPS3  DTD2  ACSS1  SIAE  SEMA6B  F2R  IGFBP6  CHPF2  IGFBP5  ATP8B2  IGFBP4  IGFBP3  CYB561A3  GTF3A  RITA1  TRAPPC9  PHKB  PGD  REEP2  DUSP23  TFDP1  REEP6  TPP1  PCYOX1  RAB5B  EIF4G3  CSF1R  RAB3A  SLC26A2  CALCOCO1  CLSTN3  CNP  LINC00261  INSIG1  FAM216A  AK4  ZBED6CL  MAEA  MYOM3  TM7SF2  CDC14B  NPEPPS  TM7SF3  SALL2  FGD5-AS1  ABHD10  BAG1  C1QTNF6  GYG1  PLS3  KCNJ12  HMGCS1  LANCL1  SPACA6P  PTPN11  SORD  DDHD2  ZNF75D  PTPN14  TPCN1  NIPSNAP1  C15ORF57  CUL4B  CDK5  MFAP3  ZG16B  NES  SERINC1  SLC26A4  EIF4E2  OSCP1  BMPR2  FECH  COL14A1  GLO1  DUSP19  MTM1  LBH  FGGY  PLTP  DENND3  MGAT5B  VPS54  RPL36A  SERAC1  LINC01521  TPPP  RALGDS  SLC37A4  MAGI1  PLA2G12A  HN1L  IRX3  PLEKHG3  RALGAPA2  RPH3AL  ILF3-AS1  RPL35A  CTDSPL  TSN  SELENBP1  DNAJA1  CLEC2B  TST  POLR1B  CCDC64  ANG  EIF4A2  HDAC6  NCKAP5  LSP1  HDAC1  KIAA0100  SEMA3F  SEMA3G  DCAF17  HECTD3  GM2A  RNPEP  SCRN3  WDR6  UBTF  BCO2  RPL7A  MLXIP  CDON  TUB  SS18L1  KRT4  KRT8  TNRC6C-AS1  PPP2R1A  P3H4  SEPP1  MTURN  CYP2U1  SUCLG2  TLN2  OBSL1  APP  SLC35B4  IFITM2  IFITM3  FGL1  IFITM1  VIPR1  SLC46A1  SLC35B2  AQP5  HS6ST1  CARHSP1  AQP1  HS6ST3  AQP2  AMER1  UHMK1  C3ORF18  ANTXR1  NIPAL3  MEST  RGAG4  PTGDS  INSR  TMEM50B  ARRDC3  ARRDC4  SLC6A12  HLA-A  ACADSB  PTOV1  CMBL  HLA-H  GBAS  VASN  IL17RC  VASP  INPP4B  ALDH7A1  PARP10  ADCK3  KLHDC8B  CYP2S1  TSHZ1  CUTA  SPARC  ARL6IP1  FAM89B  PDE1B  TSHZ2  TXK  MLC1  TXN  TMTC4  RAPGEFL1  IFIT2  IFIT1  C14ORF93  NLK  GMCL1  NLN  PRRC1  MARC1  EPM2AIP1  SYNPR  RPS18  RPS17  MAATS1  PTDSS1  C10ORF32  PDK4  ARL6IP5  TREX1  TP53BP1  RPS12  ERMAP  TRPM7  PDK3  TNS3  SKP1  SKP2  TNS2  AGFG2  ZER1  BTN3A2  BTN3A1  PTGES3  GNB2L1  BTN3A3  NME1  CLDN12  CLDN11  CLDN10  KBTBD6  CEACAM1  TPPP3  KRT19  MAPT  KRT18  VWA1  FAM20C  TFF3  CLPTM1L  MRRF  IGSF9  SLC44A2  MAGEF1  SLC35F5  CD99L2  LIMD2  CX3CL1  HINT2  ZMYM3  MAGT1  CAMKV  MLEC  STRA6  CLIC3  ASIC1  MAP9  ASIC2  PRMT7  TM4SF18  ZCCHC24  PRMT6  PALD1  LMLN  SUZ12  CBR1  CGREF1  ATP6V0E2  FAM46C  PARP3  PRMT2  MCCC1  SUSD3  SCD5  MCCC2  SACM1L  PARP1  SLC35G1  VASH1  VASH2  THOC3  ZDHHC17  PODN  RAB11A  PARP9  ALDH5A1  PRPF39  SPCS2  LOC79160  HNRNPH2  RAB9B  TRIP13  C21ORF2  CDK5R1  ANKRA2  DCP2  TJP3  HRSP12  FEN1  RYR3  MEGF6  ZDHHC20  CTNNB1  HCCS  RSF1  FCGBP  CST4  AMN1  CST1  CXCR4  SOSTDC1  LMO2  KLHDC3  CA12  QPCTL  CA11  GINS1  TGFB2  EPDR1  LIFR  TARBP1  PRSS56  GRHL3  ATP1B1  ARHGAP24  MYCL  MAU2  SHISA2  DNAJC30  BAMBI  GSTA4  FAM185A  ELMO1  TGFBI  PDE7B  B9D1  GALNT15  TNXB  SUCNR1  TMEM59L  HFE  FBLN2  TOMM20  EFNA5  NOL6  GALNT11  DOCK11  EFNB3  LOC220729  EFNB1  TMEM129  DNAJC22  C1RL  HCFC1R1  GPBP1L1  FOPNL  MYH10  GAS2L1  REPS2  TRAPPC2L  ACBD7  IDH1  WSCD1  MPST  EPHX2  DNAJC12  TBCE  TBCD  CTPS2  SMAD6  UPK3B  PDIA5  EFNA1  TBCK  DNMBP  DLG3  TMEM9  DNAJC10  TRNP1  EBPL  CDH15  SERTAD4-AS1  MATN3  KIAA1191  ISYNA1  DOCK4  ROGDI  ARHGAP18  LTBP3  C17ORF96  MYL6B  PLCXD3  TMEM100  ARHGAP23  TMEM101  ZMAT3  FAM162A  LARGE  ALDH3B1  MAN2A1  SVEP1  ABCC4  CMTM8  KDM4B  CBX6  JAGN1  HSPA8  CMTM6  CBX2  CBX1  IQCE  RPS3A  OAS1  TGFBR2  HPDL  GRHPR  COQ4  FAM189B  MEIS2  RBL2  MEIS1  RASA4  B4GAT1  H3F3AP4  PDE3B  RBMS1  FAM84B  FAN1  IDI1  C2ORF44  HAPLN1  HSPB8  OPRL1  HS2ST1  LRRTM4  MFSD3  MFSD1  PDE4B  TCFL5  PTGIS  CAMK1D  TCF7L2  TMEM30A  GLP2R  KCTD3  C14ORF132  POMGNT2  MBNL1-AS1  WEE1  MLPH  CLDN4  TRAPPC6A  ANGEL1  VDAC3  VDAC1  SPRY1  SSBP2  PFDN5  SSBP3  PXYLP1  B4GALT6  TSEN34  ABCD4  ZNF174  TM2D3  PCDHGB5  TM2D2  BRAT1  TM2D1  EHMT2  PPP2R2B  DDN  THBS3  HSPD1  ATP5G1  CCDC28B  RPS4X  RMND5A  RBM3  ALDH1B1  CHCHD7  ENOSF1  LRIG1  CAMK2G  GUSB | NIPSNAP3A  C5ORF15  APCDD1L  CACNA2D4  LNPEP  PRKAB1  C12ORF76  PSMB9  COL1A2  RBP1  RFX5  FAR2  CYB5RL  LETMD1  HCP5  JRK  PGAP2  HPGD  SLC5A3  SZRD1  CLDN2  PRKAA1  TFPI  TPRG1L  DPYSL2  SNX29  STAT6  PPARGC1A  GBP2  MUC6  LINC00996  GBP4  ABCA3  GSTK1  HLA-DRB5  ATP6AP2  ABCA7  ERLIN1  LOC642852  VPS13C  FAM136A  HDHD1  DHFR  C12ORF49  VAMP8  QARS  PKIB  C6ORF89  LYPD6B  PLBD2  VAMP5  VAMP3  NOX3  HLA-DRB1  TAPBPL  RBPJ  C3  LIPK  PRKAR2A  SMCO4  LTA4H  CYB5R1  JUP  FN1  CP  SUMF1  DCBLD2  CS  TOMM40L  SUMF2  GPRC5A  GPRC5C  GPRC5B  LYRM5  PXMP2  BHLHE41  PIGM  TMEM106B  AVL9  ARF3  FAXDC2  ARF1  IQGAP1  IQGAP2  SLC7A2  AACS  TMEM25  CGNL1  MAMDC2  STAT2  MBD2  RNASE4  SLC39A13  CNPY4  RNASE1  TMEM59  NUDT21  ADAM15  ZNF219  HLA-DPB1  C11ORF54  HEXA  QPRT  ATL3  PTGS1  IKBKB  PSMB8  HLA-DMB  SMPD1  HLA-DPA1  AGR2  ZNF689  HLA-DMA  MYLIP  SEC24C  GOLM1  TUBA4A  SLC9A7  PDCD4  S100A4  CDKN2AIPNL  SQSTM1  ROR1  AGRN  TRABD2B  C11ORF71  UBE2D1  FOXO4  DEFB1  HSP90B1  BCL2L13  LAPTM4B  SLC25A44  KCNH2  ATP9A  NEK9  TMEM19  BIRC3  FTO  COL17A1  ZNF512B  FAM13B  TACSTD2  HNMT  HHAT  SNN  FNTA  EIF4EBP2  BMF  SYBU  ZKSCAN1  DGCR2  PSMD5-AS1  ITGA2  TNFRSF1B  ARL4C  UST  NPC2  TMBIM4  ZNF638  HLA-DRA  TRIM56  NFE2L3  MCFD2  FOCAD  ADAMTS10  ADAMTS15  MARCKSL1  MBOAT7  ERAP1  ERAP2  TP53INP1  SORL1  EIF3M  TXNIP  C1R  ITGB5  C1S  ITGB4  KIAA1551  NENF  ETFB  PBXIP1  TNFSF13B  PPM1H  ZNF608  ARRB1  SLITRK6  PCDH7  ITGB8  LRRC8B  HMG20A  ITGAV  NOMO1  PLEKHA1  MAPK14  GSTZ1  SFRP5  ST5  MGP  PPP1R3D  TRIM14  ITGBL1  PEX11G  LRRN1  MXRA5  EPHA4  TP53  MT1E  ITIH4  CCDC69  PLEKHB1  PEX11B  LRP5  PSMB10  RAB11FIP1  C1GALT1C1  BSG  ADGRA3  TGOLN2  SLC12A8  WWTR1  TNFSF18  TNFSF15  XRCC6  IQSEC2  TGDS  RASSF8  RAB27A  PTK7  IFFO2  MYO5C  CYSTM1  PPP1R11  PTTG1IP  RCBTB1  WFDC3  C14ORF2  C7ORF55  TSPAN8  ZBED1  APOE  TNFSF10  ARSD  CTSB  BBS1  NCOA4  KLHL3  KLHL4  MRPS6  PTPRE  BST2  TOX3  ALDH6A1  ID2  CCNG1  CRELD1  TMED10  HIBADH  MRPS12  CCDC103  OMA1  CTSZ  TGFA  PTPRK  TNFRSF11B  CTSO  SH3BGRL  ARV1  IDS  ORMDL3  MLX  CTSF  CTSD  SPRYD4  SORT1  GAA  AKR1C1  AKR1C3  KAZALD1  TCEA2  CD46  AKR1B1  SLC43A2  PRICKLE1  PTMS  CRIP2  SH3RF2  SERTAD4  POFUT1  OS9  ENC1  HYAL4  DMBT1  SDF4  THSD7A  RAC1  LOC100506548  CD59  GIT1  PDGFRB  UBE2H  CD74  MORC4  TBC1D2B  RARRES3  RARRES2  KLHL24  KLHL21  EEF1A2  ALDH4A1  UBE2Z  TMX4  YPEL2  SIDT2  NDRG4  SCNN1A  CELSR2  SLC1A7  CAAP1  NDRG2  ATOH8  TMEM245  FAM172A  PERP  GPC6  ARHGEF40  TCAF1  DDX58  DSTN  MR1  TMEM230  CAMKMT  MFGE8  MXD4  BLVRB  HOMEZ  LGALS3BP  JADE2  NAT14  PLAU  PLAT  HTRA1  RASGRP3  HCAR1  RGS4  PPIP5K1  CALHM3  RGS1  SCN9A  FAM174A  MUC16  BCKDHA  GALNT7  NCBP2  MMP1  MMP2  CAV2  MSN  GALNT2  CAV1  ETV1  ABHD14B  LAMB2  PEX2  HIGD2A  GULP1  TAPBP  BMP4  CPS1  RCN1  PEX6  MANSC1  CHI3L1  DNAH2  UBA7  CHD6  PIK3R3  PLD1  PHTF2  PLD3  PHTF1  SYNGR2  IRAK2  ERLEC1  SARAF  FBXL19-AS1  PDLIM5  DACT2  MBTPS1  FAM98A  GLRX5  DAPK1  BISPR  ATP2B2  FNDC3B  YWHAZ  SOD1  GALC  PNKD  CAT  MAN1B1  GALM  CHURC1  SNTB2  RERG  LAMA4  MVP  EMC10  CLCN3  ABHD16A  FAM73A  NPTN  CDK19  AHNAK2  GLCE  AFAP1  SYTL2  KIAA1462  SYTL5  KIAA1217  CPEB2  OSTM1  CD9  GPD1L  PLCD1  CPVL  TES  PRSS2  PRSS1  MAML3  AHNAK  SERPINE2  DCAF8  DCAF7  GNG2  FAM120A  NWD1  GNG4  CREB3L4  FLRT3  ABI3BP  FAM120B  PLCE1  SERPINF1  S100A10  APOL2  NCSTN  APOL1  APOL3  FZD2  NDFIP1  TSPYL1  TSPYL2  AFAP1L2  FZD6  FZD5  PLK2  MCAT  BACE1  BCAM  SSPN  HBP1  CFB  CFI  TRIL  STMN3  GNS  ATXN7L3B  KIAA0141  GLDN  SCPEP1  AP1S3  FLJ23867  LINC00657  KIAA0319L  ANXA4  CARD6  FAM102A  FAM210B  RNF141  GAS7  BHLHB9  BTG1  CHM  TRIM2  SERPINA5  SERPINA6  FAM171A1  TMEM168  ERMP1  C16ORF58  SSR1  TIMP3  TIMP2  DMTN  SLC2A12  ELP2  ZNF33B  CLIP4  AGBL5  PLCB4  LPCAT3  GRN  LPCAT2  KIAA1033  TCF21  PRSS21  TMSB4X  FCGRT  ING4  TCTN2  GPNMB  TCTN1  METTL7A  ANKRD52  TMEM38A  RNF44  BCHE  SEMA4F  TAP1  HIPK2  RAB30  DISP1  ITGA10  SYPL1  TRAFD1  KAT6B  ITGA11  ADAM8  TCTA  C8ORF33  FKBP7  FAM8A1  CD177  NATD1  CSF1  TCEAL1  CLU  LCLAT1  GLB1L2  FAM105A  GOLGA7B  TCEAL8  ABLIM3  RAB26  CDH2  CDH1  TPM1  PAXIP1-AS2  AP3M1  SIAE  TRAPPC1  SEMA6B  F2R  CHPF2  IGFBP3  CYB561A3  PHKB  PDCD6IP  TPP1  EIF4G2  SLC26A2  CALCOCO1  CLSTN3  CBFB  INSIG2  ZBED6CL  HID1  MYOM3  CDC14B  NPEPPS  TM7SF3  ABHD10  C1QTNF6  SLC15A4  C15ORF52  PTPN14  TPCN1  DNAJC3  SMDT1  CPD  ZG16B  SERINC1  SFT2D3  SLC26A4  SERINC3  COL14A1  LBH  PSAP  ZNF507  ZNF503  TPPP  RALGDS  MAGI1  MTOR  CTDSPL  SELENBP1  DNAJA1  PPAP2A  APOC1  TST  MGAT4B  ASPH  ANG  LEPROT  RNF13  WDR1  LSP1  SEMA3C  SEMA3F  THSD4  AP1G1  HECTD3  RNPEP  UGT1A6  BCO2  UGGT1  CACNG6  CDON  SLC35A2  ANKEF1  TNRC6C-AS1  P3H4  KATNAL1  OBSL1  APP  SLC35B4  SLC46A1  IFITM1  TRANK1  HS6ST1  HS6ST3  UHMK1  NIPAL3  ARFIP1  B2M  TMEM50B  ARRDC4  HLA-C  HLA-A  HLA-B  PUM2  IL17RE  HLA-H  VASN  ARFGAP2  HLA-E  IL17RC  PARP14  INPP4B  PARP12  PARP10  RPS23  CDS2  TSHZ2  TMTC4  GDE1  TEX261  PPL  IFIT2  IFIT1  IFIT3  SYNPR  GOLGA1  PDK4  TREX1  FAM65C  TNS3  SKP1  AGFG2  TNS2  BTN3A2  BTN3A1  PTGES3  BTN3A3  ITFG1  CEACAM1  VWA1  TFF3  AKR1B10  MEGF8  DAZAP2  CD99L2  ZMYM3  TMED4  TMED2  TMED7  TMED5  ASIC2  TM4SF18  ZCCHC24  LMLN  SVIL  ATP6V0E2  FAM46C  SUSD1  PARP1  PACS1  RAB11A  SVIP  PARP9  SPCS3  LOC79160  NOV  NBR1  RAB9B  SLC22A23  MAPRE1  FYTTD1  ZDHHC20  NUCKS1  FCGBP  CST4  CST1  CUX1  KIF13A  DLAT  CA12  CA11  EPDR1  LIFR  GRHL3  ATP1B1  TUG1  ARHGAP26  ARHGAP24  CAPZA2  ELMO1  TGFBI  LRP10  PSMB8-AS1  SUCNR1  CWC25  MANEA  GALNT10  PURB  EFNB1  C1RL  S1PR3  ZC3H13  EPHX1  MESDC2  HIST2H2BE  EFNA1  DNMBP  TMEM9  TRNP1  SERTAD4-AS1  ATP6V1A  ARHGAP18  LTBP3  SEZ6L2  PLCXD3  TMEM100  TMEM101  ZMAT3  COL4A3BP  MAN2A2  LARGE  ALDH3B1  TMEM109  SVEP1  KLRC2  FAM81A  KDM4B  DARS  JAGN1  CMTM6  OAS1  COQ6  TGFBR3  MEIS2  RBL2  LPIN1  RASA4  PDE3B  FAM84B  MAN2B1  GPSM3  NCAM1  HSPB8  LRRTM4  MFSD6  IGBP1  PTGIS  CAMK1D  TMEM30A  GLP2R  C14ORF132  SHCBP1  CLDN4  TRAPPC6A  SSBP2  B4GALT5  DHTKD1  CAMK2D  ACADVL  THRB  THBS3  RMND5A  RPS6KA3  SAP30L  PCDHGC3  LRIG1  GUSB | STARD4  NIPSNAP3A  LUM  FNBP4  NIPSNAP3B  BTBD19  STARD9  RFX3  DICER1  C20ORF194  RFX2  GRIN2D  PRKAB2  EML3  EML4  L1CAM  COL1A2  COL1A1  SCD  RBP3  RFX7  RBP1  RFX5  RUFY3  CYB5RL  SKAP2  CGGBP1  PGAP1  JRK  MTMR4  SAMD9  IFT172  CRABP2  C4BPA  C4BPB  DLL1  CLDN2  TFPI  CLDN1  UACA  LSM11  MUC1  MUC2  TMEM67  AKT3  BBX  SNORA18  LINC01604  GBP1  GBP3  MUC6  GBP4  PROSER3  ABCA2  SNORA51  GABARAPL1  PRPF38B  ABCA6  ABCA7  LOC389602  KLF10  LOC642852  NR2F1  PHKA2  VPS13A  PBX1  MOV10  GIGYF1  HSPG2  MLLT10  FASN  FOXL2NB  RNF207  VAMP1  DZIP3  EZH1  NOX3  LOC729218  MDC1  SNORA40  LXN  TDRKH  ZNF251  CEP85L  DDX12P  PDGFC  PDGFB  BAZ2A  ZNF22  F11R  LIPC  ISLR  GATA2  TRIOBP  SUV420H2  C3  LIPG  KIAA1107  SULT1A1  KIAA1109  EPB41L1  TDO2  EPB41L2  LIPH  ZC3H12B  CFAP69  ZNF248  ZBTB7B  TMEM8B  ZNF488  RHBDL1  ANKIB1  CTC1  CROCCP3  EMP1  CP  TMC6  TMC5  GOLGA2P5  GPRC5A  GPRC5B  COL3A1  TMCO6  AGO4  INCENP  ADGRL1  ZNF236  TMEM106B  ZNF234  IL18BP  EXD3  FAXDC2  PIF1  IQGAP2  PEAR1  NKILA  BBC3  ATXN3  TCP11L2  RPL23AP64  DLGAP1  ZNF225  ZNF224  LOC146880  PCMTD2  ELOVL2  MEF2D  MBD6  MAMDC4  IFT140  LOC101928100  RNASE4  LRG1  SPSB3  ADAM17  ETNK2  FLJ22447  PKN3  ZNF217  MYRF  PKN2  SLC29A2  PTPRCAP  SLC29A4  CDKL1  STAG3L2  ZNF211  ERO1B  MTMR11  LRRC14  ZNF692  QPRT  JRKL  PTGS1  NUDT12  NDST2  IKBKB  GLRA3  KIAA1147  MPRIP  CAPN10-AS1  AGR2  RICTOR  NINL  C5ORF63  THAP9-AS1  FADS2  ZNF202  JAK2  ZSWIM4  MYLIP  ZBTB18  FAM227A  LRRC61  ZBTB12  NEAT1  SLC9A3  ATXN2L  CLUHP3  DMTF1  TUBGCP6  CNTROB  ROR1  SLC25A37  PVRL2  MIR210HG  TRABD2B  ZNF432  SLC25A36  SLC27A1  CCNT2  ZBTB26  CASC8  HR  ATN1  LOC100289230  PLOD2  FOXO4  DEFB1  NPAS2  HSP90B1  CTGF  ATXN1L  FRZB  SH3BP2  LCNL1  SH2B1  FLNB  SLC16A4  NKTR  SLC16A5  ATP9A  C21ORF58  LINC01000  LRRC49  KCND1  SLC38A2  ADGRV1  BNIP3L  LRRC40  C1ORF220  INADL  FBXO36  FOXN3  IBTK  PIAS3  GABPB1-AS1  MED23  MAG  KIAA1755  MLXIPL  ZNF419  CDC42EP4  ZNF418  LCN2  MDM4  BIRC7  SNRNP200  BCAT1  BIRC2  BIRC3  ZNF652  FTO  KMT2E  PFKFB2  PTOV1-AS2  EXOC3L4  ZRANB2-AS1  ZNF512B  KMT2B  BOD1L1  LOC102724312  UBFD1  LRRC56  ELFN2  GLIS2  MUM1  CTIF  NIPBL  HHAT  ZKSCAN8  KIAA0895  PIM1  FUK  BMF  EDIL3  TRIM66  PSMD5-AS1  LINC00330  CYB5B  PPP1R12B  APAF1  AHSA2  LOC644656  SPTB  TNFRSF1A  QKI  DBNDD1  LOC100130987  ERBB2IP  DSG2  OGT  ZNF37BP  NFE2L3  IREB2  CNTNAP3  GTF3C3  L3MBTL1  NREP  EFCAB13  FOXK1  LRRC16A  BLACAT1  SAV1  MLLT6  ADAMTS10  CRHR1-IT1  ADAMTS14  TMEM147-AS1  KIAA1522  RDH11  C10ORF10  B4GALNT4  B4GALNT3  SRCIN1  ACVR2B-AS1  GNRHR2  ZBTB39  PVRIG  ERAP2  NICN1  ARNT  TP53INP1  TSPAN15  SYT12  ZNF618  HNRNPU-AS1  TXNIP  ZNF614  PABPC1L  KCTD12  ZMYND8  LINC00342  C1R  TRIM31  C1S  ITGB4  ITGB3  KIAA0895L  KIAA1551  DNHD1  ZBTB46  LENG8  PBXIP1  SLC25A27  SYNGAP1  PPP1R13L  ZNF608  KCTD20  LOC100288123  ZNF204P  IGF2BP2  KRIT1  CCL2  EPHB3  ZNF841  EPHA7  PLEKHA2  ARL14  PLEKHA1  TM4SF4  MOGS  NBEAL2  MYO7A  PLEKHA4  MAPK15  PLEKHA8  TM4SF1  DHX40  MAPK12  ST5  CCDC80  HIP1R  PPP1R3E  MTCP1  FAM76B  ULK1  ZNF711  COL9A3  MXRA5  EPHA2  ALPK1  STX1B  SNORD45A  SNORD45B  CCDC68  PEX11A  LRP8  PARD6B  HERC3  ELMSAN1  IFI16  C7ORF31  SLAMF8  PPP1R32  CCNL2  CCNL1  ARVCF  FER1L4  YES1  ZNF436-AS1  PPP1R26  TNFSF18  FAM9B  FAM78B  XRCC4  RASSF9  IGF2  ANK1  IFFO2  EWSAT1  SNHG10  FBXL2  RGL4  TLCD2  GRB7  FBXL8  RCOR2  SUGP2  PAXIP1  MTCL1  LGALSL  LOC100129917  ICAM5  MTMR9LP  GCC2  IKZF4  ZBTB5  CCT6P3  RASSF3  FAM86B3P  MBTD1  SNHG20  TNFSF10  ACAD11  LGALS8  RIMKLA  SEC31B  GABBR1  NBPF9  NBPF8  TMEM198B  FLJ10038  PRPF40B  NBPF12  KLHL5  NBPF15  BMS1P4  NR5A2  ICK  PTPRF  SLC25A25-AS1  TOX3  TOX2  ID1  CCNG2  CCNG1  FAM193B  CD22  SLC35E2B  DOT1L  DDR1  GIPR  USPL1  LINC00284  PTPRN  PCSK9  EVPL  MIR4697HG  PTPRH  SBNO2  SEC14L1  STK36  CNTRL  BBS9  CTSH  CCR7  DFNB31  PCK1  SOCS6  H1FX-AS1  ANO8  MIB1  PER3  SGSM2  SH3RF3  ETAA1  NOL4L  TTC6  PER1  KCNQ4  FOXD2-AS1  MGARP  ENGASE  LOC100288637  PNPLA3  ERRFI1  FOXA1  SIRPG-AS1  TBX2-AS1  DERL3  SERTAD4  MNT  ZFP14  DMBT1  THSD7A  LOC100506548  DKFZP434I0714  MGAT3  ANKRD13B  RARRES1  PDGFRB  SPINK4  SUGT1P1  DENND4B  RARRES3  KLHL24  CRIM1  VCAM1  TANC1  IFI44L  TANC2  DOK3  SLFN12  MZF1  COL5A2  YPEL2  SIDT2  AMOTL2  CD69  OTUD4  YAP1  PRUNE  PVT1  LOXL4  SCNN1A  SLC1A3  ASAP2  PRKX  KLHL36  CELSR2  ATAD2B  SAMD9L  NUP160  KIF5A  ATOH8  RELN  BAHCC1  TAF1D  FAM196B  HSF4  SGK494  PPFIA4  SREBF1  TTC37  SREBF2  GGT5  ARHGEF11  TCAF2  TTC33  ARHGEF16  ARHGEF17  MINK1  SSH3  NAT6  PAN3  PAN2  TAF4B  RSRP1  CARD14  MIR3064  ARHGEF25  ICE2  KLHL17  SLC41A3  CCDC125  HTRA3  UVSSA  PLAT  ETS1  CACNA1I  DDX60  RGS5  FAM198B  TGM2  CNN3  CNN2  RASGRP1  GYLTL1B  RGS4  CALHM3  PALM  RGS2  ADGRG6  SBF2  BCL9L  ARID2  ZNF385A  TTC14  SHROOM3  SSTR1  TNIP1  ETV4  GPCPD1  SSTR5  ACVR2B  CLK2  BMP4  IRF1  CLCN6  C7ORF60  ELF3  BMP1  RBM20  IRF9  RAD51-AS1  LTB4R  NNMT  DNAH2  ZNF192P1  CHD6  HMGB2  TNFAIP2  PLD2  CHD3  PLD1  CHD2  AGPAT4  PDLIM1  CYP26B1  SYNGR3  MKNK2  ECT2  CA9  PDLIM5  WNT4  LOC100506127  DACT2  GALNT9  CDK11B  CDK11A  DDX17  LOC101928505  ATP2B2  LMBRD1  PSRC1  TBC1D4  C19ORF66  ADGRB1  ADGRB2  CRY1  DAAM1  MIR5047  RBM44  MVK  CASC11  RERE  LDB1  LAMA5  TEX101  ZNF292  RB1CC1  HMGCR  MSMO1  LENG8-AS1  CRYZ  IL1RL1  CYTH3  HSP90B2P  MAT2A  EPS8L2  BNIP3  RBM33  CDK18  DNAAF3  MGC57346  ZNF160  AFAP1  SOX12  IFT81  IFT80  GTPBP2  GANC  MEX3A  LOC730101  MARCKS  LGALS12  LINC00887  PLCD1  NHLRC3  ALG10B  PRR7-AS1  PCED1B-AS1  TES  PRSS2  SNORD80  EHF  PRSS1  ZNF395  MAML3  COL12A1  LOC284454  BACH1  FNBP1L  ADCY7  SSFA2  GNG2  PRTG  MILR1  CREB3L1  FBXL20  KIAA0355  MXI1  ABI3BP  PLCE1  ST3GAL5  HOXA3  SERPINF1  HOXA2  HOXA1  CHST15  FBRS  FDFT1  HOXA5  FZD2  TSPYL4  ST18  TSPYL2  DNAH11  FZD5  TIGD7  PLK2  PRRC2B  FOSL2  EHD2  SLC6A6  SQLE  BCAM  SLC6A9  WDR60  TCN1  ZNF702P  HOXB4  HOXB3  PRKD2  WDR5B  HBP1  HOXB9  HOXB6  NAA40  TRIO  CFI  STMN3  NEDD4L  RAPH1  ATXN7L3B  MYSM1  SMC5  GRPR  PLAC8  ALCAM  MYO18A  HOXA11-AS  PLCG1  GARNL3  HOXB-AS1  CCL26  GSDMB  RPS6KL1  SHPRH  ANXA3  ANXA4  HOXB-AS3  NFATC2  NFATC1  WDR35  MIR614  SCARNA27  PGM5P2  NCOR2  SIKE1  PAM  SERPINA3  WDR26  BTG2  WDR27  TINAGL1  YTHDC2  ABHD4  PRDM15  TRIM2  METTL21B  VSIG10L  SERPINA5  FAM160B2  GJC1  LOC100130238  TRIM5  RHPN1  PLLP  LOC100131564  CIC  TEAD2  FAM160B1  TIGD1  MSTO2P  BRD3  MBNL2  SLC2A12  SERPINB1  ZNF580  WDR19  NR1H4  CIT  SERPINB9  DHRS3  KIAA0195  TMEM154  SYNJ2BP  LZTFL1  ZNF33B  TMEM156  CLIP4  AGBL5  NFIB  NFIC  COL4A5  TOM1L1  UBE2V1  ZNF337  ZNF577  ANGPTL2  ZNF334  ZNF333  SETD5  CDKN1B  PFAS  PHLPP1  CDKN1C  ZNF570  PHF2  SPIN3  STON1  GLI2  SLC8A1  GRAMD3  LMAN1  IFIH1  LFNG  ZSCAN30  ING4  ADAMTSL4  LOC100129034  TCTN2  MIR22HG  METTL7A  PCGF2  ANKRD52  ZNF320  RNF44  SEMA4B  ATHL1  ANKRD27  PLXNC1  ANKRD23  MITF  SEMA4G  MOB3A  RAB30  PNRC1  NRBP2  C6ORF141  OBSCN  CNOT8  KAT6B  SLCO2A1  LIFR-AS1  ABI2  LTB4R2  GAPDHS  ANKRD22  BAIAP2-AS1  PLXNB3  ZNF558  PLXNB2  PLXNB1  SREK1  DHCR7  ANKRD36C  CEP126  LINC01573  COL16A1  PHLDB2  ZNF550  PHLDB1  RNMT  ANKRD36  PLXND1  C6ORF132  HOTAIRM1  PAPSS2  AP3M2  DACH1  EPYC  RAB26  CDH2  CDH1  ZNF767P  RBBP4  PAXIP1-AS2  MISP  ZNF302  ACSS1  LDLR  PNISR  LINC01123  SEMA6B  ATP8B4  OSBPL7  F2R  IGFBP6  ATP8B3  BTN2A2  ATP8B1  TNK2  TTC21A  FRMD4B  CTC-338M12.4  RCAN1  MAFF  PITPNM2  TRAF5  KIFC2  MIR1204  MALAT1  STAG3L5P-PVRIG2P-PILRB  MAFK  NACAD  ZNF775  MAP3K12  RAD9A  CACNB1  ANKRD33B  CUL9  CALCOCO1  CLSTN3  INSIG2  ANKRD12  INSIG1  GLUD1P3  PHF21A  RLF  ZFP62  KRBA1  CDC14B  RHOBTB3  SALL2  SALL4  HOXA-AS2  C1QTNF6  HOXA-AS3  PLXNA3  ZNF766  CCDC39  KCNIP2  HMGCS1  LANCL1  DNMT3B  SPACA6P  CDC42BPG  C15ORF52  FAM219B  MARCH8  DDHD2  CDC42BPB  NSUN5P1  TPCN2  CUL4B  BCL9  CDK6  FRG1BP  BCL6  ESRP2  BCL3  SLC26A9  CEP162  NF1  EFHC1  LINC01537  ZNF514  SERINC4  ZNF512  SLC26A4  SLC26A6  NLGN2  LMBR1L  MRI1  UBR5  PTPN23  ADRA1D  CYR61  PTBP2  KLC4  METTL15  DENND3  ZNF507  RPL36A  CCDC33  PDDC1  LBR  AMH  WASF1  LINC00674  MCL1  CNGA1  ZNF740  LINC00472  MAGI1  LINC00473  IRX3  SMARCC2  LIMCH1  TRIQK  COL27A1  PLEKHG2  SPAG8  OXR1  PLEKHG3  EP400NL  CCDC57  SPAG4  ILF3-AS1  KLF5  KLF3  ZFP90  KLF9  KLF7  CTDSP2  SH3D21  ZNF738  CCDC64  ANG  CCDC65  HDAC5  HDAC6  PLEKHH3  LSP1  RPL32P3  SEMA3F  PRR5L  PIK3C2B  TTN  PIK3C2A  FKBP9P1  WDR90  CACNG8  LOC100130899  PSD3  SCRN2  AP1G2  NHS  BCO2  PLEKHH2  CACNG6  TUB  SS18L1  ACSL3  PRSS30P  KRT8  P3H2  TNRC6C-AS1  YOD1  MAPK8IP3  SEPP1  SRRM2  MTUS1  MALL  TRIM52-AS1  XRN1  TBKBP1  NEIL1  LUCAT1  TECPR1  IFITM2  IFITM3  SLC46A3  IFITM1  C1RL-AS1  TRANK1  FRG1HP  CBLB  AQP1  AMER1  C3ORF17  OLMALINC  NIPAL1  TBL1X  APBB3  MEST  SRPK3  HLA-L  TMEM184A  LINC01089  STK11IP  ARRDC3  ARRDC4  LOC152225  PARP16  TRERF1  PARP14  SNHG4  INPP4B  CBARP  RPS27  FCHSD1  CCDC88A  STAG2  PWAR5  PARP10  TNNT1  CYP2S1  RBAK  PDE1B  TXK  CREBRF  DCUN1D2  CTDSPL2  PPL  IFIT2  FRG1JP  IFIT1  EPM2AIP1  CMYA5  INPP5E  PDK4  ATP6V0A4  TNRC18  RALGPS1  FAM65C  SOX9  FAM65B  SOX7  PDK1  SYNPO  TNS4  TNS3  ARGLU1  RALGPS2  SOX6  TNS2  EDN1  TSC22D2  BTN3A2  SLC35E2  BTN3A1  GAS8-AS1  LINC00174  LIF  BTN3A3  MLF1  RBM12B  CLDN11  P4HA1  CEACAM1  KRT19  KRT18  MIR17HG  VWA1  ALDOC  AKR1B10  CC2D1A  SGK1  TLR3  TOP2B  C2CD5  ZCCHC11  SLC16A1-AS1  MORF4L2-AS1  ZMYM3  MAP4  PRX  GSE1  ANKRD1  STRA6  KIAA1804  ASIC1  ASIC2  TM4SF18  FAM46A  THOC2  SORBS3  LIMA1  ATAT1  PDP1  PRPF39  PDXDC2P  B3GNT7  DLC1  TSSK4  LY6G5B  ZNF354B  TSSK3  SLC22A23  DCP2  TRIP10  ROBO3  ROBO4  GBE1  MRPS31P5  TRAK1  AMN1  EPB41L4B  DZANK1  RPARP-AS1  GABRE  RAB8B  KLHDC3  CA11  SZT2  TGFB2  VWDE  ZFC3H1  ARHGAP29  LIFR  LMO7  TSC1  TUG1  TMEM135  ARHGAP33  TBL1XR1  FAP  FAM161A  TMEM139  TGFBI  RELL2  MYADM  BRWD1  SCG2  BRWD3  CHRNB1  ZSCAN2  ZCCHC8  ARID4A  ZFP36L1  CCBL1  FBLN2  DNAJC27  EFNA5  CMIP  DOCK11  PRSS53  ARHGAP42  LOC220729  TMEM129  DNAJC22  NLRP1  KNTC1  PDE8A  PPP4R1L  REPS2  SMAD3  POSTN  USP49  USP43  HMHA1  GLTSCR1  UPK3B  AZIN2  ACSF2  C20ORF96  ARHGAP12  EFNA1  DLG1  EFNA3  DLG3  LOX  CAPRIN2  MATN2  CLASRP  ACBD4  SERTAD4-AS1  EIF3J-AS1  DOCK6  AKNA  ISYNA1  MAGEB17  DOCK9  SLC2A3  ARHGAP18  LTBP3  LTBP4  AGER  NR4A1  SEPSECS  PLCXD1  CREBZF  TMEM100  SYDE1  EME2  STARD4-AS1  SVEP1  USP53  KLRC2  ANKZF1  USP25  SETD1A  USP21  PACERR  SETD1B  CBX2  NFKBIZ  ARAP3  ARAP1  ANXA2R  DNM1  IQCK  IQCH  TGFBR2  MEIS2  FER  RBL2  LPIN1  JMY  PIKFYVE  FAM84B  RBMS2  LPIN3  LUC7L3  RBMS3  KDM5B  HAPLN1  B4GALT1  USP32  C2ORF48  LOC101929705  USP35  NR2C1  NR2C2  DBP  LRRTM4  POM121L9P  NRIP2  MAN2C1  BDKRB2  PDE4B  FGA  LONRF1  KDM6B  FGB  PTGIS  SPEN  FGG  LMNTD2  LCA5  KCTD3  ARID1A  C14ORF132  KCTD7  CLDN7  FAM167A  ALDH1A1  PPP2R2D  DDIT4  CLK2P1  KIAA0907  PDE5A  SPRY1  PXYLP1  HCN3  THRA  C2ORF68  PPP2R2B  SLC4A3  KANSL1  THBS3  CCDC28B  SENP7  RMND5A  SEPT8  PCED1A  RPS6KA3  PCED1B  POLG  POLI | PDGFRB  CA11  NIPSNAP3A  RARRES3  PLEKHA1  TNRC6C-AS1  KLHL24  LIFR  COL1A2  ST5  RBP1  RFX5  SIDT2  TGFBI  YPEL2  CYB5RL  MXRA5  ROR1  TRABD2B  JRK  IFITM1  SCNN1A  CELSR2  FOXO4  DEFB1  TFPI  GNG2  ATOH8  RAB26  CDH2  ABI3BP  PLCE1  SERPINF1  MUC6  GBP4  FZD2  SEMA6B  TSPYL2  TNFSF18  F2R  FZD5  ARRDC4  PLK2  LOC642852  IFFO2  INPP4B  EFNA1  BCAM  PARP10  HBP1  SERTAD4-AS1  NOX3  FTO  CALCOCO1  CLSTN3  CFI  STMN3  ARHGAP18  LTBP3  PLAT  ATXN7L3B  IFIT2  IFIT1  CDC14B  TMEM100  CALHM3  C1QTNF6  PDK4  TNFSF10  SVEP1  BMF  TNS3  TNS2  PSMD5-AS1  BTN3A2  BTN3A1  ANXA4  BTN3A3  CP  GPRC5A  BMP4  MEIS2  RBL2  CEACAM1  VWA1  FAM84B  CCNG1  SLC26A4  TMEM106B  DNAH2  CHD6  TRIM2  IQGAP2  SERPINA5  ADAMTS10  ZMYM3  LRRTM4  ASIC2  DACT2  TM4SF18  MAGI1  PTGIS  SLC2A12  TP53INP1  RNASE4  C14ORF132  ZNF33B  AGBL5  TXNIP  ANG  LSP1  ITGB4  QPRT  KIAA1551  SEMA3F  THBS3  PBXIP1  PTGS1  RMND5A  SERTAD4  ING4  ZNF608  DMBT1  TCTN2  METTL7A  THSD7A  AGR2  BCO2 |
