## Supplemental Table 3 for "MondoA mediates transcriptional coordination between the MYC network and the integrated stress response in pancreatic ductal adenocarcinoma"

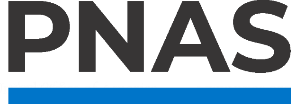

**Supporting Information for**

MondoA mediates transcriptional coordination between the MYC network and the integrated stress response in pancreatic ductal adenocarcinoma.

Paste the full author list here Erin L. Ramsey^1^, Stephanie Dobersch^2^, Brian Freie^1^, Nan Hyung Hong^1^, Xiaoying Wu^1^, Sita Kugel^2^, Robert N. Eisenman^1,^*, Patrick A. Carroll^1,^*

^1^Basic Sciences Division, Fred Hutchinson Cancer Center, Seattle, USA

^2^Human Biology Division, Fred Hutchinson Cancer Center, Seattle, USA

*Corresponding authors Robert N. Eisenman and Patrick A. Carroll

**This PDF file includes:**

Supplemental Table 3

**Supplemental Table 3: Kaplan-Meier survival curve gene lists from TCGA.** BOLD: denotes individually prognostic

| **MYC:AMP Up and siMondoA Down** | **MYC:AMP Up and SBI-477 Down** | **MYC:AMP Down and siMondoA Up** | **MYC:AMP Down and SBI-477 Up** |
| --- | --- | --- | --- |
| EGLN3; JUP; **AHNAK2**; IGFBP3; MBOAT2; PPA1; SORD; NUTF2; **YWHAZ**; SUMF1; MRPS6; APRT; HPDL; **GPRC5A**; PTMA; KRT19; NET1; EBPL; VDAC1; EIF3E; MAD2L1; **NUP37**; **COL17A1**; SLC44A2; ITGB4; TXN; SCNN1A; AK4; **UNG**; **LDHA**; EXOSC5; FAM162A; SUMO3; **PERP**; **AP1S3**; CLIC3; **ZNF488**; **EIF2A** | ANXA3; ITGB4; PVT1; SCNN1A; ASAP2; C6ORF132; **TRERF1**; TM4SF1; **GPRC5A**; MALL; **TBL1XR1**; KRT19; EFNA3; **TNNT1**; EPB41L1; SYT12; ESRP2; CDH1; FLNB; **ECT2**; TNS4; **ZNF488**; **NFE2L3**; TRIP10 | TAPT1; EGR3; **AKNA**; XBP1; ZNF592; KSR1; GPS2; SECISBP2; ZBTB21; RNFT1; RPGR; FAM193A; ENPP3; **PI4KB**; **CAMTA2**; DENND5B; ABL1; KDM6B; **TNRC6C** | NRROS; **SYNM**; TTLL11; GPS2; **NCOA5**; RDH8; LANCL2; **PITPNA**; TCEAL1; TIMM22; FGFR1; ZBED1; QRICH2 |
