## Supplemental Tables 4-6 for "MondoA mediates transcriptional coordination between the MYC network and the integrated stress response in pancreatic ductal adenocarcinoma"

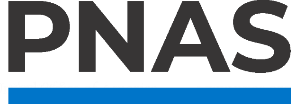

**Supporting Information for**

MondoA mediates transcriptional coordination between the MYC network and the integrated stress response in pancreatic ductal adenocarcinoma.

Paste the full author list here Erin L. Ramsey^1^, Stephanie Dobersch^2^, Brian Freie^1^, Nan Hyung Hong^1^, Xiaoying Wu^1^, Sita Kugel^2^, Robert N. Eisenman^1,^*, Patrick A. Carroll^1,^*

^1^Basic Sciences Division, Fred Hutchinson Cancer Center, Seattle, USA

^2^Human Biology Division, Fred Hutchinson Cancer Center, Seattle, USA

*Corresponding authors Robert N. Eisenman and Patrick A. Carroll

**This PDF file includes:**

Supplemental Tables 4 to 6

**Supplemental Table 4: siRNA Details.** All siRNAs from Qiagen. Control siRNA catalogue numbers provided, targeted gene mixes are FlexiTube Gene Solutions Cat. No. 1027416, gene ID and target transcripts provided.

| **Target** | **Cat. No.** | **Full Name** |
| --- | --- | --- |
| siControl | 1027281 | AllStars Neg. Control |
| siDeath | 1027299 | AllStars Hs Cell Death Control |
| **Target** | **Gene Globe ID** | **Target Transcripts** |
| siARRDC4 | GS91947 | Hs_ARRDC4_2, Hs_ARRDC4_6, Hs_ARRDC4_8, Hs_ARRDC4_9 |
| siATF4 | GS468 | Hs_ATF4_5, Hs_ATF4_8, Hs_ATF4_9, Hs_ATF4_10 |
| siFTO | GS79068 | Hs_FTO_5, Hs_FTO_6, Hs_FTO_7, Hs_FTO_8 |
| siMLX | GS6945 | Hs_MLX_1, Hs_TCFL4_2, Hs_TCFL4_3, Hs_TCFL4_4 |
| siMLXIP | GS22877 | Hs_MLXIP_1, Hs_MLXIP_2, Hs_MLXIP_3, Hs_MLXIP_4 |
| siMLXIPL | GS51085 | Hs_MLXIPL_1, Hs_MLXIPL_2, Hs_MLXIPL_3, Hs_WBSCR14_3 |
| siMNT | GS4335 | Hs_MNT_5, Hs_MNT_6, Hs_MNT_7, Hs_MNT_8 |
| siMYC | GS4609 | Hs_MYC_5, Hs_MYC_7, Hs_MYC_8, Hs_MYC_9 |
| siPPP1R15A | GS23645 | Hs_PPP1R15A_5, Hs_PPP1R15A_6, Hs_PPP1R15A_7, Hs_PPP1R15A_8 |
| siTXNIP | GS10628 | Hs_TXNIP_5, Hs_TXNIP_6, Hs_TXNIP_7, Hs_TXNIP_8 |

**Supplemental Table 5: Antibody Details.**

| **Application** | **Target** | **Species** | **Company** | **Cat. No.** | **Concentration** |
| --- | --- | --- | --- | --- | --- |
| Western Blot | Mouse Secondary-HRP | Horse | Cell Signaling | 7076S | 1:2,500 |
| Western Blot | Rabbit Secondary-HRP | Goat | Cell Signaling | 7074S | 1:2,500 |
| Western Blot | ATF4 | Rabbit | Cell Signaling | 11815 | 1:1,000 |
| Western Blot | ATF6 | Rabbit | Cell Signaling | 65880 | 1:1,000 |
| Western Blot | CHOP | Mouse | Cell Signaling | 2895S | 1:500 |
| Western Blot | eIF2α | Rabbit | Cell Signaling | 5324 | 1:3,000 |
| Western Blot | FTO | Mouse | ProteinTech | 68111-1-Ig | 1:10,000 |
| Western Blot | FTO | Rabbit | ProteinTech | 27226-1-AP | 1:2,000 |
| Western Blot | GADD34/PPP1R15A | Rabbit | ProteinTech | 10449-1-AP | 1:1,000 |
| Western Blot | MondoA/MLXIP | Rabbit | ProteinTech | 13614-1-AP | 1:2,000 |
| Western Blot | MYC | Rabbit | Cell Signaling | 13987 | 1:1,000 |
| Western Blot | P- eIF2α | Rabbit | Cell Signaling | 3398 | 1:1,000 |
| Western Blot | Puromycin | Mouse | Millipore | MABE343 | 1:1,000 |
| Western Blot | TXNIP | Mouse | MBL International | K0205-3 | 1:2,000 |
| Western Blot | XBP-1s | Rabbit | Cell Signaling | 12782 | 1:1,000 |
| Western Blot | βActin | Mouse | Sigma-Aldrich | A5441 | 1:5,000 |
| Western Blot | γH2AX | Rabbit | Abcam | Ab2893 | 1:2,500 |
| Western Blot | γTubulin | Mouse | Sigma-Aldrich | T5326 | 1:10,000 |
| Immunofluorescence | Mouse Secondary- AlexaFluor 448 | Goat | Invitrogen | A11001 | 1:400 |
| Immunofluorescence | Rabbit Secondary- AlexaFluor 568 | Goat | Invitrogen | A11011 | 1:400 |
| Immunofluorescence | m^6^A | Mouse | ProteinTech | 68055-1-Ig | 1:1,000 |
| Immunofluorescence | m^6^A | Rabbit | Novus Biologicals | NBP3-05657 | 1:500 |
| Flow Cytometry | Rabbit Secondary- AlexaFluor 750 | Goat | Invitrogen | A21039 | 1:200 |
| Flow Cytometry | m^6^A | Rabbit | Novus Biologicals | NBP3-05657 | 1:200 |

**Supplemental Table 6: Characterization of Patient Derived Organoids.**

| **PPTO** | **Biobank ID** | **Age** | **Sex** | **STR** | **Mycoplasma** | **Classification** |
| --- | --- | --- | --- | --- | --- | --- |
| 174 | 107407 | 67 | Female | Match patient | Negative | Basal |
| 69 | 91168 | 50 | Male | Match patient | Negative | Classical |
| 74 | 91416 | 62 | Female | Match patient | Negative | Basal |
| 97 | 93429 | 68 | Female | Match patient | Negative | Classical |
| 165 | 106808 | 52 | Male | Match patient | Negative | Classical |
| 105 | 95139 | 53 | Male | Match patient | Negative | Classical |
| 154 | 104760 | 59 | Male | Match patient | Negative | Basal |
